# Detecting selection from linked sites using an F-model

**DOI:** 10.1101/737916

**Authors:** Marco Galimberti, Christoph Leuenberger, Beat Wolf, Sándor Miklós Szilágyi, Matthieu Foll, Daniel Wegmann

**Author notes:** Department of Biology, University of Fribourg, Chemin du Musée 10, 1200 Fribourg, Switzerland. Where authors are identified as personnel of the International Agency for Research on Cancer / World Health Organization, the authors alone are responsible for the views expressed in this article and they do not necessarily represent the decisions, policy or views of the International Agency for Research on Cancer / World Health Organization.

## Abstract

Allele frequencies vary across populations and loci, even in the presence of migration. While most differences may be due to genetic drift, divergent selection will further increase differentiation at some loci. Identifying those is key in studying local adaptation, but remains statistically challenging. A particularly elegant way to describe allele frequency differences among populations connected by migration is the *F*-model, which measures differences in allele frequencies by population specific *F*_ST_ coefficients. This model readily accounts for multiple evolutionary forces by partitioning *F*_ST_ coefficients into locus and population specific components reflecting selection and drift, respectively. Here we present an extension of this model to linked loci by means of a hidden Markov model (HMM) that characterizes the effect of selection on linked markers through correlations in the locus specific component along the genome. Using extensive simulations we show that our method has up to two-fold the statistical power of previous implementations that assume sites to be independent. We finally evidence selection in the human genome by applying our method to data from the Human Genome Diversity Project (HGDP).

---

Migration is a major evolutionary force homogenizing evolutionary trajectories of populations by promoting the exchange of genetic material. At some loci, however, the influx of new genetic material may be modulated by selection. In case of strong local adaptation, for instance, migrants may carry maladapted alleles that are selected against. Identifying loci that contribute to local adaptation is of major interests in evolutionary biology because these loci are thought to constitute the first step towards ecological speciation (e.g. Wu 2001; Feder *et al*. 2012) and allow us to understand the role of selection in shaping phenotypic differences between populations and species (e.g. Bonin *et al*. 2006; Fournier-Level *et al*. 2011).

A simple yet flexible and useful approach to identify loci contributing to local adaptation is to scan the genome using statistics that quantify divergence between populations. One frequently used such statistics is *F*_ST_ that measures population differentiation, and loci with much elevated *F*_ST_ have been reported for many population comparisons (e.g. Jones *et al*. 2012; Andrew and Rieseberg 2013; Stölting *et al*. 2013). While other statistics measuring absolute divergence (Cruickshank and Hahn 2014) or assessing incongruence between a population tree and the genealogy at a locus (Durand *et al*. 2011; Peter 2016) may be more suited in some situations, genome scans suffer from two inherent limitations. First, multiple evolutionary scenarios may explain the deviations in those statistics, making interpretation difficult (Cruickshank and Hahn 2014; Eriksson and Manica 2012). Second, the definition of outliers is arbitrary, allowing for the detection of candidate loci only. Indeed, loci also vary in their divergence between populations that were never subjected to selection, but outlier approaches would still be identifying outliers.

Multiple methods have thus been developed that explicitly incorporate the stochastic effects of genetic drift. A first important step to improve the reliability of outlier scans was the proposal to compare observed values of such statistics against the distribution expected under a null model. Among the first, Beaumont and Nichols (1996) proposed to obtain the distribution of *F*_ST_ through simulations performed under an island model. While the idea to evidence selection by comparing *F*_ST_ to its expectations is far from new (e.g. Lewontin and Krakauer 1973), the difficulty to properly parameterize the null model was quickly realized (e.g. Nei and Maruyama 1975). The success of the method by Beaumont and Nichols (1996) relies on tailoring the parameters of the underlying island model to match the observed heterozygosity at each locus, an approach that is also easily extended to structured island models (Excoffier *et al*. 2009).

A more formal approach is given by means of the *F*-model (Balding 2003; Falush *et al*. 2003; Gaggiotti and Foll 2010; Rannala and Hartigan 1996), under which allele frequencies are measured by locus and population specific 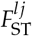 coefficients that reflect the amount of drift that occurred in population *j* at locus *l* since its divergence from a common ancestral population. In the case of bi-allelic loci, the current frequencies 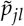 are then given by a beta distribution (Beaumont and Balding 2004)

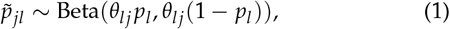

where *p_l_* are the frequencies in the ancestral population and *θ_lj_* is given by

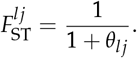

It is straightforward to extend this model to account for different evolutionary forces that effect the degree of genetic differentiation. Beaumont and Balding (2004), for instance, proposed to partition the effects of genetic drift and selection into locus specific and a population specific components *α_l_* and *β_j_*, as well as a locus-by-population specific error term *γ_ij_*:

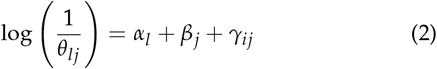

Loci with *α_l_* ≠ 0 are interpreted to be targets of either balancing (*α_l_* < 0) or divergent (*α_l_* > 0) selection (Beaumont and Balding 2004). Targets of selection may then be identified by contrasting models with *α_l_* = 0 or *α_l_* ≠ 0 for each locus *l*, either through Bayesian variable selection (Riebler *et al*. 2008) or via reversible-jump MCMC, as is done in the popular software `BayeScan` (Foll and Gaggiotti 2008).

A common problem of this and many other genome-scan methods is the assumption of independence among loci, which is easily violated when working with genomic data. By evaluating information from multiple linked loci jointly, however, the statistical power to detect outlier regions is likely increased considerably. Indeed, even a weak signal of divergence may become detectable if it is shared among multiple loci. Similarly, false positivs may be avoided as their signals is unlikely shared with linked loci.

Unfortunately, fully accounting for linkage is often statistically challenging as well as computationally very costly. One solutions is to split the problem by first inferring haplotypes for each sample, and then performing selection scans on the haplotype structure. The extended haplotype homozygosity (EHH) and its derived statistics (Sabeti *et al*. 2002; Voight *et al*. 2006; Sabeti *et al*. 2007; Tang *et al*. 2007), for instance, identify shared haplotypes of exceptional length. More recently, Fariello *et al*. (2013) introduced methods that identify haplotype clusters with particularly large frequency differences between populations and showed that using haplotypes rather than single markers increases power substantially.

An alternative solution is to model linkage through the auto-correlation of hierarchical parameters along the genome, which does not require knowledge on the underlying haplotype structure. Boitard *et al*. (2009) and Kern and Haussler (2010), for instance, proposed a genome-scan method in which each locus was classified as selected or neutral, and then used a Hidden Markov Model (HMM) to account for the fact that linked loci likely belonged to the same class, while ignoring auto-correlation in the genetic data itself.

Here we build on this idea to develop a genome-scan method based on the *F*-model. While an HMM implementation of the *F*-model was previously proposed to deal with linked sites when inferring admixture proportions (Falush *et al*. 2003), we use it here to characterize auto-correlations in the strength of selection *α_l_* among linked markers. As we show using both simulations and an application to human data, aggregating information across loci results in up to two-fold power at the same false-discovery rate.

## Methods

### A Model for Genetic Differentiation and Observations

We assume the classic *F*-model in which *J* populations diverged from a common ancestral population. Since divergence, each population experienced genetic drift at a different rate. We quantify this drift of population *j* = 1, …, *J* at locus *l* = 1, …, *L* by *θ_jl_*. We further assume each locus to be bi-allelic with ancestral frequencies *p_l_*, in which case the current frequencies 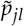 are given by a beta distribution (Beaumont and Balding 2004), as shown in (2). We thus have

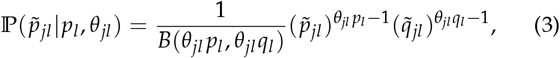

where *q_l_* = 1 – *p_l_*, 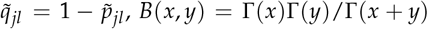 and Γ(·) is the gamma function.

Let *n_jl_* denote the allele counts in a sample of *N_jl_* haplotypes from population *j* at locus *l*, which is given by a binomial distribution

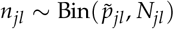

and hence

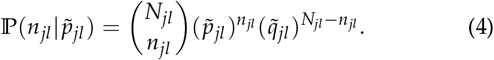

Equations (3) and (4) combine to a beta-binomial distribution

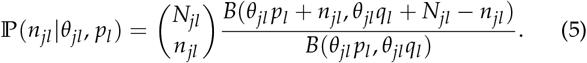

### Model of selection

In the absence of selection, all loci are assumed to experience the same amount of population specific drift. We thus decompose *θ_jl_* into a population-specific component *β_j_* shared by all loci, and a locus-specific component *α_l_* shared by all populations:

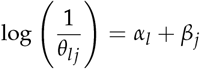

Note that we adopt here the formulation of Foll and Gaggiotti (2008) and omit the error term *γ_ij_* of the original model of Beaumont and Balding (2004) shown in (2), as there is generally not enough information to estimate these parameters from the data (Beaumont and Balding 2004).

To account for auto-correlation among the locus-specific component, we propose to discretize *α_l_* = *α*(*s_l_*), where *s_l_* = −*s*_max_, −*s*_max_ + 1, …, *s*_max_ are the states of a ladder-type Markov model with *m* = 2*s*_max_ + 1 states such that

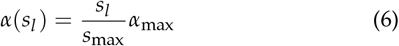

for some positive parameters *α*_max_. The transition matrix of this Markov model shall be a finite-state birth-and-death process

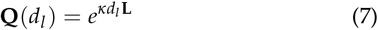

with elements 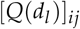 denoting the probabilities to go from state *i* at locus *l* – 1 to state *j* at locus *l* given the strength of auto-correlation measured by the positive scaling parameter *κ* and the known distance *d_l_* between these loci, either in physical or in recombination space. Here, **L** is the *m* × *m* generating matrix

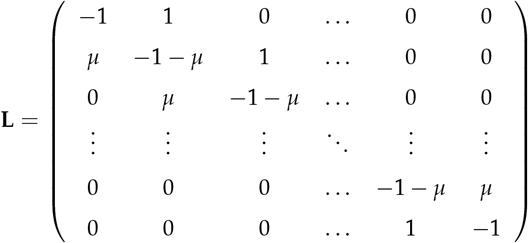

where the middle row at position *s*_max_ +1 reflects neutrality and is given by the element

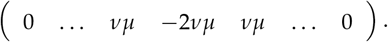

As exemplified in Figure 1, the two parameters *μ* and *v* control the distribution of sites under selection in the genome with large *v* affecting the number of selected regions and *μ* their extent and selection strength, with higher values leading to more sites under selection. The stationary distribution of this Markov chain is given by

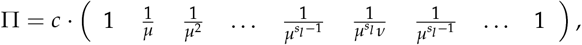

with

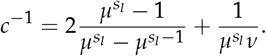

**Figure 1.**
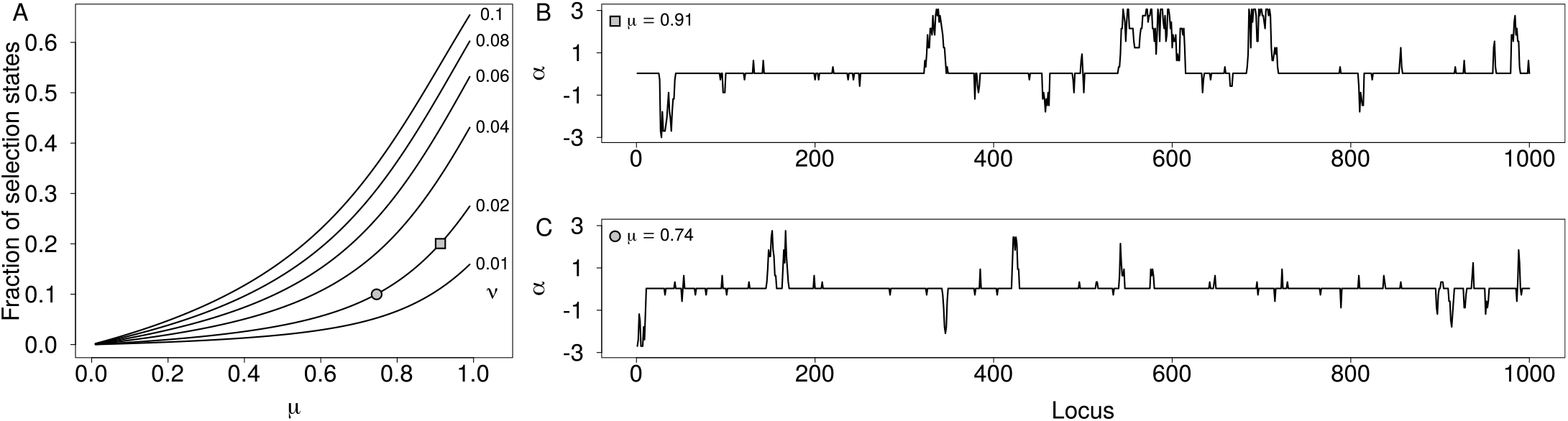
(A) expected proportion of neutral sites as a function of rates *μ* and *v*. (B and C) Example paths of *α_l_* along 1,000 loci simulated at a distance of *d_l_* = 100 with *s*_max_ = 10 positive and negative states up to *α*_max_ = 3.0. Autocorrelation among loci was simulated with log(*κ*) = −3.0, *v* = 0.02 and *μ* = 0.91 (B, square) or *μ* = 0.74 (C, circle), respectively. The two cases correspond to an expected proportion of 20% and 10% of the genome under selection, as marked in A.

Note that as *κ* → ∞, our model approaches that of Foll and Gaggiotti (2008) implemented in `BayeScan` but with discretized *α_l_*.

### Hierarchical Island Models

Hierarchical island models, first introduced by Slatkin and Voelm (1991), address the fact the divergence might vary among groups of populations. They were previously used to infer divergent selection, both using a simulation approach (Excoffier *et al*. 2009) as well as in the case of *F*-models (Foll *et al*. 2014). Here we describe how our model is readily extended to additional hierarchies.

Consider *G* groups each subdivided into *J_g_* populations with population specific allele frequencies 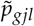 that derive from group-specific frequencies *p_gl_* as described above with group-specific parameters *μ_g_*, *v_g_* and *κ_g_*. Analogously, we now assume group-specific frequencies to have diverged from a global ancestral frequency *P_l_* according to locus-specific and group-specific parameters Θ_*gl*_. Specifically,

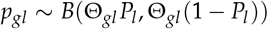

such that

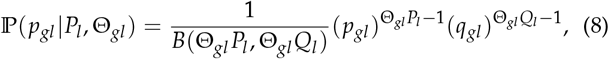

where 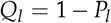 and *q_gl_* = 1 – *p_gl_*. The parameter Θ_*gl*_ is given by

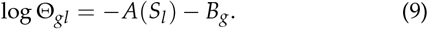

As above, *B_g_* quantifies group specific drift, *S_l_* = −*s*_max_, −*s*_max_ + 1, …, *s*_max_ are the states of a Markov model with *m* states and transition matrix **Q**_*l*_ = *e*^*κd_l_***L**^ with parameters *μ* and *ν*, a positive scaling parameter *κ* and *A*(*S_l_*) and *A*_max_ defined as in (6). Hence, we assume independent HMM models of the exact same structure at all levels of the hierarchy, as outlined in Figure 2.

**Figure 2.**
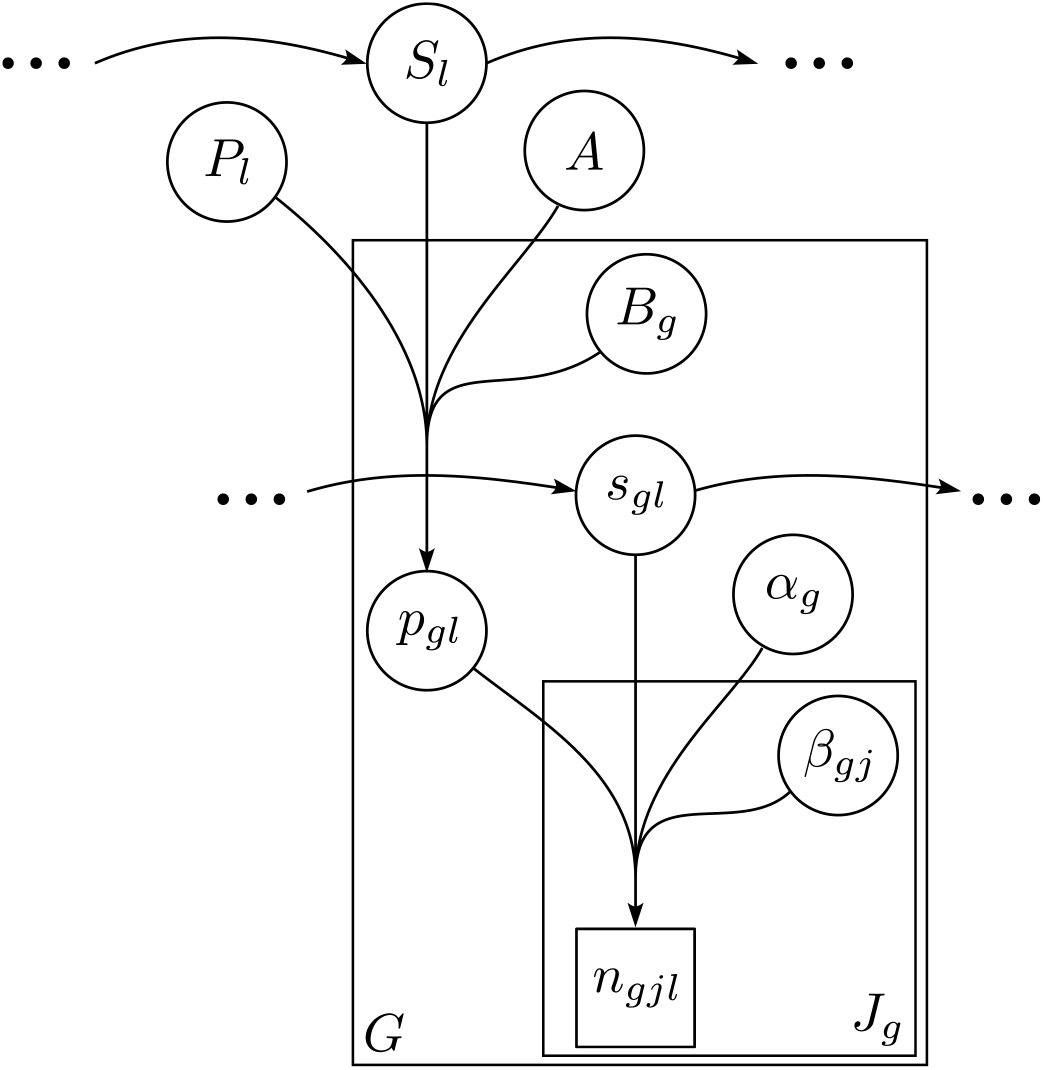
A directed acyclic graph (DAG) of the proposed hierarchical model.

### Inference

We implemented a Bayesian inference scheme for the proposed model using a Markov chain Monte Carlo (MCMC) approach using Metropolis–Hastings updates, as detailed in the Supplementary Material. As priors, we used

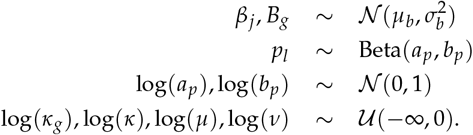

Following Beaumont and Balding (2004), we used *μ_b_* = −2 and 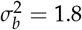 throughout. We further set *a_p_* = *b_p_* = 1.

To identify candidate regions under selection, we used our MCMC samples to determine the false-discovery rates

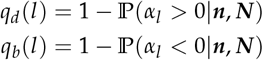

for divergent and balancing selection, respectively, where ***n*** = {*n*_11_, …, *n_JL_*} and ***N*** = {*N*_11_, …, *N_JL_*} denote the full data.

### Implementation

We implemented the proposed Bayesian inference scheme in the easy-to-use `C++` program `Flink`.

Given the heavy computational burden of the proposed model, we introduce several approximations. Most importantly, we group the distances *d_l_* into *E* + 1 ensembles such that *e_l_* = ⌈log_2_ *d_l_*⌉, *e_l_* = 0, …, *E* and use the same transition matrix **Q**(2^*e*^) for all loci in ensemble *e*. We then calculate **Q**(1) for the first ensemble using the computationally cheap yet accurate approximation

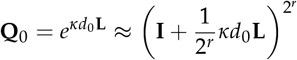

with *r* = log_2_(*D*/3) + 10 where *D* = 2*s*_max_ + 1 is the dimensionality of the transition matrix (Ferrer-Admetlla *et al*. 2016). The transition matrices of all other ensembles can then be obtained through the recursion **Q**(*e*) = **Q**(*e* – 1)^2^. (See Supplementary Information for other details regarding the implementation).

### Data availability

The authors affirm that all data necessary for confirming the conclusions of the article are present within the article or available from repositories as indicated. The source-code of `Flink` is available through the git repository https://bitbucket.org/wegmannlab/flink, along with detailed information on its usage. Additional scripts used to conduct simulations are found https://doi.org/10.5281/zenodo.3949763.

## Comparison with BayeScan

### Simulation parameters

To quantify the benefits of accounting for auto-correlation in the locus specific components *α_l_* among linked loci, we used simulations to compare the power to identify loci under selection of our method implemented in `Flink` against the method implemented in `BayeScan` (Foll and Gaggiotti 2008). All simulations were conducted under the model laid out above for a single group using routines available in `Flink` and with parameter settings similar to those used in (Foll and Gaggiotti 2008). Specifically, we focused on a reference simulation in which we sampled *N* = 50 haplotypes from *J* = 10 populations with *β_j_* chosen such that 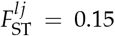 in the neutral case (*α_l_* = 0). We then varied the number of populations *J*, the sample size *N*, 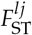 or the strength of auto-correlation *κ* individually, while keeping all other parameters constant (Table 1). Following Foll and Gaggiotti (2008), we simulated according to our model all *p_l_* ~ Beta(0.7,0.7) and 20% of sites under selection by setting *μ* = 0.91 and *v* = 0.02. We further set *s*_max_ = 10 (resulting in *m* = 21 states) and *α*_max_*g*__=3 for all simulations. We simulated 10^3^ loci for each of 10 chromosomes, with a distance of 100 positions between adjacent sites. We further added a case without linkage (i.e. *κ* → ∞) by simulating each locus on its own chromosome.

**Table 1.** Parameters used in simulations

| Name | J | $F_{ST}$ | N | $\log(\kappa)$ |
| --- | --- | --- | --- | --- |
| Reference | 10 | 0.15 | 50 | -3 |
| Pop-2 | 2 | 0.15 | 50 | -3 |
| Pop-5 | 5 | 0.15 | 50 | -3 |
| Pop-20 | 20 | 0.15 | 50 | -3 |
| Pop-50 | 50 | 0.15 | 50 | -3 |
| $F_{ST}$-0.01 | 10 | 0.01 | 50 | -3 |
| $F_{ST}$-0.05 | 10 | 0.05 | 50 | -3 |
| $F_{ST}$-0.1 | 10 | 0.1 | 50 | -3 |
| $F_{ST}$-0.25 | 10 | 0.25 | 50 | -3 |
| Haplo-10 | 10 | 0.15 | 10 | -3 |
| Haplo-20 | 10 | 0.15 | 20 | -3 |
| Haplo-100 | 10 | 0.15 | 100 | -3 |
| Haplo-200 | 10 | 0.15 | 200 | -3 |
| $\log \kappa$-1 | 10 | 0.15 | 50 | -1 |
| $\log \kappa$-5 | 10 | 0.15 | 50 | -5 |
| $\log \kappa$-7 | 10 | 0.15 | 50 | -7 |
| $\log \kappa$-9 | 10 | 0.15 | 50 | -9 |

To infer parameters with `Flink`, we set *s*_max_ and *α*_max_ to the true values and ran the MCMC for 7 · 10^5^ iterations, of which we discarded the first 2 · 10^5^ as burnin. During the chain, we recorded all parameter values every 100 iterations as posterior samples. To infer parameters with `Bayescan`, we used version 2.1 and set the prior odds for the neutral model to 50, which we found to result in the same power as `Flink` in the reference simulation (see below) and in the absence of linkage (*κ* → ∞). We identified loci under selection at a False-Discovery-Rate (FDR) threshold of 5% for both methods.

### Power of inference

We first evaluated the power of `Flink` in inferring the hierarchical parameters *β_j_*, *v*, *μ* and *κ*. As shown through the distributions of posterior means across all simulations, these estimates were very accurate and unbiased, regardless of the parameter values used in the simulations (Figure 3). This suggests that the power to identify selected loci is not limited by the number of loci we used to infer hirearchical parameters.

**Figure 3.**
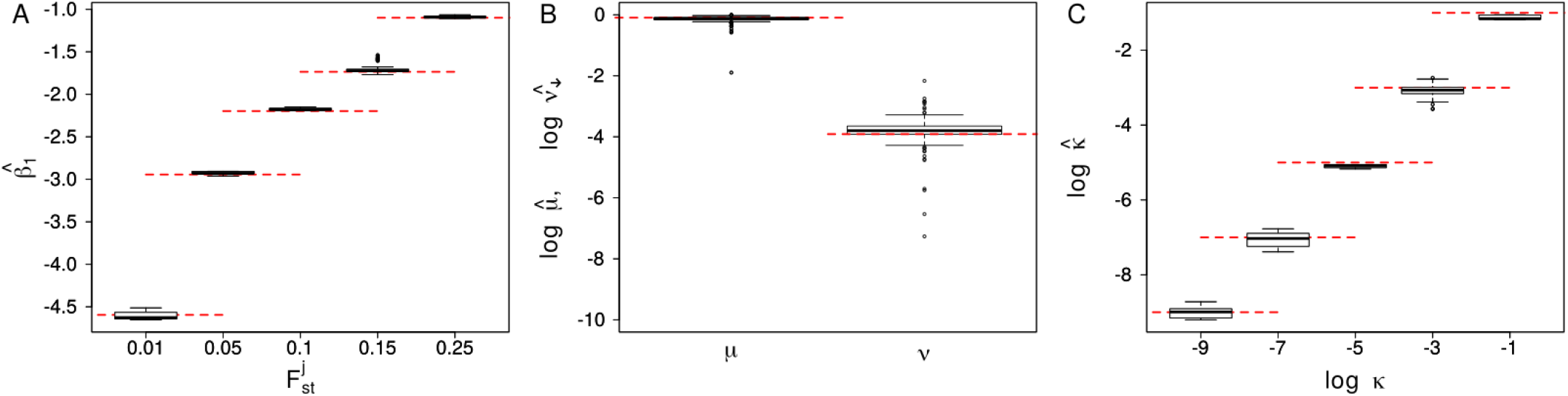
Boxplot of the parameters *β*_1_ (left), *v* and *μ* (center) and log(*κ*) (right). The values are obtained from the mean of the posterior distributions obtained using `Flink` on the 10 simulations run for each of the set of parameters reported in Table 1. The red dotted lines show the true values of the respective parameters.

We next studied the impact of the sample size and the strength of population differentiation on power. In line with findings reported by Foll and Gaggiotti (2008), power generally increased with 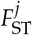, the number of sampled haplotypes and the number of sampled populations (Figure 4A-C). Importantly, larger sample sizes or stronger differentiation was particularly relevant for detecting loci under balancing selection, for which the power was generally lower and virtually zero at low differentiation 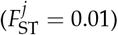 or if only few populations were sampled (*J* = 2).

**Figure 4.**
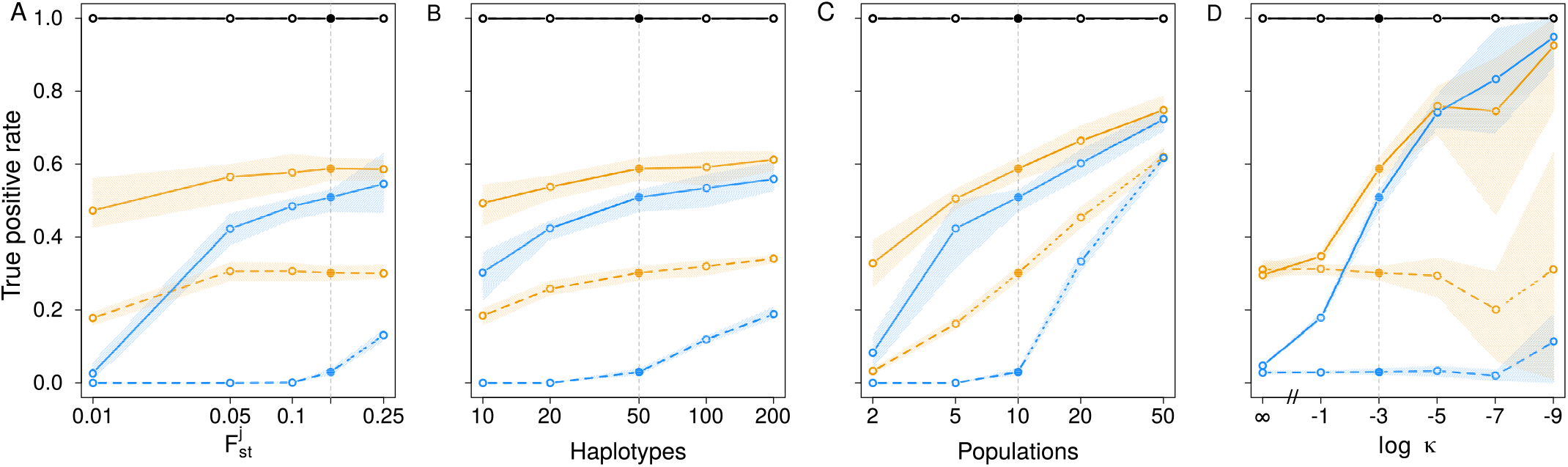
The true positive rate in classifying loci as neutral (black) or under divergent (orange) or balancing selection (blue) as a function of the *F*_ST_ between populations (A), the number of haplotypes *N* (B), the number of populations *J* (C) and the strength of auto-correlation *κ* (D). Lines indicate the mean and range of true positive rates obtained with `Flink` (solid) and `BayeScan` (dashed) across 10 replicate simulations. Filled dots and the vertical gray line indicate the reference simulation shown in each plot.

We finally compared the power of `Flink` to that of `BayeScan` on the same set of simulations. As shown in Figure 4, `Flink` had a higher power at the same FDR across all simulations, and often considerably so, unless the number of populations sampled was large. If *J* = 10 populations were sampled, for instance, the power of `Flink` was about 0.2 higher for loci under divergent selection, and even up to 0.4 higher for those under balancing selection (Figure 4A,B).

Importantly, this increase in power described here is fully explained by `Flink` accounting for auto-correlation among the *α_l_* values as we chose the prior odds in `BayeScan` to result in the same power if the strength of auto-correlation vanishes (i.e. *κ* → ∞). Exploiting information from linked sites to identify divergent or balancing selection can thus strongly increase power, certainly if linkage extends to many loci. This is maybe best illustrated by the much higher power of `Flink` to identify loci under balancing selection at low differentiation (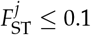, Figure 4A), in which case even many neutral loci are expected to show virtually no difference in allele frequency and only an aggregation of such loci can be interpreted as a reliable signal for selection (Foll and Gaggiotti 2008).

### Runtime

Thanks to careful optimization, there is little to no overhead of our implementation compared to that of `BayeScan`. On the reference simulation of 10^4^ loci from 10 populations, for instance, `Flink` took on average 130 minutes on a modern computer if calculations were spread over 4 CPU cores. On the same data, `BayeScan` took 361 minutes. However, we note that comparing the two implementations is difficult due to many settings that strongly impact run times such as the number of iterations or the use of pilot runs in `BayeScan`. Without pilot runs, the run time of `BayeScan` reduced to 182 minutes on average for the default number of iterations (10^5^ including burnin). In the same time, `Flink` runs for close to 10^6^ iterations, but also requires more to converge.

But since computation times scale linearly with the number of loci, they remain prohibitively slow for whole genome applications in a single run. However, they computations are easily spread across many computers by analyzing the genome in independent chunks such as for each chromosome or chromosome arm independently. This is justified because 1) linkage does not persist across chromosome boundaries and is usually also weak across the centromere and 2) because our simulations indicate that 10^4^ polymorphic loci were sufficient to estimate the hierarchical parameters accurately.

## Effect of model misspecification

The F-model makes the explicit assumption that the allele frequencies in a structured population can be characterized by a multinomial Dirichlet distribution. This distribution is appropriate for a wide range of demographic models, but not if some pairs of populations share a more recent ancestry than others (Beaumont and Balding 2004; Excoffier *et al*. 2009). Unsurprisingly, several previous studies found high false-positive rates when challenging `BayeScan` with models of isolation-by-distance (IBD), recent range expansions, recent admixture or asymmetric divergence (e.g. Lotterhos and Whitlock 2014; Luu *et al*. 2017). These high false-positive rates are partially mitigated by choosing higher prior odds (e.g. 50 as used here, Lotterhos and Whitlock 2014) or when using the hierarchical version of `BayeScan` (Foll *et al*. 2014), particularly in case of asymmetric divergence. In the case of a recent range expansion or recent admixture, however, the F-model is unlikely appropriate and other methods have been shown to outperform `BayeScan`, in particular `hapFLK` (Fariello *et al*. 2013) and `pcadapt` (Luu *et al*. 2017).

Here we investigated how the sensitivity of the linkage-aware implementation of an F-model in `Flink` is affected by such model misspecifications. We focused on the case of a recent range expansion as this model is difficult to accommodate even with a hierarchical F-model. Specifically, we used `quantiNemo` (Neuenschwander *et al*. 2018) to simulate genomic data from 11 populations with carrying capacity 1,000 each that form a one-dimensional stepping-stone model. Initially, only the left-most population contained individuals that then colonized the remaining populations through symmetric dispersal between neighboring populations at rate 0.1 and with a population growth rate of 0.1. After 1,000 generations, 20 diploid individuals were sampled from each population. We simulated 10 independent chromosomes of 10,000 neutral loci each with initial allele frequencies drawn from a Beta distribution *f_l_* ~ Beta(0.7,0.7). We run these simulations for different recombination rates by setting the total length of the genetic map per chromosome to either 1, 10 or 100 centimorgans. We then inferred selection on all loci still polymorphic at the end of the simulations with both `BayeScan` and `Flink` for 10 replicates per set.

Across all simulations, `BayeScan` identified no locus under balancing selection and only 0.16% under divergent selection. This low false positive rate is consistent with the generally low power of `BayeScan` to infer loci under balancing selection as well as the used prior odds of 50 in favor of the neutral model. Similar results were obtained with `Flink` on the simulations with high recombination (genetic map of 100 centimorgans), in which case no linkage information could be exploited. Across these simulations, `Flink` inferred no locus under balancing selection and only 0.14% under divergent selection. The number of false positives, however, was rising sharply with decreasing recombination rate. At a genetic map of 10 centimorgan, 5.0% and 2.8% of all loci were wrongly inferred as under balancing and divergent selection, respectively. At a genetic map of only 1 centimorgan and hence tight linkage, the corresponding false positive rates were 22.7% and 7.5%, respectively. These results thus highlight that the power gained by `Flink` in exploiting linkage information also translates into a higher false positive rate in case the model is misspecified. Under such scenarios, other methods such as `hapFLK` (Fariello *et al*. 2013) or `PCAdapt` (Luu *et al*. 2017) are thus more appropriate.

## Application to Humans

To illustrate the usefulness of `Flink` we applied it to SNP data of 46 populations analyzed as part of the Human Genome Diversity Project (HGDP) (Rosenberg N.A. *et al*. 2002; Rosenberg *et al*. 2005) and available at https://www.hagsc.org/hgdp/files.html. We then used `Plink` v1.90 (Chang *et al*. 2015) to transpose the data into vcf files and used the `liftOver` tool of the UCSC Genome Browser (James Kent *et al*. 2002) to convert the coordinates to the human reference GRCh38.

We divided the 46 populations into 6 groups (Table 2) of between 4 and 15 populations each according to genetic landscapes proposed by Peter *et al*. (2017). We then inferred divergent and balancing selection using the hierarchical version of `Flink` on all 22 autosomes, but excluded 5 Mb on each side of the centromer and adjacent to the telomeres. The final data set consists in total of 563,589 SNPs. We analyzed each chromosome arm individually with *α*_max_ = 4.0, *s*_max_ = 10 and using an MCMC chain with 7 · 10^5^ iterations, of which we discarded the first 2 · 10^5^ as burnin. Estimates of hierarchical parameters are shown in Figure S2 and the locus-specific FDRs *q_d_*(*l*) and *q_b_*(*l*) are shown for all loci, all groups as well as the higher hierarchy in Supplementary Figures S4-S42. All regions identified as potential targets for selection are further detailed in Supplementary Files. As summarized in Table 2, we discovered between 759 and 1,889 and between 433 and 1,735 candidate regions for divergent and balancing selection, respectively, spanning together about 10% of the genome.

**Table 2.** Population groups analyzed

| Group | Populations | Divergent |  |  | Balancing |  |  |
| --- | --- | --- | --- | --- | --- | --- | --- |
|  |  | SNPs (%) | Regions | Length<sup>a</sup> | SNPs (%) | Regions | Length<sup>a</sup> |
| Africa | Bantu N.E., Biaka Pygmies, Mandenka, Mbuti Pygmies, San, Yoruba | 8,020 (1.42) | 759 | 16.8 | 8,026 (1.42) | 433 | 30.2 |
| Middle East | Mozabite, Palestinian, Druze, Bedouin | 14,324 (2.54) | 1,137 | 20.6 | 18,432 (3.27) | 848 | 41.2 |
| Europe | Adygei, French, French Basque, North Italian, Orcadian, Russian, Sardinian, Tuscan | 19,128 (3.39) | 1,466 | 22.0 | 37,736 (6.7) | 1,382 | 48.3 |
| America | Colombians, Karitiana, Maya, Pima, Surui | 33,062 (5.87) | 1,889 | 29.8 | 34,499 (6.12) | 1,735 | 39.4 |
| Central Asia | Balochi, Brahui, Burusho, Hazara, Kalash, Makrani, Pathan, Sindhi | 16,663 (2.96) | 1,290 | 22.6 | 25,473 (4.52) | 1,132 | 44.5 |
| East Asia | Uyghur, Dai, Daur, Han, Hezhen, Lahu, Miaozi, Mongola, Naxi, Oroqen, She, Tu, Tujia, Xibo, Yizu | 20,528 (3.64) | 1,832 | 17.3 | 33,678 (5.98) | 1,656 | 35.2 |
| Higher hierarchy | N/A | 24,595 (4.36) | 1,692 | 26.8 | 20,156 (3.58) | 1,074 | 31.2 |

<sup>a</sup> Median length of the regions in kb.

### Comparison with BayeScan

We first validated our results by running `BayeScan` on the same data. We then identified divergent regions as continuous sets of SNP markers that passed an FDR threshold of 0.01 or 0.01 for each method and determined the FDR threshold necessary to identify at least one locus within these regions by the other method. To ensure the observed differences between methods is due to accounting for linkage only, we used the hierarchical version `BayeScanH` (Foll *et al*. 2014) that also implements the same hierarchical island model as `Flink`.

As shown in Figure 5A for selected regions among Europeans, the majority of regions identified by `BayeScanH` were replicated by `Flink` at small FDR thresholds. In contrast, most of the regions identified by `Flink` were not replicated by `BayeScanH`, in line with a higher statistical power for the former. Visual inspection indeed revealed that for most regions identified by `Flink` but not `BayeScanH`, the latter also showed a signal of selection at multiple markers, each of which not passing the FDR threshold individually (see Figure 5B for examples). In contrast, sites identified by `BayeScanH` but not `Flink` usually consisted of a signal at a single site, suggesting many of those are likely false positives (Figure 5C).

**Figure 5.**
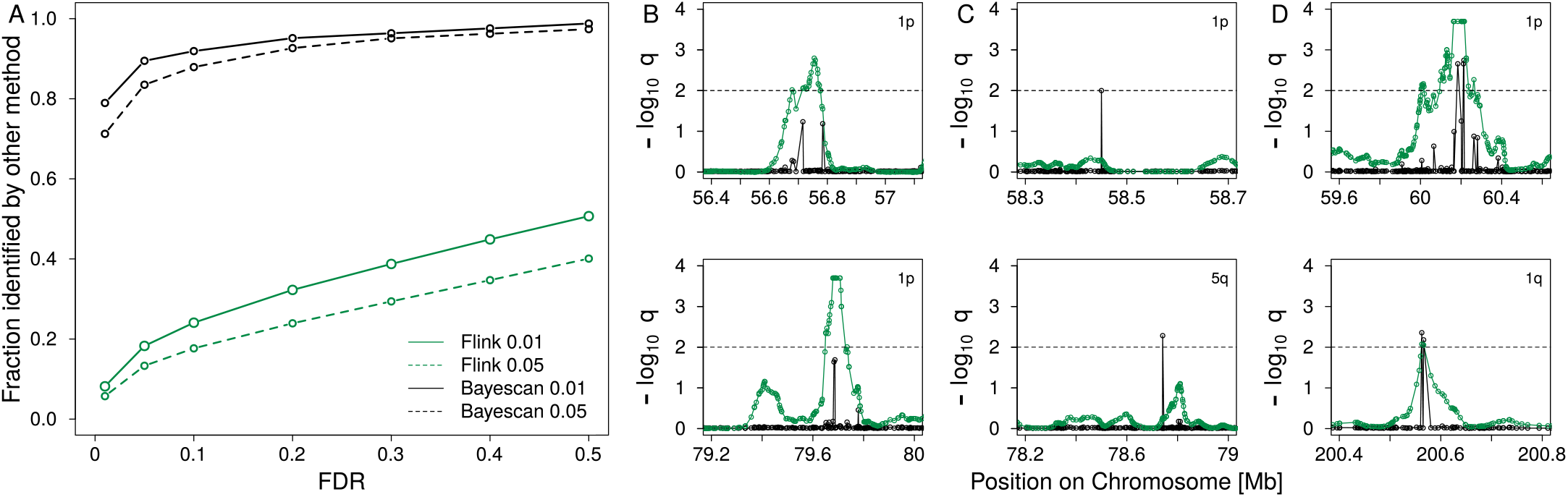
(A) The fraction of regions identified as divergent among Europeans by `Flink` (green) and `Bayescan`H (black) at a false discovery rate (FDR) of 0.01 (solid) and 0.05 (dashed) also identified by the other method at different FDR. (B-D) Examples of regions found under divergent selection by `Flink` (B), `BayeScanH` (C) or both (D) among Europeans. Dashed lines indicate the 0.01 FDR threshold.

Results were similar for the other groups (Figure S3), but the correspondence between the methods was higher for African group and considerably lower for the American group, likely due to the different patterns of divergence among populations (Figure S2).

### Comparison with a recent scan for selective sweeps

Since positive selection might affect a subset of populations only and hence lead to an increase in population differentiation (Nielsen 2005), we compared our outlier regions also to those of a recent scan for positive selection that combined multiple test for selection using a machine learning approach (Sugden *et al*. 2018). Among the 593 candidate loci reported for the CEU population of the 1000 Genomes Project (1000 Genomes Project Consortium *et al*. 2015) and overlapping the chromosomal segments studied here, 293 loci (49.4%) fall within a region we identified as under divergent selection either among European populations (154 loci), at the higher hierarchy (132 loci), or both (7 loci).

To test if this overlap exceeds random expectations, we generated 10,000 bootstrapped data sets by randomly sampling the same amount of loci among all those found polymorphic in the 1000 Genome Project CEU samples and within the chromosomal segments studied here. We then determined the overlap with our outlier regions for each data set. On average, 46.6 loci overlapped with our regions identified among European populations or at the higher hierarchy. Importantly, the largest overlap observed among the bootstrapped data set (72 loci) was much smaller than that observed (293 loci, *P* < 10^−4^).

### Example: The LCT region

As illustration, we show the FDRs *q_d_*(*l*) and *q_b_*(*l*) for 30 Mb around the *LCT* gene in Figure 6 for the higher hierarchy as well as the European, Middle Eastern and East Asian group. The *LCT* gene is a well studied target of positive selection which has acted to increase lactase persistence in several human populations, including Europeans (Nielsen *et al*. 2007). Lactase persistence varies among Europeans and decreases on a roughly north-south cline (Bersaglieri *et al*. 2004; Leonardi *et al*. 2012; Burger *et al*. 2007; Itan *et al*. 2009), consistent with the signal of divergent selection we detected among European populations (Figure 6). In line with previous findings (e.g. Grossman *et al*. 2013), we detected a signal of divergent selection among Europeans also in various genes around *LCT*, most notably in *R3HDM1* but also *MIR128-1, UBXN4* and *DARS*. In contrasts, we detected no such signal for the other groups.

**Figure 6.**
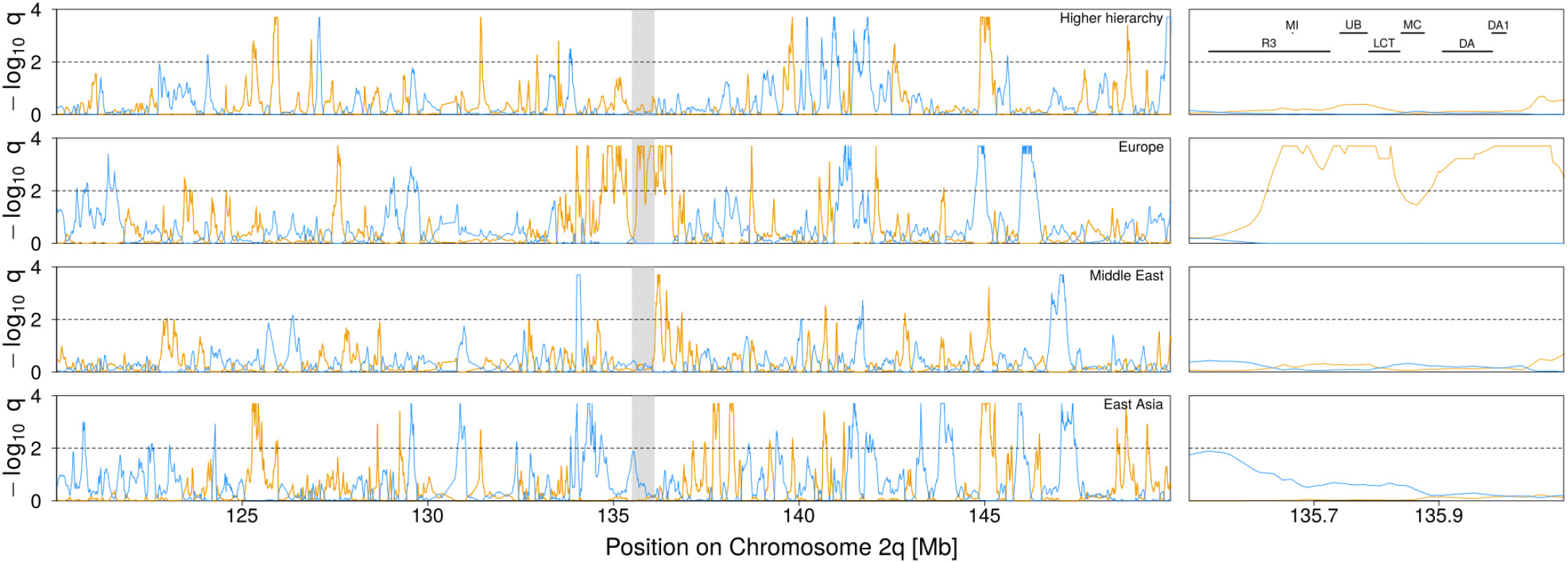
Signal of selection around the *LCT* gene on Chromosome 2q. The orange and blue lines indicate the locus-specific FDR for divergent (orange) and balancing (blue) selection, respectively. The black dashed line shows the 1% FDR threshold. A zoom of the highlighted region is shown on the right indicating the position of several genes: *R3HDM1* (R3), *MIR128-1* (MI), *UBXN4* (UB), *MCM6* (MC), *DARS* (DA) and *DARS-AS1* (DA1). The entire Chromosome 2q is shown in Supplementary Figure S7.

## Discussion

Genome scans are common methods to identifying loci that contribute to local adaptation among populations. Here we extend the particularly powerful method implemented in `BayeScan` (Foll and Gaggiotti 2008) to linked sites.

Accounting for linkage in population genetic methods, while desirable, is often computationally hard. We propose to alleviate this problem by modeling the dependence among linked sites through auto-correlation among hierarchical parameters, rather than the population allele frequencies or haplotypes themselves. In the context of genome scans, this has been previously successfully done by classifying each locus as selected or neutral using Hidden-Markov Models (Boitard *et al*. 2009; Kern and Haussler 2010). Here, we extend this idea by modeling auto-correlation among the strength of selection acting at individual loci. While ignoring auto-correlation at the genetic level certainly leads to a loss of information, the resulting method remains computationally tractable. And as we show with simulations and an application to human data, the resulting method features much improved statistical power compared to `BayeScan`, a similar method that ignores linkage completely.

This is particularly evident for loci with more similar allele frequencies among populations than expected by the genome-wide divergence. These loci are generally interpreted as being under balancing selection (Foll and Gaggiotti 2008; Beaumont and Balding 2004), but may also be the result of purifying selection restricting alleles from reaching high allele frequencies. Given the large number of loci we inferred in this class from the HGDP data (about 5% of the genome), we speculate that balancing selection is unlikely the main driver, and caution against over-interpreting these results. But we note that the empirical false discovery rate for loci under balancing selection was extremely low in our simulations, except if the assumptions underlying F-model was violated.

A benefit of accounting for auto-correlation among locus-specific effects was previously postulated by Guo *et al*. (2009), who proposed a conditional autoregressive (CAR) prior on *α_l_* such that

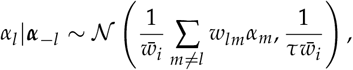

where ***α***_−*l*_ denotes the collection of all other *α_m_*, *m* ≠ *l*, *w_lm_* indicates the covariance between loci *l* and *m*, which is assumed to decrease exponentially with distance, and 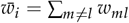.

While Guo *et al*. (2009) did not evaluate the benefit of their CAR implementation on the power of selection inference, they found that it was a better fit to high resolution data. Here we show that the power increase by exploiting auto-correlation among loci is substantial: of all regions identified as under divergent selection by `Flink`, less than halve were also identified by `BayeScan`, despite evidence that these consist mostly of true outliers.

In this context, it is important to note that due to computational challenges, Guo *et al*. (2009) suggested to run their method on low-resolution data with few markers first, and then to apply the CAR version on the so inferred candidate regions only. As our analysis suggests, such an approach would likely fail to harvest the full benefit of accounting for auto-correlation among locus-specific parameters. This is possible using `Flink`, probably because the first-order Markov assumption on locus-specific effects allows for cheap MCMC updates of *α_l_* at a single locus that does not require a recalculation of the prior on the full vector ***α*** = {*α*_1_, …, *α_L_*}. Unfortunately, however,no implementation of the method by Guo *et al*. (2009) is available for a direct comparison.

As second major difference between `Flink` and the CAR method by Guo *et al*. (2009) is that the former discretizes the locus-specific effects *α_l_*. While such a discretization leads to a loss of precision in estimating locus specific effects, it allows to directly calculate a false-discovery rate to identify outlier loci at any desired level of confidence, similar to `BayeScan`. or the method of (Riebler *et al*. 2008). In contrast, the method by Guo *et al*. (2009) identifies outliers indirectly as those for which the posterior distributions on *θ_l_* are significantly different from the distribution of *θ_l_* values under the inferred hyper-parameters. Importantly, the discretization seems to come at no cost on power: in our simulations, `Flink` and `BayeScan` hat virtually identical power if we simulated unlinked data. Also, our simulations show that the parameters of the discrete Markov model proposed here to describe auto-correlation among locus-specific effects are all estimated well. The corresponding parameters of the exponential decay in the model by Guo *et al*. (2009), however, need to be fixed upfront due to numerical instabilities.

An obvious draw-back of modeling the locus-specific selection coefficients as a discrete Markov Chain is that for most candidate regions we detected, multiple loci showed a strong signal of selection, making it difficult to identify the causal variant. However, once a region is identified, estimates of selections coefficients can be obtained for each locus individually to identify the locus with the strongest signal, for which one might then also use complementary methods.

We finally note that the implementation provided through `Flink` allows to group populations hierarchically. Accounting for multiple hierarchies was previously shown to reduce the number of false positives in *F*_ST_ based genome scans (Excoffier *et al*. 2009) and also applied in an *F*-model setting (Foll *et al*. 2014). Aside from accounting for structure more accurately, a hierarchical implementation also allows for genome-wide association studies (GWAS) with population samples. In such a setting, each sampling location would constitute a “group” of, say, two “populations”, one for each phenotype (e.g. cases and controls). The parameters at the higher hierarchy will then accurately describe population structure and loci associated with the phenotype will be identified as those highly divergent between the two “populations”. A natural assumption would then be that the locus-specific coefficients *α_l_* are shared among all groups, i.e. that they are governed by a single HMM. While we have not made use of such a setting here, we note that it is readily available as an option in `Flink`.

## Acknowledgments

We are grateful for the very constructive feedback of two anonymous reviewers on an earlier version of this manuscript. This study was supported by two Swiss National Foundation grants to DW with numbers 31003A_149920 and 31003A_173062.

## Notes

### Competing Interest Statement

The authors have declared no competing interest.

## Literature Cited

1000 Genomes Project Consortium, A. Auton, L. D. Brooks, R. M. Durbin, E. P. Garrison, et al., 2015 A global reference for human genetic variation. Nature 526: 68–74.

Andrew, R. L. and L. H. Rieseberg, 2013 Divergence is focused on few genomic regions early in speciation: Incipient speciation of sunflower ecotypes. Evolution 67: 2468–2482.

Balding, D. J., 2003 Likelihood-based inference for genetic correlation coefficients. Theoretical Population Biology 63: 221 – 230, Uses of DNA and genetic markers for forensics and population studies.

Beaumont, M. and R. A. Nichols, 1996 Evaluating loci for use in the genetic analysis of population structure. P.Roy.Soc.Lond.B 263: 1619–1626.

Beaumont, M. A. and D. J. Balding, 2004 Identifying adaptive genetic divergence among populations from genome scans. Molecular Ecology 13: 969–980.

Bersaglieri, T., P. C. Sabeti, N. Patterson, T. Vanderploeg, S. F. Schaffner, et al., 2004 Genetic Signatures of Strong Recent Positive Selection at the Lactase Gene. The American Journal of Human Genetics 74: 1111–1120.

Boitard, S., C. Schlötterer, and A. Futschik, 2009 Detecting selective sweeps: A new approach based on hidden Markov models. Genetics 181: 1567–1578.

Bonin, A., P. Taberlet, C. Miaud, and F. Pompanon, 2006 Explorative genome scan to detect candidate loci for adaptation along a gradient of altitude in the common frog (Rana temporaria). Mol.Biol.Evol. 23: 773–783.

Burger, J., M. Kirchner, B. Bramanti, W. Haak, and M. G. Thomas, 2007 Absence of the lactase-persistence-associated allele in early Neolithic Europeans. Proceedings of the National Academy of Sciences 104: 3736–3741.

Chang, C. C., C. C. Chow, L. C. Tellier, S. Vattikuti, S. M. Purcell, et al., 2015 Second-generation PLINK: Rising to the challenge of larger and richer datasets. GigaScience 4: 1–16.

Cruickshank, T. E. and M. W. Hahn, 2014 Reanalysis suggests that genomic islands of speciation are due to reduced diversity, not reduced gene flow. Mol.Ecol. 23: 3133–3157.

Durand, E. Y., N. Patterson, D. Reich, and M. Slatkin, 2011 Testing for ancient admixture between closely related populations. Mol.Biol.Evol. 28: 2239–2252.

Eriksson, A. and A. Manica, 2012 Effect of ancient population structure on the degree of polymorphism shared between modern human populations and ancient hominins 109: 13956–13960.

Excoffier, L., T. Hofer, and M. Foll, 2009 Detecting loci under selection in a hierarchically structured population. Heredity 103: 285–298.

Falush, D., M. Stephens, and J. K. Pritchard, 2003 Inference of population structure using multilocus genotype data: Linked loci and correlated allele frequencies. Genetics 164: 1567–1587.

Fariello, M. I., S. Boitard, H. Naya, M. SanCristobal, and B. Servin, 2013 Detecting signatures of selection through haplotype differentiation among hierarchically structured populations. Genetics 193: 929–941.

Feder, J. L., S. P. Egan, and P. Nosil, 2012 The genomics of speciation-with-gene-flow. Trends in Genetics 28: 342–350.

Ferrer-Admetlla, A., C. Leuenberger, J. D. Jensen, and D. Wegmann, 2016 An Approximate Markov Model for the Wright-Fisher Diffusion. Genetics 203: 831–846.

Foll, M. and O. Gaggiotti, 2008 A genome-scan method to identify selected loci appropriate for both dominant and codominant markers: A Bayesian perspective. Genetics 180: 977–993.

Foll, M., O. E. Gaggiotti, J. T. Daub, A. Vatsiou, and L. Excoffier, 2014 Widespread signals of convergent adaptation to high altitude in Asia and America. American Journal of Human Genetics 95: 394–407.

Fournier-Level, A., A. Korte, M. D. Cooper, M. Nordborg, J. Schmitt, et al., 2011 A map of local adaptation in arabidopsis thaliana. Science 334: 86–89.

Gaggiotti, O. E. and M. Foll, 2010 Quantifying population structure using the F-model. Molecular Ecology Resources 10: 821–830.

Grossman, S. R., K. G. Andersen, I. Shlyakhter, S. Tabrizi, S. Winnicki, et al., 2013 Identifying recent adaptations in large-scale genomic data. Cell 152: 703–13.

Guo, F., D. K. Dey, and K. E. Holsinger, 2009 A Bayesian hierarchical model for analysis of SNP diversity in multilocus, multipopulation samples. Journal of the American Statistical Association 104: 142–154.

Itan, Y., A. Powell, M. A. Beaumont, J. Burger, and M. G. Thomas, 2009 The origins of lactase persistence in Europe. PLoS Computational Biology 5: 17–19.

James Kent, W., C. W. Sugnet, T. S. Furey, K. M. Roskin, T. H. Pringle, et al., 2002 The human genome browser at UCSC. Genome Research 12: 996–1006.

Jones, F. C., M. G. Grabherr, Y. F. Chan, P. Russell, E. Mauceli, et al., 2012 The genomic basis of adaptive evolution in three-spine sticklebacks. Nature 484: 55–61.

Kern, A. D. and D. Haussler, 2010 A population genetic hidden markov model for detecting genomic regions under selection. Molecular Biology and Evolution 27: 1673–1685.

Leonardi, M., P. Gerbault, M. G. Thomas, and J. Burger, 2012 The evolution of lactase persistence in Europe. A synthesis of archaeological and genetic evidence. International Dairy Journal 22: 88–97.

Lewontin, R. C. and J. Krakauer, 1973 Distribution of gene frequency as a test of the theory of the selective neutrality of polymorphisms. Genetics 74: 175–195.

Lotterhos, K. E. and M. C. Whitlock, 2014 Evaluation of demographic history and neutral parameterization on the performance of fst outlier tests. Molecular Ecology 23: 2178–2192.

Luu, K., E. Bazin, and M. G. B. Blum, 2017 pcadapt: an r package to perform genome scans for selection based on principal component analysis. Molecular Ecology Resources 17: 67–77.

Nei, M. and T. Maruyama, 1975 Lewontin-krakauer test for neutral genes. Genetics 80: 395–395.

Neuenschwander, S., F. Michaud, and J. Goudet, 2018 QuantiNemo 2: a Swiss knife to simulate complex demographic and genetic scenarios, forward and backward in time. Bioinformatics 35: 886–888.

Nielsen, R., 2005 Molecular signatures of natural selection. Annual Review of Genetics 39: 197–218, PMID: 16285858.

Nielsen, R., I. Hellmann, M. Hubisz, C. Bustamante, and A. G. Clark, 2007 Recent and ongoing selection in the human genome. Nature Reviews Genetics 8: 857–868.

Peter, B. M., 2016 Admixture, population structure, and f-statistics. Genetics 202: 1485–1501.

Peter, B. M., D. Petkova, and J. Novembre, 2017 Genetic landscapes reveal how human genetic diversity aligns with geography. bioRxiv pp. 1–24.

Rannala, B. H. and J. A. Hartigan, 1996 Estimating gene flow in island populations. Genetical Research 67: 147–158.

Riebler, A., L. Held, and W. Stephan, 2008 Bayesian variable selection for detecting adaptive genomic differences among populations. Genetics 178: 1817–1829.

Rosenberg, N. A., S. Mahajan, S. Ramachandran, C. Zhao, J. K. Pritchard, et al., 2005 Clines, clusters, and the effect of study design on the inference of human population structure. PLoS Genetics 1: 0660–0671.

Rosenberg N.A., Pritchard J.K., Weber J.L., Cann H.M., Kidd K.K., et al., 2002 Genetic structure of human populations. Science 298: 2981–2985.

Sabeti, P. C., D. E. Reich, J. M. Higgins, H. Z. P. Levine, D. J. Richter, et al., 2002 Detecting recent positive selection in the human genome from haplotype structure. Nature 419: 832–837.

Sabeti, P. C., P. Varilly, B. Fry, J. Lohmueller, E. Hostetter, et al., 2007 Genome-wide detection and characterization of positive selection in human populations. Nature 449: 913–918.

Slatkin, M. and L. Voelm, 1991 FST in a hierarchical island model. Genetics 127: 627–9.

Stölting, K. N., R. Nipper, D. Lindtke, C. Caseys, S. Waeber, et al., 2013 Genomic scan for single nucleotide polymorphisms reveals patterns of divergence and gene flow between ecologically divergent species. Mol.Ecol. 22: 842–855.

Sugden, L. A., E. G. Atkinson, A. P. Fischer, S. Rong, B. M. Henn, et al., 2018 Localization of adaptive variants in human genomes using averaged one-dependence estimation. Nature Communications 9: 1–14.

Tang, K., K. R. Thornton, and M. Stoneking, 2007 A new approach for using genome scans to detect recent positive selection in the human genome. PLOS Biology 5: 1–16.

Voight, B. F., S. Kudaravalli, X. Wen, and J. K. Pritchard, 2006 A map of recent positive selection in the human genome. PLoS biology 4: e72.

Wu, 2001 The genic view of the process of speciation. J.Evol.Biol. 14: 851–865.

