## Supplementary Information for "Detecting selection from linked sites using an F-model"

### Detecting selection from linked sites using an F-model Supplemental Material

[*]Department of Biology and Biochemistry, University of Fribourg, Fribourg, Switzerland, [†]Swiss Institute of Bioinformatics, Fribourg, Switzerland, [‡]Department of Mathematics, University of Fribourg, Fribourg, Switzerland, [§]iCoSys, University of Applied Sciences Western Switzerland, Fribourg, Switzerland, [**]Department of Informatics, University of Medicine, Pharmacy, Science and Technology of Târgu Mureş, Târgu Mureş, Romania, [††]International Agency for Research on Cancer (IARC-WHO), Lyon, France

#### MCMC inference

New parameters are proposed using a symmetric transition kernel which will thus not appear in the Hastings ratios. The proposed parameter will be accepted with a probability given by the minimum between 1 and the Hastings ratio.

##### *Updating* $\beta_{gj}$

Propose a new $\tilde{\beta}_{gj}$, recalculate $\tilde{\theta}_{gjl}$ using equation 2 for all $l = 1, \ldots, L$, and use equation 5 to calculate the Hastings ratio

$$\begin{aligned} h &= \frac{\pi(\tilde{\beta}_{gj})}{\pi(\beta_{gj})} \prod_{l=1}^{L} \frac{\mathbb{P}(n_{gjl}|\tilde{\theta}_{gjl}, p_{gl})}{\mathbb{P}(n_{gjl}|\theta_{gjl}, p_{gl})} \\ &= \frac{\pi(\tilde{\beta}_{gj})}{\pi(\beta_{gj})} \prod_{l=1}^{L} \frac{\Gamma(\tilde{\theta}_{gjl} p_{gl} + n_{gjl})}{\Gamma(\tilde{\theta}_{gjl} p_{gl})} \frac{\Gamma(\tilde{\theta}_{gjl}(1-p_{gl}) + N_{gjl} - n_{gjl})}{\Gamma(\tilde{\theta}_{gjl}(1-p_{gl}))} \frac{\Gamma(\tilde{\theta}_{gjl})}{\Gamma(\tilde{\theta}_{gjl} + N_{gjl})} \\ &\cdot \frac{\Gamma(\theta_{gjl} p_{gl})}{\Gamma(\theta_{gjl} p_{gl} + n_{gjl})} \frac{\Gamma(\theta_{gjl}(1-p_{gl}))}{\Gamma(\theta_{gjl}(1-p_{gl}) + N_{gjl} - n_{gjl})} \frac{\Gamma(\theta_{gjl} + N_{gjl})}{\Gamma(\theta_{gjl})} \end{aligned}$$

Using the functional equation $\Gamma(n+1) = n\Gamma(n)$, we obtain

$$\begin{aligned} h &= \frac{\pi(\tilde{\beta}_{gj})}{\pi(\beta_{gj})} \prod_{l=1}^{L} \frac{\overbrace{((\tilde{\theta}_{gjl} p_{gl} + n_{gjl}) - 1) \cdot ((\tilde{\theta}_{gjl} p_{gl} + n_{gjl}) - 2) \ldots (\tilde{\theta}_{gjl} p_{gl})}^{n_{gjl} \text{ elements}}}{((\theta_{gjl} p_{gl} + n_{gjl}) - 1) \cdot ((\theta_{gjl} p_{gl} + n_{gjl}) - 2) \ldots (\theta_{gjl} p_{gl})} \\ &\cdot \underbrace{\frac{(\tilde{\theta}_{gjl}(1-p_{gl}) + N_{gjl} - n_{gjl} - 1) \ldots (\tilde{\theta}_{gjl}(1-p_{gl}))}{(\theta_{gjl}(1-p_{gl}) + N_{gjl} - n_{gjl} - 1) \ldots (\theta_{gjl}(1-p_{gl}))}}_{N_{gjl} - n_{gjl} \text{ elements}} \cdot \underbrace{\frac{(\theta_{gjl} + N_{gjl} - 1) \ldots \theta_{gjl}}{(\tilde{\theta}_{gjl} + N_{gjl} - 1) \ldots \tilde{\theta}_{gjl}}}_{N_{gjl} \text{ elements}} \\ &= \frac{\pi(\tilde{\beta}_{gj})}{\pi(\beta_{gj})} \prod_{l=1}^{L} \left( \prod_{i=0}^{n_{gjl}-1} \frac{\tilde{\theta}_{gjl} p_{gl} + i}{\theta_{gjl} p_{gl} + i} \cdot \prod_{i=0}^{N_{gjl} - n_{gjl} - 1} \frac{\tilde{\theta}_{gjl}(1-p_{gl}) + i}{\theta_{gjl}(1-p_{gl}) + i} \cdot \prod_{i=0}^{N_{gjl}-1} \frac{\theta_{gjl} + i}{\tilde{\theta}_{gjl} + i} \right) \end{aligned}$$

This ratio is evaluated as a logarithmic sum:

$$\begin{aligned} \log h = \log \frac{\pi(\tilde{\beta}_{gj})}{\pi(\beta_{gj})} + \sum_{l=1}^{L} \left[ \sum_{i=0}^{n_{gjl}-1} \log \left( \frac{\tilde{\theta}_{gjl} p_{gl} + i}{\theta_{gjl} p_{gl} + i} \right) + \sum_{i=0}^{N_{gjl} - n_{gjl} - 1} \log \left( \frac{\tilde{\theta}_{gjl}(1-p_{gl}) + i}{\theta_{gjl}(1-p_{gl}) + i} \right) \right. \\ \left. + \sum_{i=0}^{N_{gjl}-1} \log \left( \frac{\theta_{gjl} + i}{\tilde{\theta}_{gjl} + i} \right) \right] \end{aligned}$$

In the case when we have a big value of $N$, we can approximate a sum $\sum_{i=0}^{N} \log(x+i)$ as the integral $\int_0^N \log(c+k)dk$. To calculate this approximation, we use the Euler-Maclaurin formula

$$\begin{aligned} \sum_{i=0}^{N-1} \log(x+i) &\approx \int_0^N \log(c+k)dk + \frac{1}{2}\left[-\log(N+c) + \log(c)\right] + \frac{1}{12}\left[\frac{1}{N+c} - \frac{1}{c}\right] \\ &= (N+c-0.5)\log(N+c) - (c-0.5)\log(c) - N + \frac{1}{12}\left[\frac{1}{N+c} - \frac{1}{c}\right] \end{aligned}$$

where the $-N$ term will be canceled developing the approximation for the numerator and denominator in the argument of the logarithms in the equation of the Hastings ratio logarithm.

***Updating*** $p_{gl}$

Propose a new $\tilde{p}_{gl}$ and use equation 5 and equation 8 to calculate the Hastings ratio

$$h = \frac{\mathbb{P}(\tilde{p}_{gl}|P_l, \Theta_{gl})}{\mathbb{P}(p_{gl}|P_l, \Theta_{gl})} \prod_{j=1}^{J_g} \frac{\mathbb{P}(n_{gjl}|\theta_{gjl}, \tilde{p}_{gl})}{\mathbb{P}(n_{gjl}|\theta_{gjl}, p_{gl})}$$

Note that the Gamma functions in equation 8 cancel for the ratio $\frac{\mathbb{P}(\tilde{p}_{gl}|P_l,\Theta_{gl})}{\mathbb{P}(p_{gl}|P_l,\Theta_{gl})}$

$$\begin{aligned} h &= \left(\frac{\tilde{p}_{gl}}{p_{gl}}\right)^{\Theta_{gl}P_l - 1} \left(\frac{1-\tilde{p}_{gl}}{1-p_{gl}}\right)^{\Theta_{gl}(1-P_l)-1} \prod_{j=1}^{J_g} \frac{\Gamma(\theta_{gjl}p_{gl})}{\Gamma(\theta_{gjl}\tilde{p}_{gl})} \frac{\Gamma(\theta_{gjl}(1-p_{gl}))}{\Gamma(\theta_{gjl}(1-\tilde{p}_{gl}))} \\ &\cdot \frac{\Gamma(\theta_{gjl}\tilde{p}_{gl} + n_{gjl})}{\Gamma(\theta_{gjl}p_{gl} + n_{gjl})} \frac{\Gamma(\theta_{gjl}(1-\tilde{p}_{gl}) + N_{gjl} - n_{gjl})}{\Gamma(\theta_{gjl}(1-p_{gl}) + N_{gjl} - n_{gjl})} \end{aligned}$$

As we did it before, we can simplify the ratio between the Gamma functions using the functional equation $\Gamma(n+1) = n\Gamma(n)$

$$\begin{aligned} h &= \left(\frac{\tilde{p}_{gl}}{p_{gl}}\right)^{\Theta_{gl}P_l - 1} \left(\frac{1-\tilde{p}_{gl}}{1-p_{gl}}\right)^{\Theta_{gl}(1-P_l)-1} \prod_{j=1}^{J_g} \overbrace{\frac{(\theta_{gjl}\tilde{p}_{gl} + n_{gjl} - 1)...(\theta_{gjl}\tilde{p}_{gl} + 1) \cdot (\theta_{gjl}\tilde{p}_{gl})}{(\theta_{gjl}p_{gl} + n_{gjl} - 1)...(\theta_{gjl}p_{gl} + 1) \cdot (\theta_{gjl}p_{gl})}}^{n_{gjl} \text{ elements}} \\ &\cdot \underbrace{\frac{(\theta_{gjl}(1-\tilde{p}_{gl}) + N_{gjl} - n_{gjl} - 1)...(\theta_{gjl}(1-\tilde{p}_{gl}) + 1) \cdot (\theta_{gjl}(1-\tilde{p}_{gl}))}{(\theta_{gjl}(1-p_{gl}) + N_{gjl} - n_{gjl} - 1)...(\theta_{gjl}(1-p_{gl}) + 1) \cdot (\theta_{gjl}(1-p_{gl}))}}_{N_{gjl} - n_{gjl} \text{ elements}} \\ &= \left(\frac{\tilde{p}_{gl}}{p_{gl}}\right)^{\Theta_{gl}P_l - 1} \left(\frac{1-\tilde{p}_{gl}}{1-p_{gl}}\right)^{\Theta_{gl}(1-P_l)-1} \cdot \prod_{j=1}^{J_g} \left( \prod_{i=0}^{n_{gjl}-1} \frac{\theta_{gjl}\tilde{p}_{gl} + i}{\theta_{gjl}p_{gl} + i} \cdot \prod_{i=0}^{N_{gjl}-n_{gjl}-1} \frac{\theta_{gjl}(1-\tilde{p}_{gl}) + i}{\theta_{gjl}(1-p_{gl}) + i} \right) \end{aligned}$$

Calculating the logarithm of $h$ we obtain

$$\begin{aligned} \log h &= (\Theta_{gl}P_l - 1) \log \left(\frac{\tilde{p}_{gl}}{p_{gl}}\right) + (\Theta_{gl}(1-P_l) - 1) \log \left(\frac{1-\tilde{p}_{gl}}{1-p_{gl}}\right) + \sum_{j=1}^{J_g} \left( \sum_{i=0}^{n_{gjl}-1} \log \left(\frac{\theta_{gjl}\tilde{p}_{gl} + i}{\theta_{gjl}p_{gl} + i}\right) \right. \\ &+ \left. \sum_{i=0}^{N_{gjl}-n_{gjl}-1} \log \left(\frac{\theta_{gjl}(1-\tilde{p}_{gl}) + i}{\theta_{gjl}(1-p_{gl}) + i}\right) \right) \end{aligned}$$

at which equation we can apply the approximation of the sums as integrals under the established conditions.

***Updating*** $p_{gl}$ ***without hierarchy (the case when g=1)***

Propose a new $\tilde{p}_{gl}$ and calculate the Hastings ratio

$$h = \frac{\pi(\tilde{p}_{gl})}{\pi(p_{gl})} \prod_{j=1}^{J_g} \frac{\mathbb{P}(n_{gjl}|\theta_{gjl}, \tilde{p}_{gl})}{\mathbb{P}(n_{gjl}|\theta_{gjl}, p_{gl})}$$

$$h = \frac{\pi(\tilde{p}_{gl})}{\pi(p_{gl})} \prod_{j=1}^{J_g} \frac{\Gamma(\theta_{gjl}p_{gl})}{\Gamma(\theta_{gjl}\tilde{p}_{gl})} \frac{\Gamma(\theta_{gjl}(1-p_{gl}))}{\Gamma(\theta_{gjl}(1-\tilde{p}_{gl}))} \frac{\Gamma(\theta_{gjl}\tilde{p}_{gl} + n_{gjl})}{\Gamma(\theta_{gjl}p_{gl} + n_{gjl})} \frac{\Gamma(\theta_{gjl}(1-\tilde{p}_{gl}) + N_{gjl} - n_{gjl})}{\Gamma(\theta_{gjl}(1-p_{gl}) + N_{gjl} - n_{gjl})}$$

As we did it before, we can simplify the ratio between the Gamma functions using the functional equation $\Gamma(n+1) = n\Gamma(n)$

$$
\begin{aligned}
h &= \frac{\pi(\tilde{p}_{gl})}{\pi(p_{gl})} \prod_{j=1}^{J_g} \frac{\overbrace{(\theta_{gjl}\tilde{p}_{gl} + n_{gjl} - 1)...(\theta_{gjl}\tilde{p}_{gl} + 1) \cdot (\theta_{gjl}\tilde{p}_{gl})}^{n_{gjl} \text{ elements}}}{(\theta_{gjl}p_{gl} + n_{gjl} - 1)...(\theta_{gjl}p_{gl} + 1) \cdot (\theta_{gjl}p_{gl})} \\
&\cdot \frac{(\theta_{gjl}(1-\tilde{p}_{gl}) + N_{gjl} - n_{gjl} - 1)...(\theta_{gjl}(1-\tilde{p}_{gl}) + 1) \cdot (\theta_{gjl}(1-\tilde{p}_{gl}))}{\underbrace{(\theta_{gjl}(1-p_{gl}) + N_{gjl} - n_{gjl} - 1)...(\theta_{gjl}(1-p_{gl}) + 1) \cdot (\theta_{gjl}(1-p_{gl}))}_{N_{gjl}-n_{gjl} \text{ elements}}} \\
&= \frac{\pi(\tilde{p}_{gl})}{\pi(p_{gl})} \prod_{j=1}^{J_g} \left( \prod_{i=0}^{n_{gjl}-1} \frac{\theta_{gjl}\tilde{p}_{gl} + i}{\theta_{gjl}p_{gl} + i} \cdot \prod_{i=0}^{N_{gjl}-n_{gjl}-1} \frac{\theta_{gjl}(1-\tilde{p}_{gl}) + i}{\theta_{gjl}(1-p_{gl}) + i} \right)
\end{aligned}
$$

Calculating the logarithm of $h$ we obtain

$$
\log h = \log \frac{\pi(\tilde{p}_{gl})}{\pi(p_{gl})} + \sum_{j=1}^{J_g} \left( \sum_{i=0}^{n_{gjl}-1} \log\left(\frac{\theta_{gjl}\tilde{p}_{gl} + i}{\theta_{gjl}p_{gl} + i}\right) + \sum_{i=0}^{N_{gjl}-n_{gjl}-1} \log\left(\frac{\theta_{gjl}(1-\tilde{p}_{gl}) + i}{\theta_{gjl}(1-p_{gl}) + i}\right) \right)
$$

at which equation we can apply the approximation of the sums as integrals under the established conditions. Cause we are using a beta distribution as a prior ($p_{gl} \sim beta(a, b)$, where the parameter $a$ and $b$ describe the shape of allele frequencies in the ancestral population), we can replace the term representing the fraction between the two priors

$$
\log h = (a-1)\log\frac{\tilde{p}_{gl}}{p_{gl}} + (b-1)\log\frac{1-\tilde{p}_{gl}}{1-p_{gl}} + \sum_{j=1}^{J_g} \left( \sum_{i=0}^{n_{gjl}-1} \log\left(\frac{\theta_{gjl}\tilde{p}_{gl} + i}{\theta_{gjl}p_{gl} + i}\right) + \sum_{i=0}^{N_{gjl}-n_{gjl}-1} \log\left(\frac{\theta_{gjl}(1-\tilde{p}_{gl}) + i}{\theta_{gjl}(1-p_{gl}) + i}\right) \right)
$$

###### ***Updating*** $S_{g,l}$

Propose a new $\tilde{S}_{g,l}$, recalculate $\tilde{\theta}_{gjl}$ using equation 2, and use equation 5 to calculate the Hastings ratio

$$
h = \frac{\mathbb{P}(\tilde{S}_{g,l}|S_{g,l-1}, \kappa_g, d_l)\mathbb{P}(S_{g,l+1}|\tilde{S}_{g,l}, \kappa_g, d_{l+1})}{\mathbb{P}(S_{g,l}|S_{g,l-1}, \kappa_g, d_l)\mathbb{P}(S_{g,l+1}|S_{g,l}, \kappa_g, d_{l+1})} \prod_{j=1}^{J_g} \frac{\mathbb{P}(n_{gjl}|\tilde{\theta}_{gjl}, p_{gl})}{\mathbb{P}(n_{gjl}|\theta_{gjl}, p_{gl})}
$$

Using the results that we have obtained previously, we can write the Hastings ratio as

$$
\begin{aligned}
\log h &= \log\left(\frac{Q_{gl}(S_{g,l-1} \to \tilde{S}_{g,l})Q_{gl+1}(\tilde{S}_{g,l} \to S_{g,l+1})}{Q_{gl}(S_{g,l-1} \to S_{g,l})Q_{gl+1}(S_{g,l} \to S_{g,l+1})}\right) + \sum_{j=1}^{J_g} \left( \sum_{i=0}^{n_{gjl}-1} \log\left(\frac{\tilde{\theta}_{gjl}p_{gl} + i}{\theta_{gjl}p_{gl} + i}\right) \right. \\
&+ \left. \sum_{i=0}^{N_{gjl}-n_{gjl}-1} \log\left(\frac{\tilde{\theta}_{gjl}(1-p_{gl}) + i}{\theta_{gjl}(1-p_{gl}) + i}\right) + \sum_{i=0}^{N_{gjl}-1} \log\left(\frac{\theta_{gjl} + i}{\tilde{\theta}_{gjl} + i}\right) \right)
\end{aligned}
$$

In this case we have to calculate all the elements of the matrices $\tilde{Q}_{gl}$ and $Q_{gl}$.

###### ***Updating*** $\ln \kappa_g$

Propose a new $\ln\tilde{\kappa}_g$ and calculate the Hastings ratio

$$
h = \prod_{l=1}^{L} \frac{\mathbb{P}(S_{g,l}|S_{g,l-1}, \ln\tilde{\kappa}_g, d_l)}{\mathbb{P}(S_{g,l}|S_{g,l-1}, \ln\kappa_g, d_l)}
$$

where $\mathbb{P}(S_{g,l}|S_{g,l-1}, \ln\tilde{\kappa}_g, d_l)$ means that we are using the transition matrix $\tilde{Q}_{gl}$ for the new $\ln\tilde{\kappa}_g$. Calling $Q_{gl}(S_{g,l-1} \to S_{g,l})$ as the probability to go from the states $S_{g,l-1}$ to the states $S_{g,l}$, and supposing to take the same priors for the matrices $\tilde{Q}_{gl}(S_{g,1} \to S_{g,2})$ and $Q_{gl}(S_{g,1} \to S_{g,2})$, we can write the Hastings ratio as

$$
h = \prod_{l=2}^{L} \frac{\tilde{Q}_{gl}(S_{g,l-1} \to S_{g,l})}{Q_{gl}(S_{g,l-1} \to S_{g,l})}
$$

To make the calculation faster, we can calculate only the element of the $Q_{gl}$ matrix (and the same for the $\tilde{Q}_{gl}$ matrix) in the i-*th* row and j-*th* column

$$Q_{gl_{i,j}} = \sum_{k=1}^{m} H_{i,k} \cdot D_{k,k} \cdot H_{k,j}^{T} \qquad \forall\, i,j,k \in \{1,2,..,m\}$$

where $D$ is the diagonal matrix which we have seen before: $D = \text{diag}(e^{\kappa_g d_l \lambda_1}, \ldots e^{\kappa_g d_l \lambda_m})$.

**Updating** $B_g$

Propose a new $\tilde{B}_g$, recalculate $\tilde{\Theta}_{gl}$ using equation 9 for all $l = 1,\ldots,L$, and use equation 8 to calculate the Hastings ratio

$$\begin{aligned} h &= \frac{\pi(\tilde{B}_g)}{\pi(B_g)} \prod_{l=1}^{L} \frac{\mathbb{P}(p_{gl}|P_l,\tilde{\Theta}_{gl})}{\mathbb{P}(p_{gl}|P_l,\Theta_{gl})} \\ &= \frac{\pi(\tilde{B}_g)}{\pi(B_g)} \prod_{l=1}^{L} \frac{\Gamma(\tilde{\Theta}_{gl})\Gamma(\Theta_{gl}P_l)\Gamma(\Theta_{gl}(1-P_l))}{\Gamma(\Theta_{gl})\Gamma(\tilde{\Theta}_{gl}P_l)\Gamma(\tilde{\Theta}_{gl}(1-P_l))} p_{gl}^{(\tilde{\Theta}_{gl}-\Theta_{gl})P_l}(1-p_{gl})^{(\tilde{\Theta}_{gl}-\Theta_{gl})(1-P_l)} \end{aligned}$$

This ratio be done evaluated as a logarithmic sum.

**Updating** $P_l$

Propose a new $\tilde{P}_l$ and use equation 8 to calculate the Hastings ratio

$$h = \frac{\pi(\tilde{P}_l)}{\pi(P_l)} \prod_{g=1}^{G} \frac{\mathbb{P}(p_{gl}|\tilde{P}_l,\Theta_{gl})}{\mathbb{P}(p_{gl}|P_l,\Theta_{gl})}$$

After the substitutions and the calculation we obtain

$$\log h = \log \frac{\pi(\tilde{P}_l)}{\pi(P_l)} + \sum_{g=1}^{G} \left( \log \frac{\Gamma(\Theta_{gl}P_l)\Gamma(\Theta_{gl}(1-P_l))}{\Gamma(\Theta_{gl}\tilde{P}_l)\Gamma(\Theta_{gl}(1-\tilde{P}_l))} + \Theta_{gl}(\tilde{P}_l - P_l)\log\left(\frac{p_{gl}}{1-p_{gl}}\right)\right)$$

Cause we are using a beta distribution as a prior ($P_l \sim beta(a,b)$, where the parameter $a$ and $b$ describe the shape of allele frequencies in the ancestral population), we can replace the term representing the fraction between the two priors

$$\log h = (a-1)\log\frac{\tilde{P}_l}{P_l} + (b-1)\log\frac{1-\tilde{P}_l}{1-P_l} + \sum_{g=1}^{G} \left( \log \frac{\Gamma(\Theta_{gl}P_l)\Gamma(\Theta_{gl}(1-P_l))}{\Gamma(\Theta_{gl}\tilde{P}_l)\Gamma(\Theta_{gl}(1-\tilde{P}_l))} + \Theta_{gl}(\tilde{P}_l - P_l)\log\left(\frac{p_{gl}}{1-p_{gl}}\right)\right)$$

**Updating** $S_l$

Propose a new $\tilde{S}_l$, recalculate $\tilde{\Theta}_{gl}$ using equation 9, and use equation 8 to calculate the Hastings ratio

$$h = \frac{\mathbb{P}(\tilde{S}_l|S_{l-1},\kappa,d_l)\mathbb{P}(S_{l+1}|\tilde{S}_l,\kappa,d_{l+1})}{\mathbb{P}(S_l|S_{l-1},\kappa,d_l)\mathbb{P}(S_{l+1}|S_l,\kappa,d_{l+1})} \prod_{g=1}^{G} \frac{\mathbb{P}(p_{gl}|P_l,\tilde{\Theta}_{gl})}{\mathbb{P}(p_{gl}|P_l,\Theta_{gl})}$$

Using the results that we have obtained previously, we can write the Hastings ratio as

$$\begin{aligned} \log h &= \log\left(\frac{Q(S_{l-1}\to\tilde{S}_l)Q(\tilde{S}_l\to S_{l+1})}{Q(S_{l-1}\to S_l)Q(S_l\to S_{l+1})}\right) + \sum_{g=1}^{G}\Big((\tilde{\Theta}_{gl}-\Theta_{gl})P_l\log p_{gl} + (\tilde{\Theta}_{gl}-\Theta_{gl})(1-P_l)\log(1-p_{gl}) \\ &+ \log\left(\frac{\Gamma(\tilde{\Theta}_{gl})\Gamma(\Theta_{gl}P_l)\Gamma(\Theta_{gl}(1-P_l))}{\Gamma(\Theta_{gl})\Gamma(\tilde{\Theta}_{gl}P_l)\Gamma(\tilde{\Theta}_{gl}(1-P_l))}\right)\Big) \end{aligned}$$

In this case we have to calculate all the elements of the matrices $\tilde{Q}_l$ and $Q_l$.

***Updating*** $\ln \kappa$

Propose a new $\ln \tilde{\kappa}$ and calculate the Hastings ratio

$$h = \prod_{l=1}^{L} \frac{\mathbb{P}(S_l | S_{l-1}, \ln \tilde{\kappa}, d_l)}{\mathbb{P}(S_l | S_{l-1}, \ln \kappa, d_l)}$$

using the same procedure that we used for $\ln \kappa_g$, where $\mathbb{P}(S_l | S_{l-1}, \ln \tilde{\kappa}_g, d_l)$ means that we are using the transition matrix $\tilde{Q}_l$ for the new $\ln \tilde{\kappa}$. As we did for $\ln \kappa_g$, we can write the Hastings ratio as

$$h = \prod_{l=2}^{L} \frac{\tilde{Q}_l(S_{l-1} \to S_l)}{Q_l(S_{l-1} \to S_l)}$$

and to make the calculation faster, we can calculate only the element of the $Q_l$ matrix (and the same for the $\tilde{Q}_l$ matrix) in the i-*th* row and j-*th* column as we saw before.

***Updating*** $\mu$ ***and*** $\nu$

For the two parameters $\mu$ and $\nu$ we use the same procedure. Let's consider $\mu$. We propose a new $\tilde{\mu}$ and we use the Hastings ratios that we have already found in the updating of $\kappa$ and $\kappa_g$

$$\log h = \frac{\pi(\tilde{\mu})}{\pi(\mu)} \log \prod_{l=2}^{L} \frac{\tilde{Q}_l(S_{l-1} \to S_l)}{Q_l(S_{l-1} \to S_l)} + \sum_{g} \log \prod_{l=2}^{L} \frac{\tilde{Q}_{gl}(S_{g,l-1} \to S_{g,l})}{Q_{gl}(S_{g,l-1} \to S_{g,l})}$$

We use the same hasting ratio when we propose a new $\tilde{\nu}$.

***Updating*** $a$ ***without hierarchy (the case when g=1)***

We are using a beta distribution $beta(a, b)$ as a prior of $p_{gl}$, where the parameter $a$ describes the shape of allele frequencies in the ancestral population. To update the parameter $a$ we use the following Hastings ratios

$$h = \frac{\pi(\tilde{a})}{\pi(a)} \prod_{l=1}^{L} \frac{\mathbb{P}(p_{gl} | \tilde{a})}{\mathbb{P}(p_{gl} | a)}$$

Calculating the logarithm of $h$ we obtain

$$\begin{aligned} \log h &= \log \frac{\pi(\tilde{a})}{\pi(a)} + \log \prod_{l=1}^{L} \frac{\Gamma(\tilde{a}+b) \cdot \Gamma(a)}{\Gamma(\tilde{a}) \cdot \Gamma(a+b)} \cdot p_{gl}^{(\tilde{a}-a)} \\ &= \log \frac{\pi(\tilde{a})}{\pi(a)} + \sum_{l=1}^{L} \left[ \log \frac{\Gamma(\tilde{a}+b) \cdot \Gamma(a)}{\Gamma(\tilde{a}) \cdot \Gamma(a+b)} + (\tilde{a}-a) \log p_{gl} \right] \\ &= \log \frac{\pi(\tilde{a})}{\pi(a)} + L \cdot \log \frac{\Gamma(\tilde{a}+b) \cdot \Gamma(a)}{\Gamma(\tilde{a}) \cdot \Gamma(a+b)} + \sum_{l=1}^{L} (\tilde{a}-a) \log p_{gl} \end{aligned}$$

***Updating*** $a$ ***with hierarchy***

In this case we are using a beta distribution $beta(a, b)$ as a prior of $P_l$, where the parameter $a$ describes the shape of allele frequencies in the ancestral population. To update the parameter $a$ we use the following Hastings ratios

$$h = \frac{\pi(\tilde{a})}{\pi(a)} \prod_{l=1}^{L} \frac{\mathbb{P}(P_l | \tilde{a})}{\mathbb{P}(P_l | a)}$$

Calculating the logarithm of $h$ we obtain

$$\begin{aligned} \log h &= \log \frac{\pi(\tilde{a})}{\pi(a)} + \log \prod_{l=1}^{L} \frac{\Gamma(\tilde{a}+b) \cdot \Gamma(a)}{\Gamma(\tilde{a}) \cdot \Gamma(a+b)} \cdot P_l^{(\tilde{a}-a)} \\ &= \log \frac{\pi(\tilde{a})}{\pi(a)} + L \cdot \log \frac{\Gamma(\tilde{a}+b) \cdot \Gamma(a)}{\Gamma(\tilde{a}) \cdot \Gamma(a+b)} + \sum_{l=1}^{L} (\tilde{a}-a) \log P_l \end{aligned}$$

**Updating $b$ without hierarchy (the case when g=1)**

We are using a beta distribution $beta(a,b)$ as a prior of $p_{gl}$, where the parameter $b$ describes the shape of allele frequencies in the ancestral population. To update the parameter $b$ we use the following Hastings ratios

$$h = \frac{\pi(\tilde{b})}{\pi(b)} \prod_{l=1}^{L} \frac{\mathbb{P}(p_{gl}|\tilde{b})}{\mathbb{P}(p_{gl}|b)}$$

Calculating the logarithm of $h$ we obtain

$$\log h = \log \frac{\pi(\tilde{b})}{\pi(b)} + L \cdot \log \frac{\Gamma(a+\tilde{b}) \cdot \Gamma(b)}{\Gamma(\tilde{b}) \cdot \Gamma(a+b)} + \sum_{l=1}^{L} (\tilde{b} - b) \log(1 - p_{gl})$$

**Updating $b$ with hierarchy**

In this case we are using a beta distribution $beta(a,b)$ as a prior of $P_l$, where the parameter $b$ describes the shape of allele frequencies in the ancestral population. To update the parameter $b$ we use the following Hastings ratios

$$h = \frac{\pi(\tilde{b})}{\pi(b)} \prod_{l=1}^{L} \frac{\mathbb{P}(P_l|\tilde{b})}{\mathbb{P}(P_l|b)}$$

Calculating the logarithm of $h$ we obtain

$$\log h = \log \frac{\pi(\tilde{b})}{\pi(b)} + L \cdot \log \frac{\Gamma(a+\tilde{b}) \cdot \Gamma(b)}{\Gamma(\tilde{b}) \cdot \Gamma(a+b)} + \sum_{l=1}^{L} (\tilde{b} - b) \log(1 - P_l)$$

#### Supplemental Figures

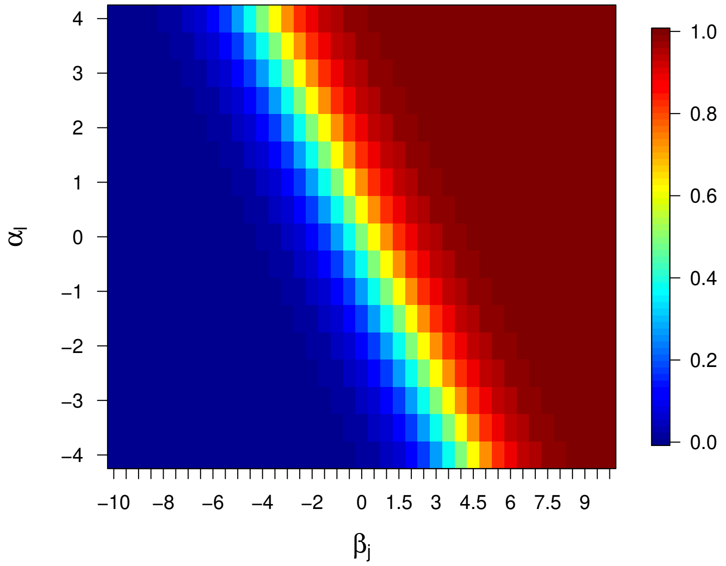


**Figure 1** Plot of the $F_{ST}^{lj}$ values as a function of $\alpha_l$ and $\beta_j$. The values of the variables $\alpha_l$ and $\beta_j$ are included in the following intervals: $\alpha_l \in [-4;4]$ and $\beta_j \in [-10;10]$. The $F_{ST}^{lj}$ values are calculated with steps of 0.5 in the $\alpha_l$ and $\beta_j$ range.

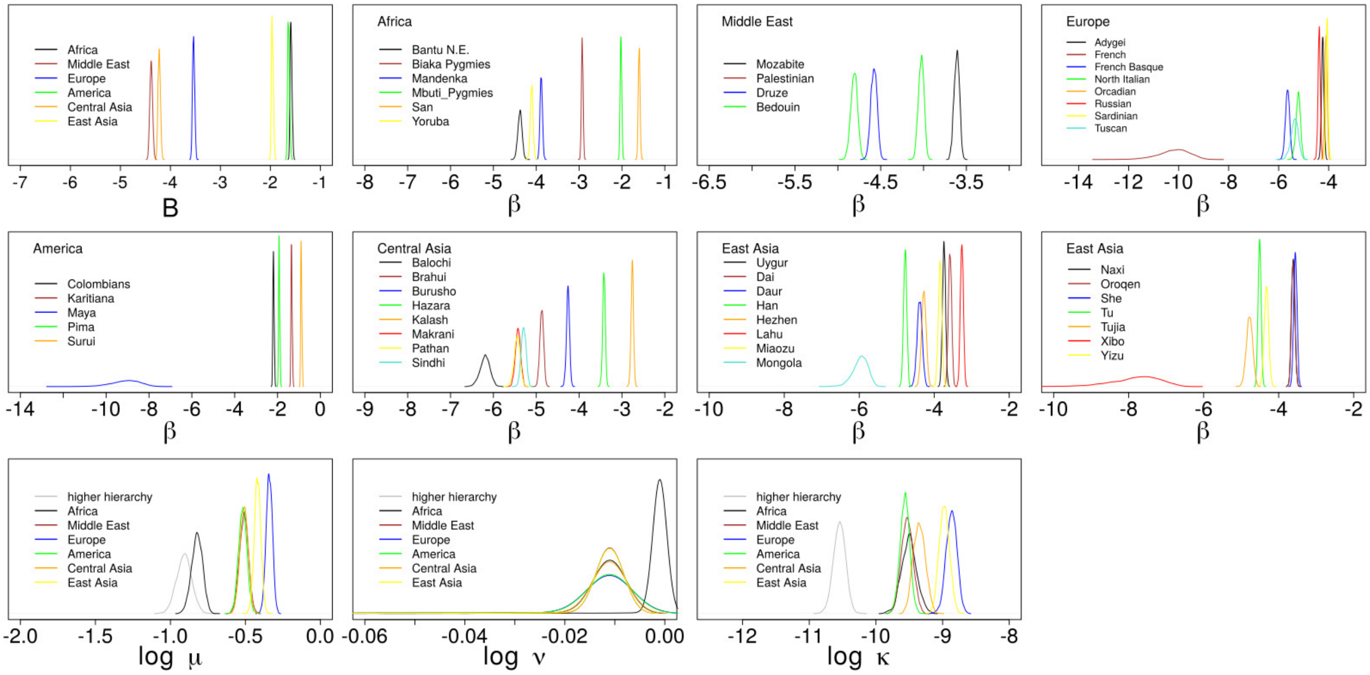


**Figure 2** Plot of the posterior distributions of the parameters inferred by Flink of the $p$ arm of the first chromosome.

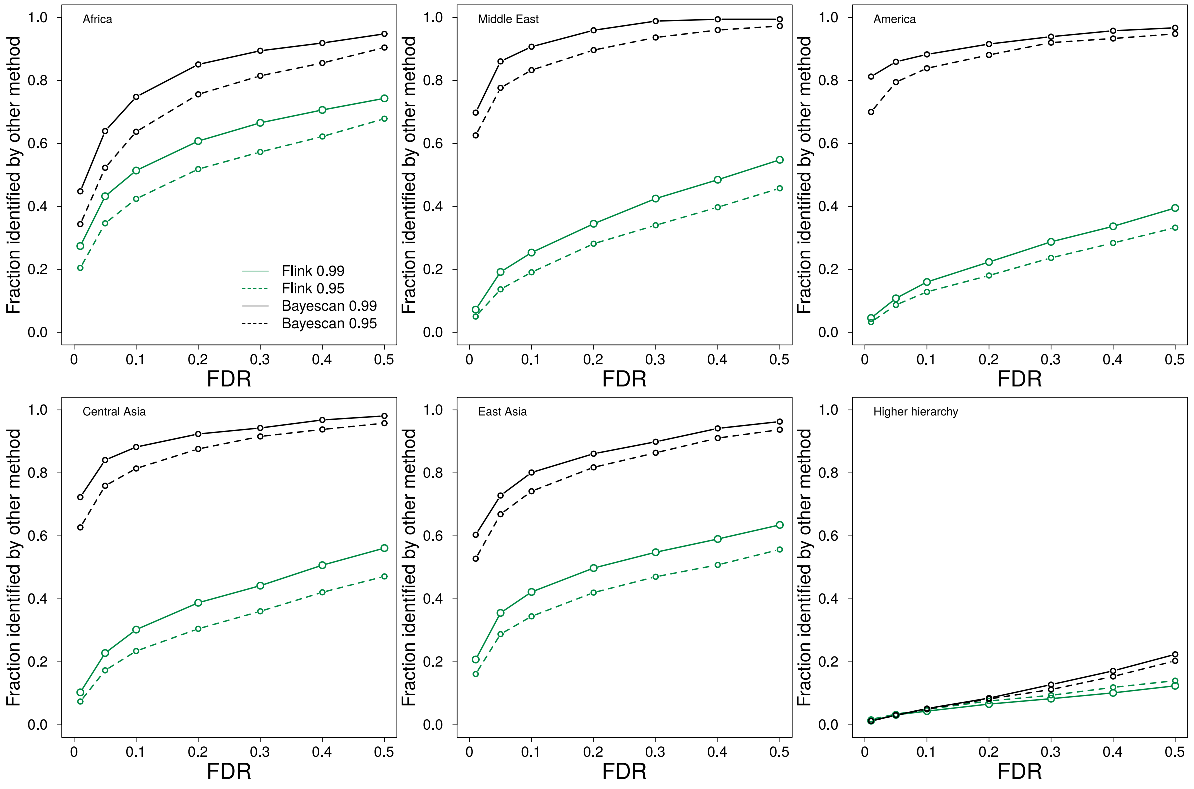


**Figure 3** The fraction of regions identified as divergent among the different groups by `Flink` (green) and `Bayescan` (black) at a false discovery rate (FDR) of 0.01 (solid) and 0.05 (dashed) also identified by the other method at different FDR.

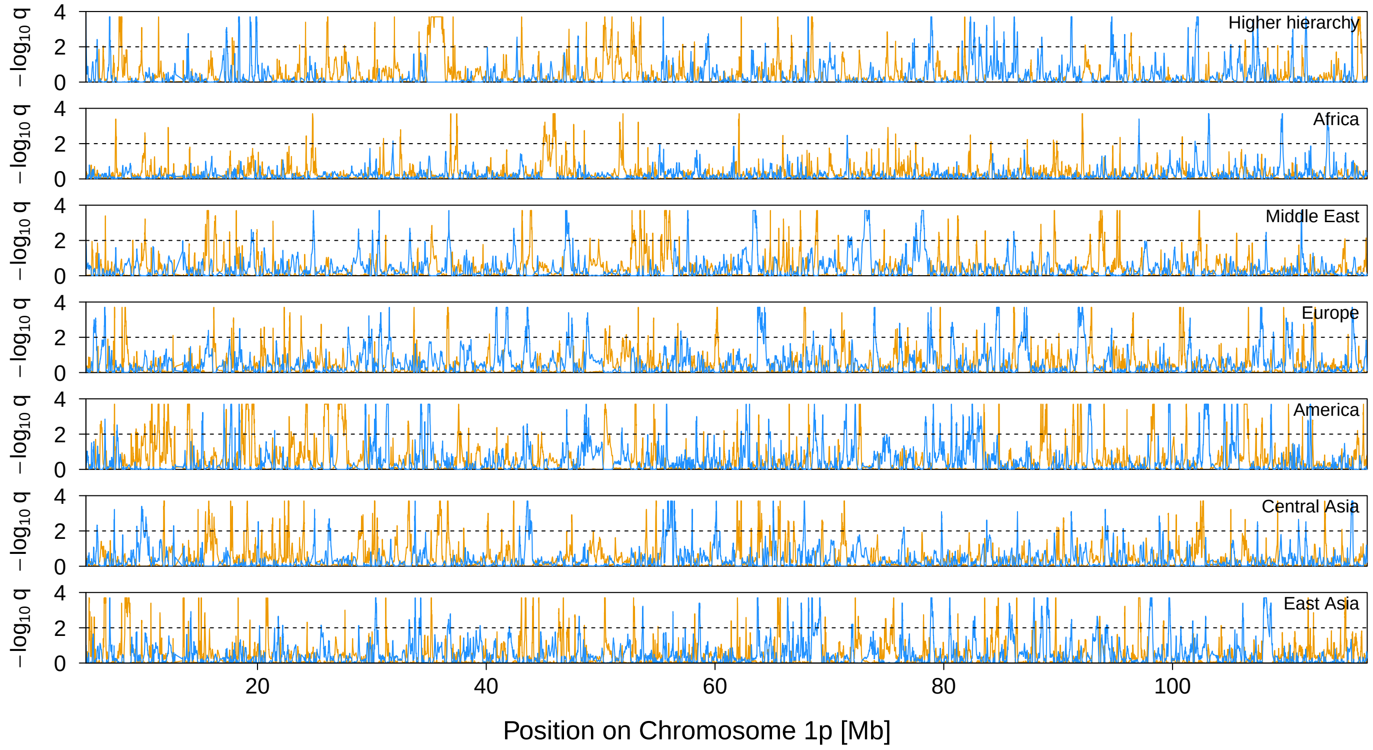


**Figure 4** Signal of selection on Chromosome 1p. The orange and blue lines indicate the locus-specific FDR for divergent (orange) and balancing (blue) selection, respectively. The black dashed line shows the 1% FDR threshold.

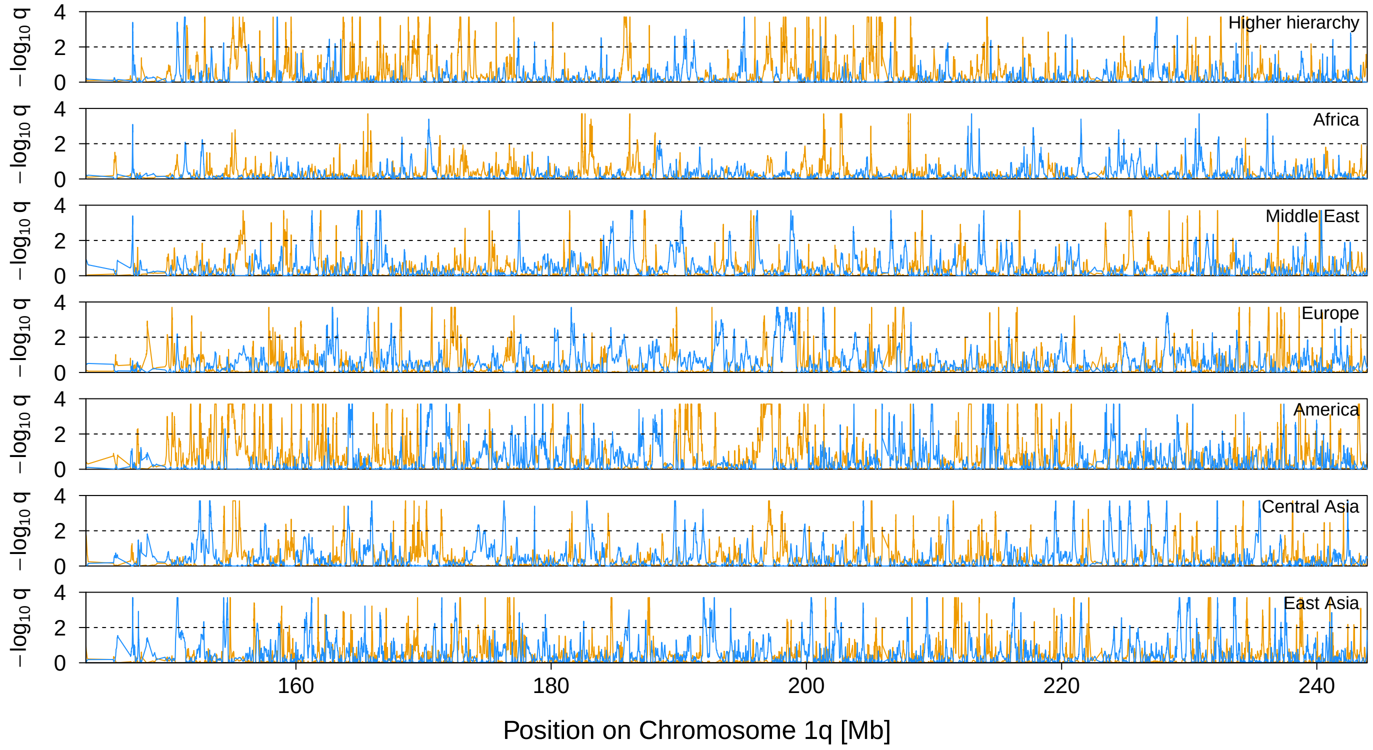


**Figure 5** Signal of selection on Chromosome 1q. The orange and blue lines indicate the locus-specific FDR for divergent (orange) and balancing (blue) selection, respectively. The black dashed line shows the 1% FDR threshold.

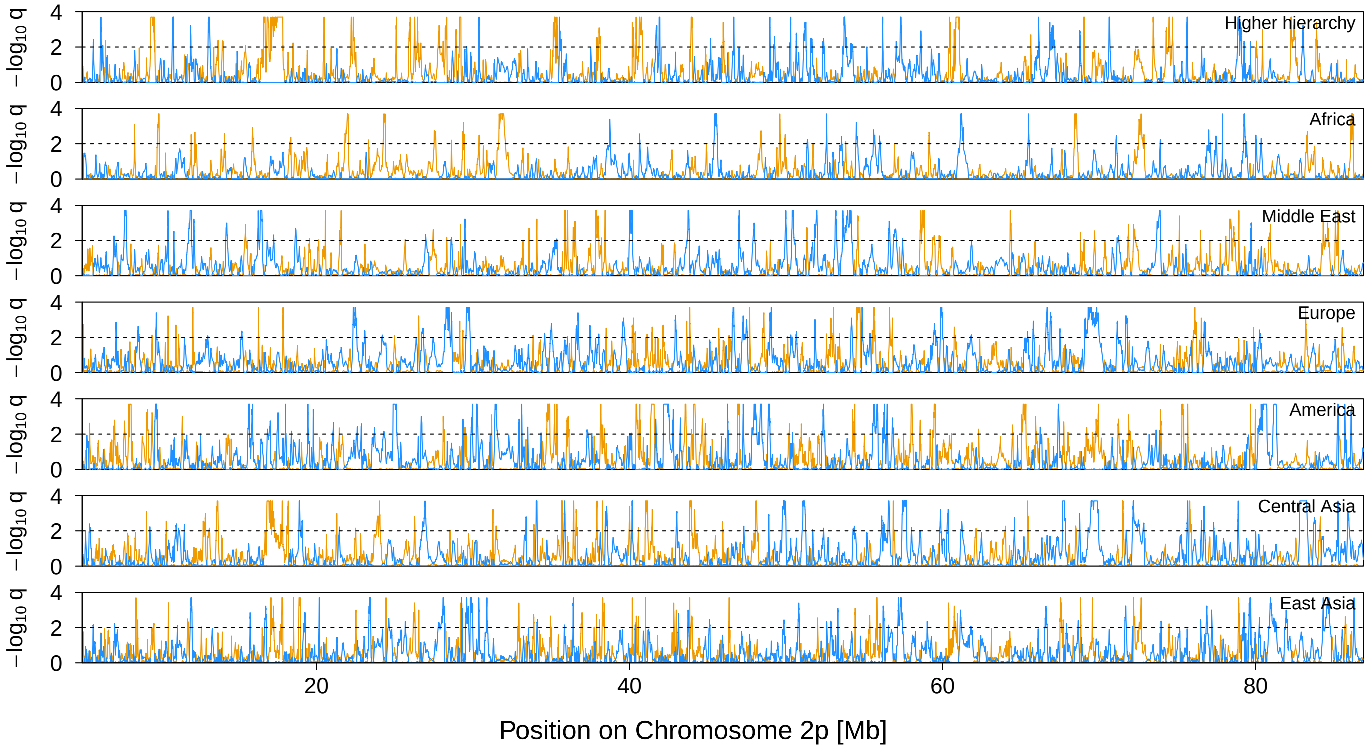


**Figure 6** Signal of selection on Chromosome 2p. The orange and blue lines indicate the locus-specific FDR for divergent (orange) and balancing (blue) selection, respectively. The black dashed line shows the 1% FDR threshold.

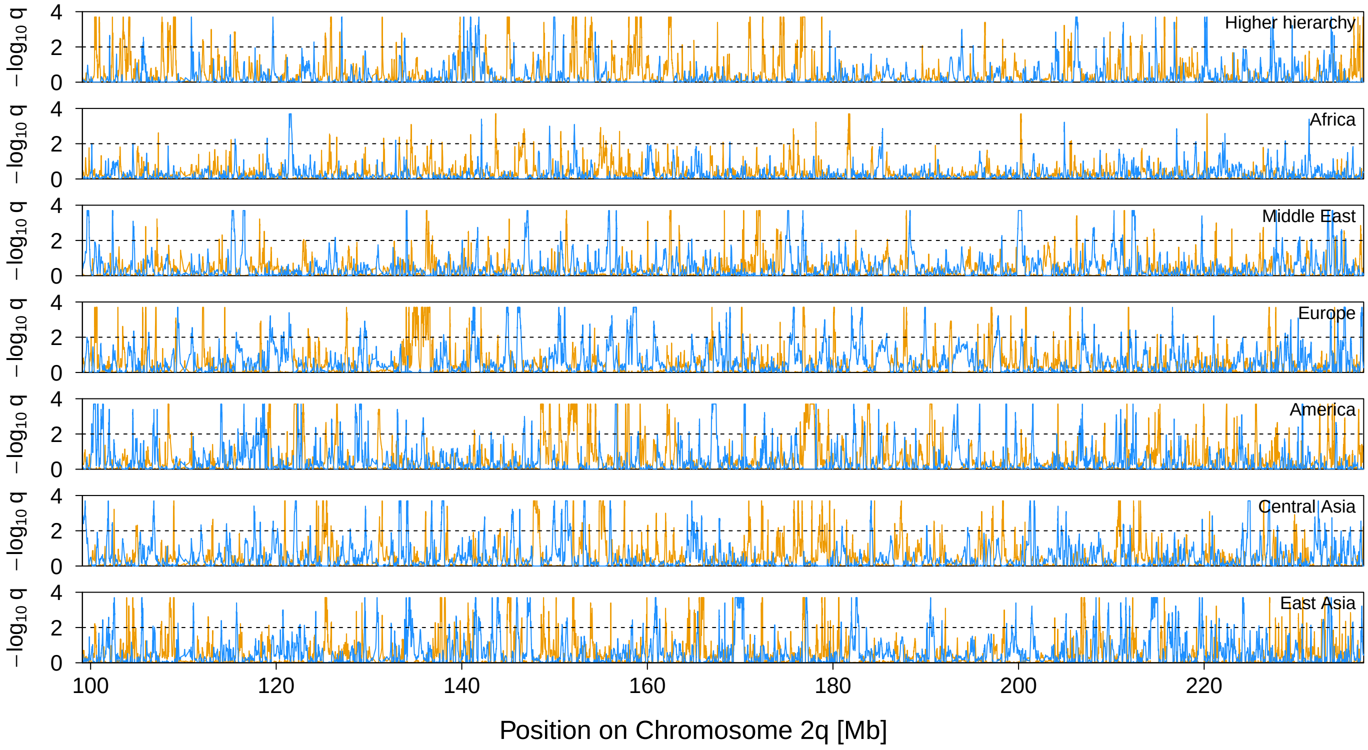


**Figure 7** Signal of selection on Chromosome 2q. The orange and blue lines indicate the locus-specific FDR for divergent (orange) and balancing (blue) selection, respectively. The black dashed line shows the 1% FDR threshold.

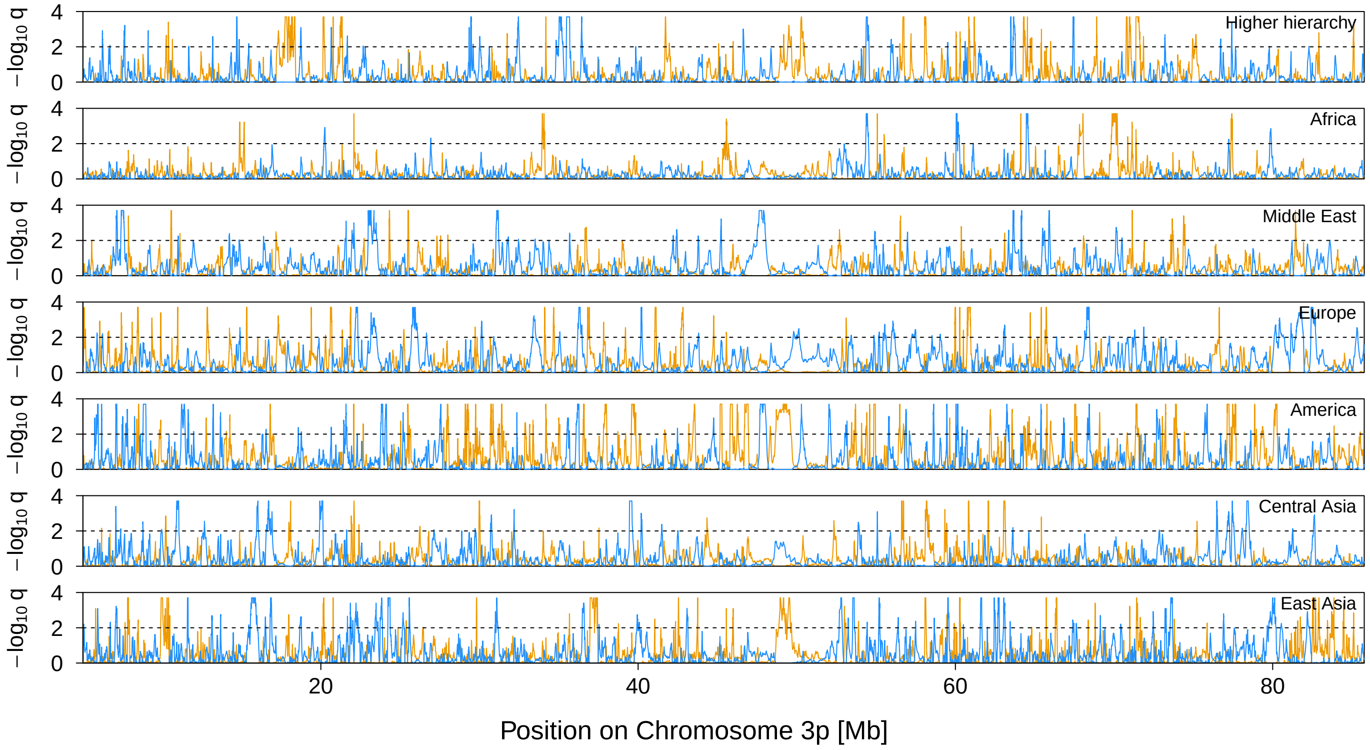


**Figure 8** Signal of selection on Chromosome 3p. The orange and blue lines indicate the locus-specific FDR for divergent (orange) and balancing (blue) selection, respectively. The black dashed line shows the 1% FDR threshold.

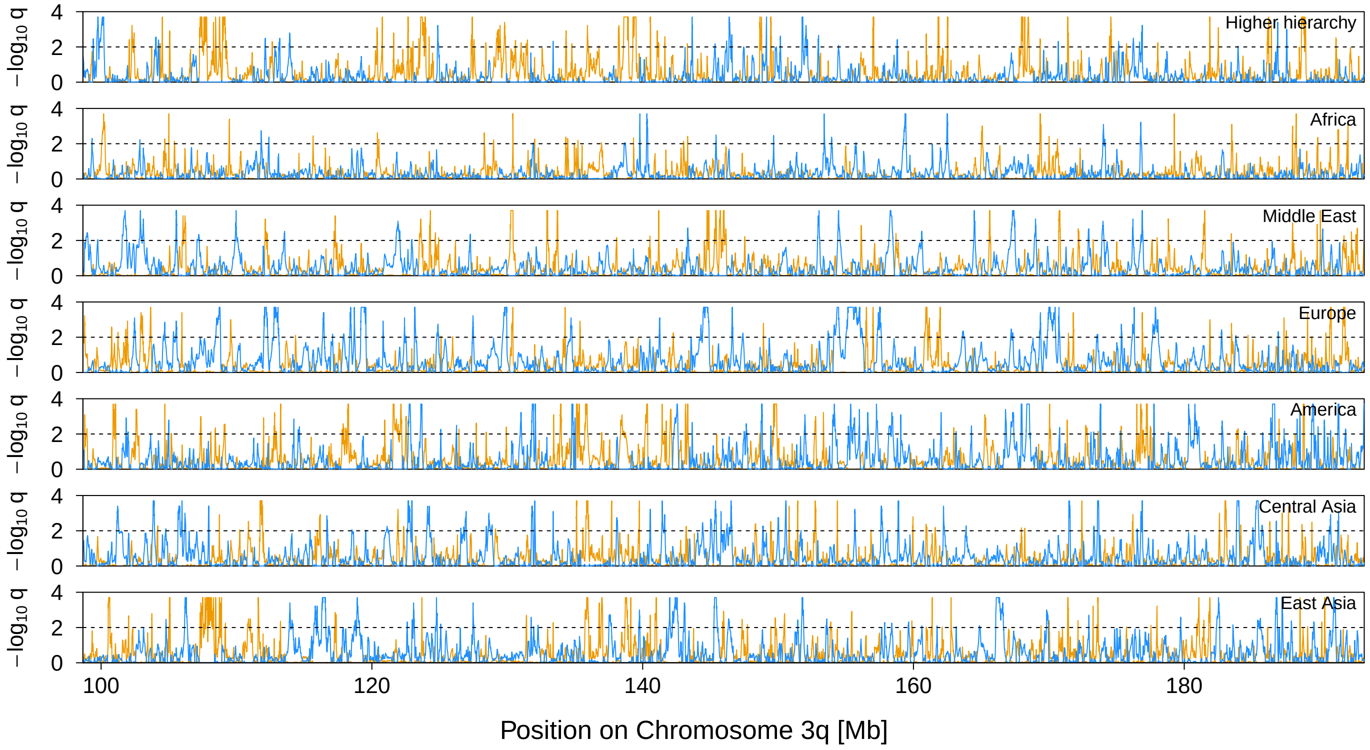


**Figure 9** Signal of selection on Chromosome 3q. The orange and blue lines indicate the locus-specific FDR for divergent (orange) and balancing (blue) selection, respectively. The black dashed line shows the 1% FDR threshold.

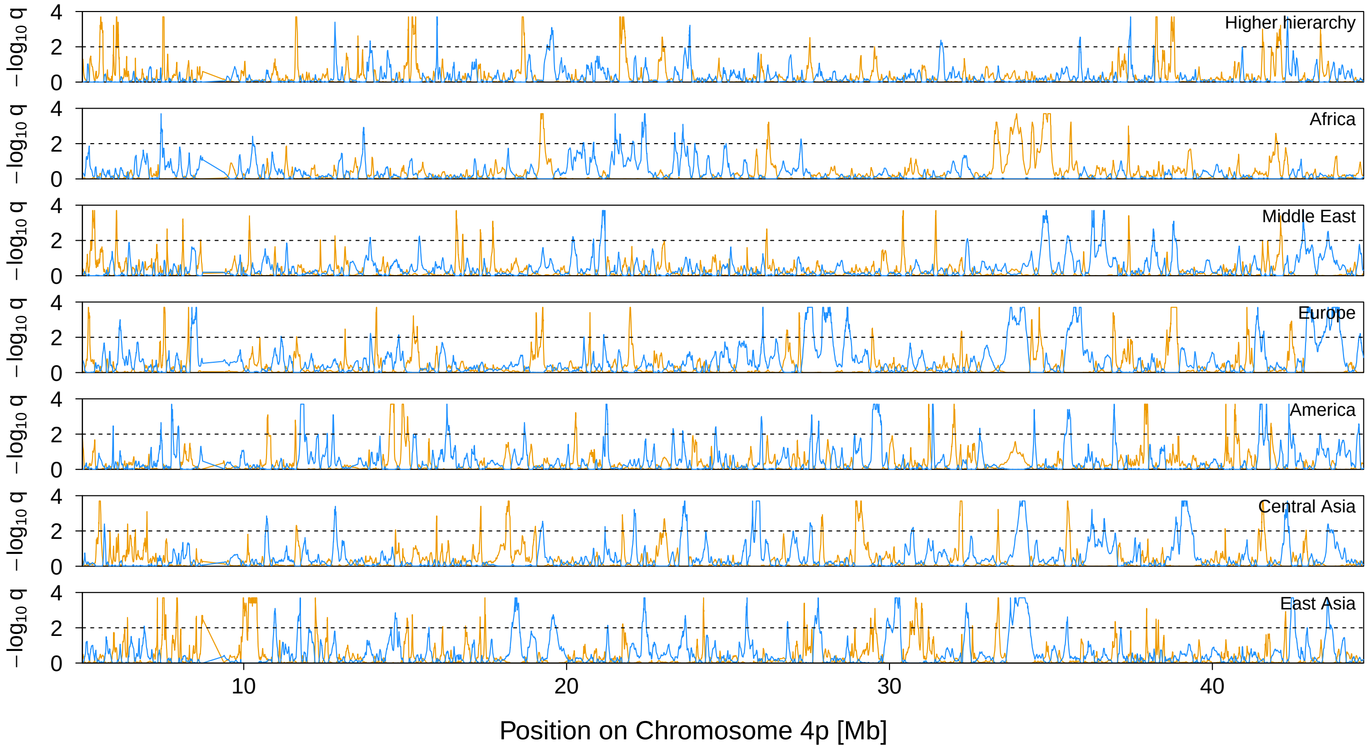


**Figure 10** Signal of selection on Chromosome 4p. The orange and blue lines indicate the locus-specific FDR for divergent (orange) and balancing (blue) selection, respectively. The black dashed line shows the 1% FDR threshold.

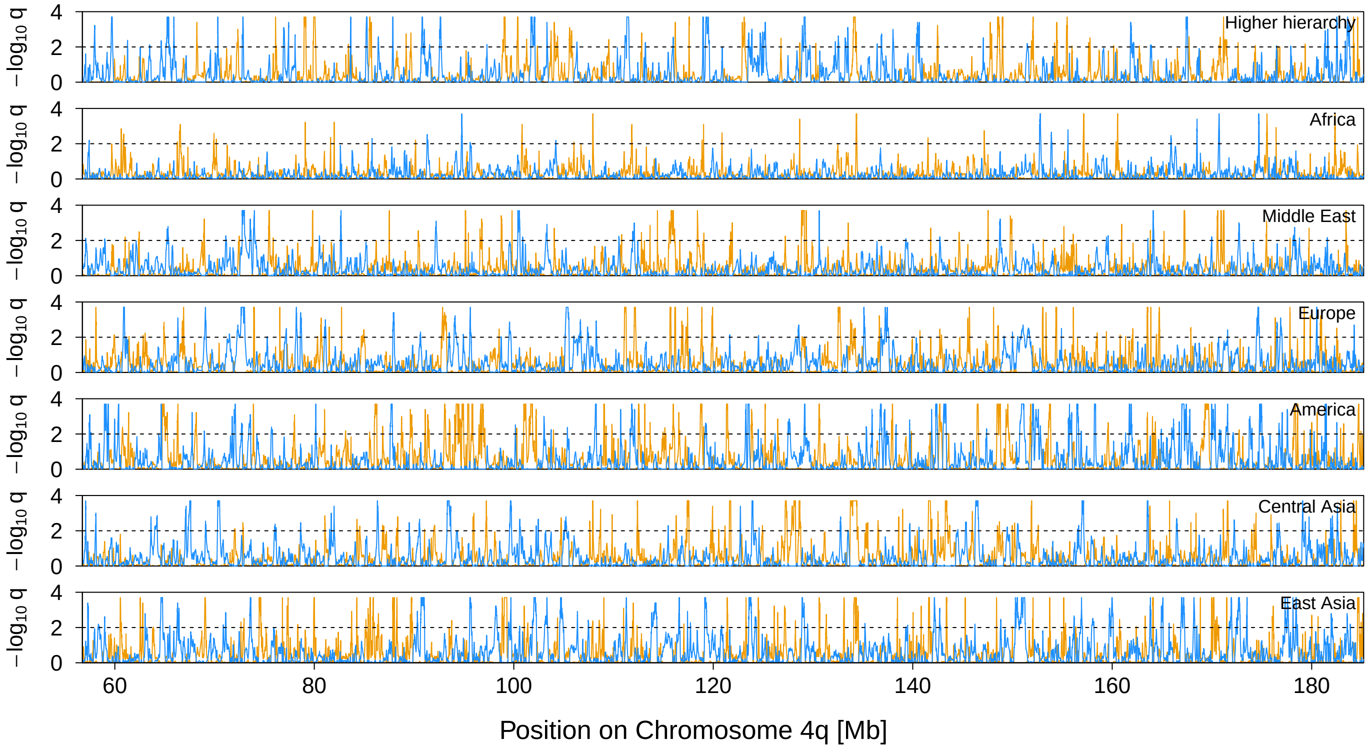


**Figure 11** Signal of selection on Chromosome 4q. The orange and blue lines indicate the locus-specific FDR for divergent (orange) and balancing (blue) selection, respectively. The black dashed line shows the 1% FDR threshold.

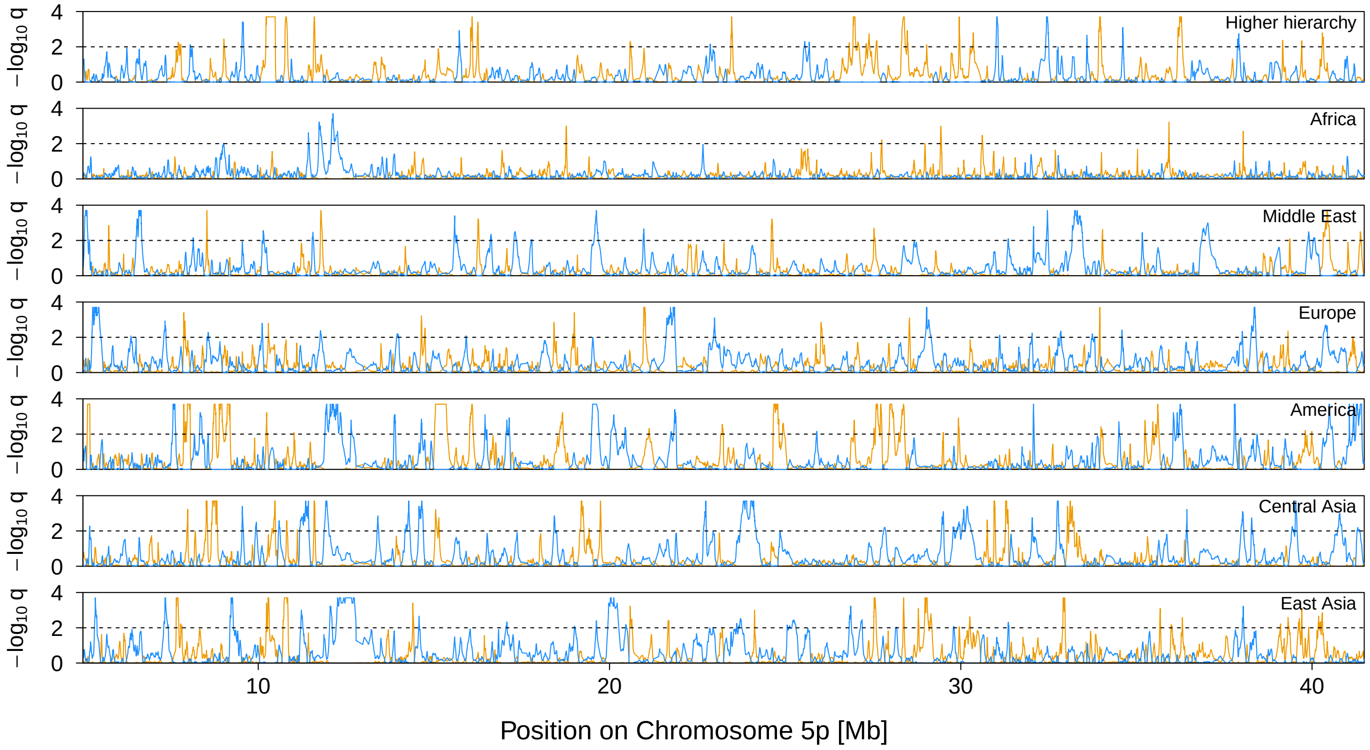


**Figure 12** Signal of selection on Chromosome 5p. The orange and blue lines indicate the locus-specific FDR for divergent (orange) and balancing (blue) selection, respectively. The black dashed line shows the 1% FDR threshold.

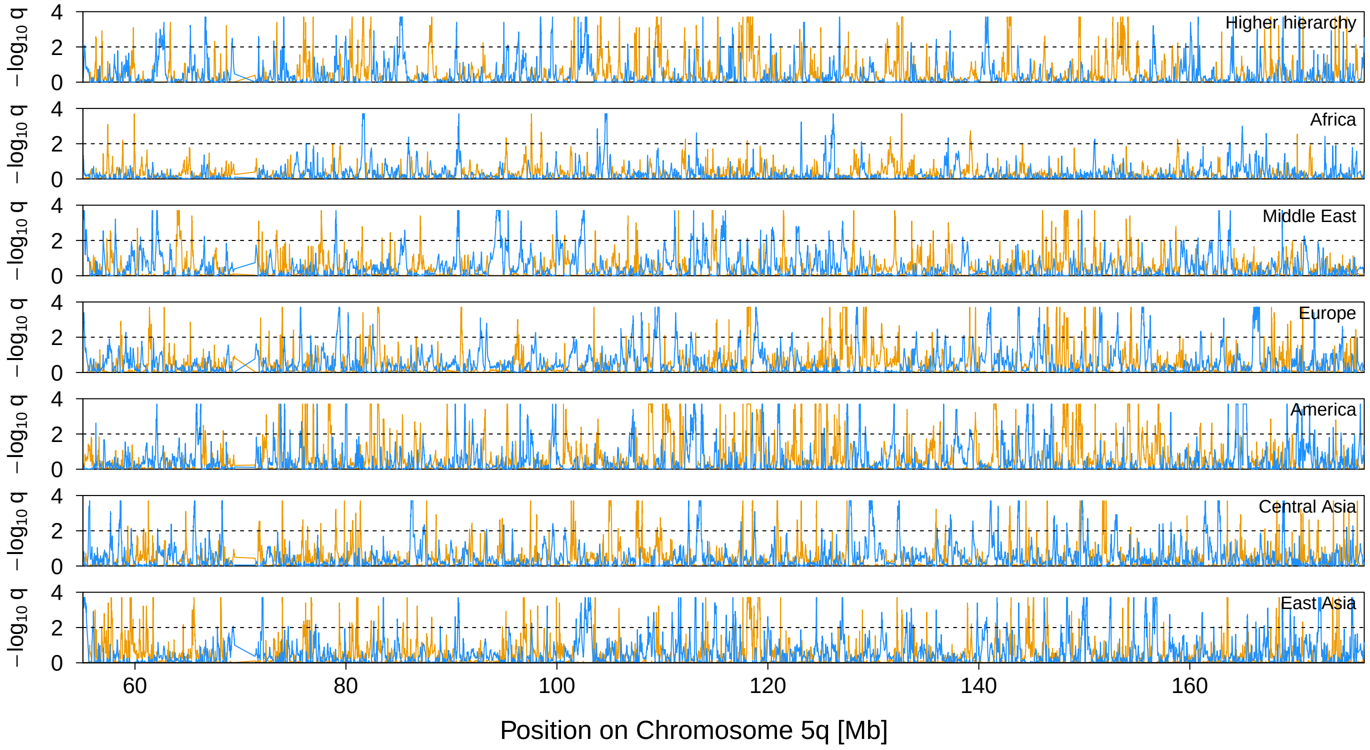


**Figure 13** Signal of selection on Chromosome 5q. The orange and blue lines indicate the locus-specific FDR for divergent (orange) and balancing (blue) selection, respectively. The black dashed line shows the 1% FDR threshold.

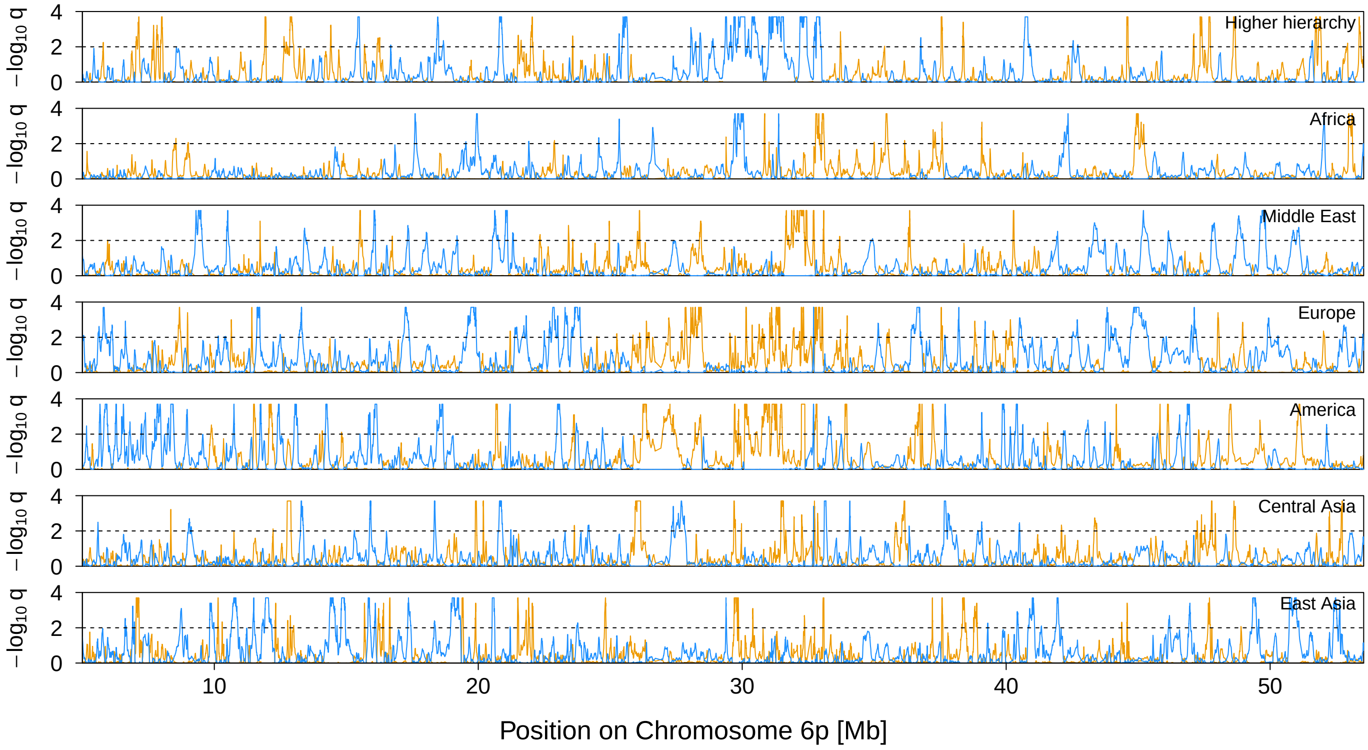


**Figure 14** Signal of selection on Chromosome 6p. The orange and blue lines indicate the locus-specific FDR for divergent (orange) and balancing (blue) selection, respectively. The black dashed line shows the 1% FDR threshold.

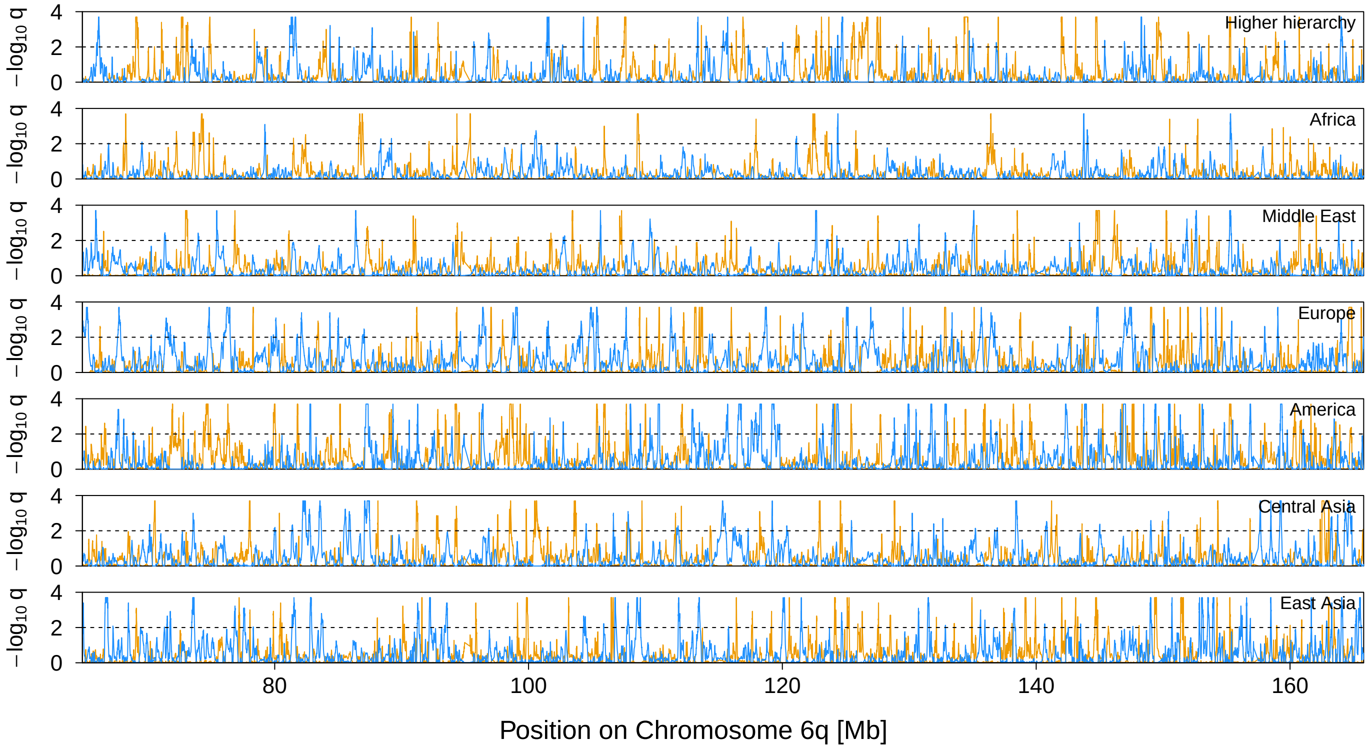


**Figure 15** Signal of selection on Chromosome 6q. The orange and blue lines indicate the locus-specific FDR for divergent (orange) and balancing (blue) selection, respectively. The black dashed line shows the 1% FDR threshold.

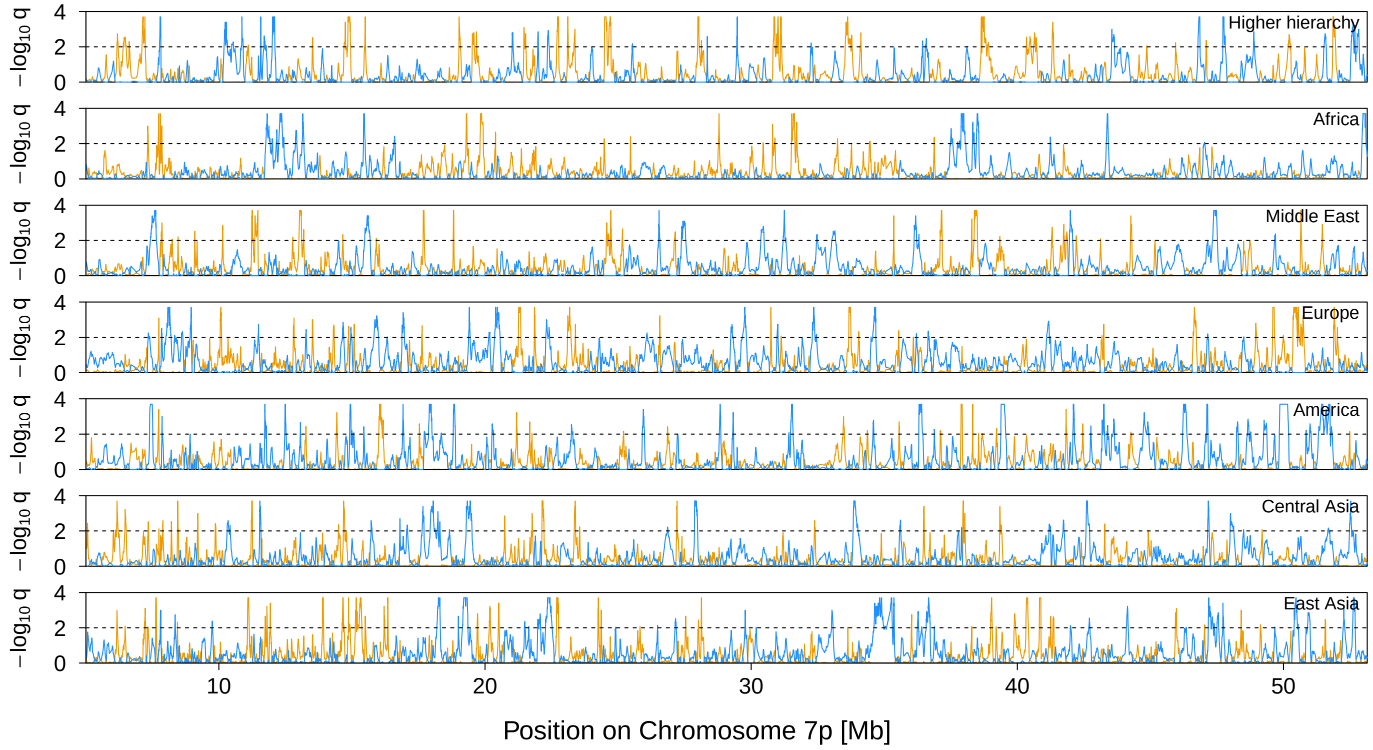


**Figure 16** Signal of selection on Chromosome 7p. The orange and blue lines indicate the locus-specific FDR for divergent (orange) and balancing (blue) selection, respectively. The black dashed line shows the 1% FDR threshold.

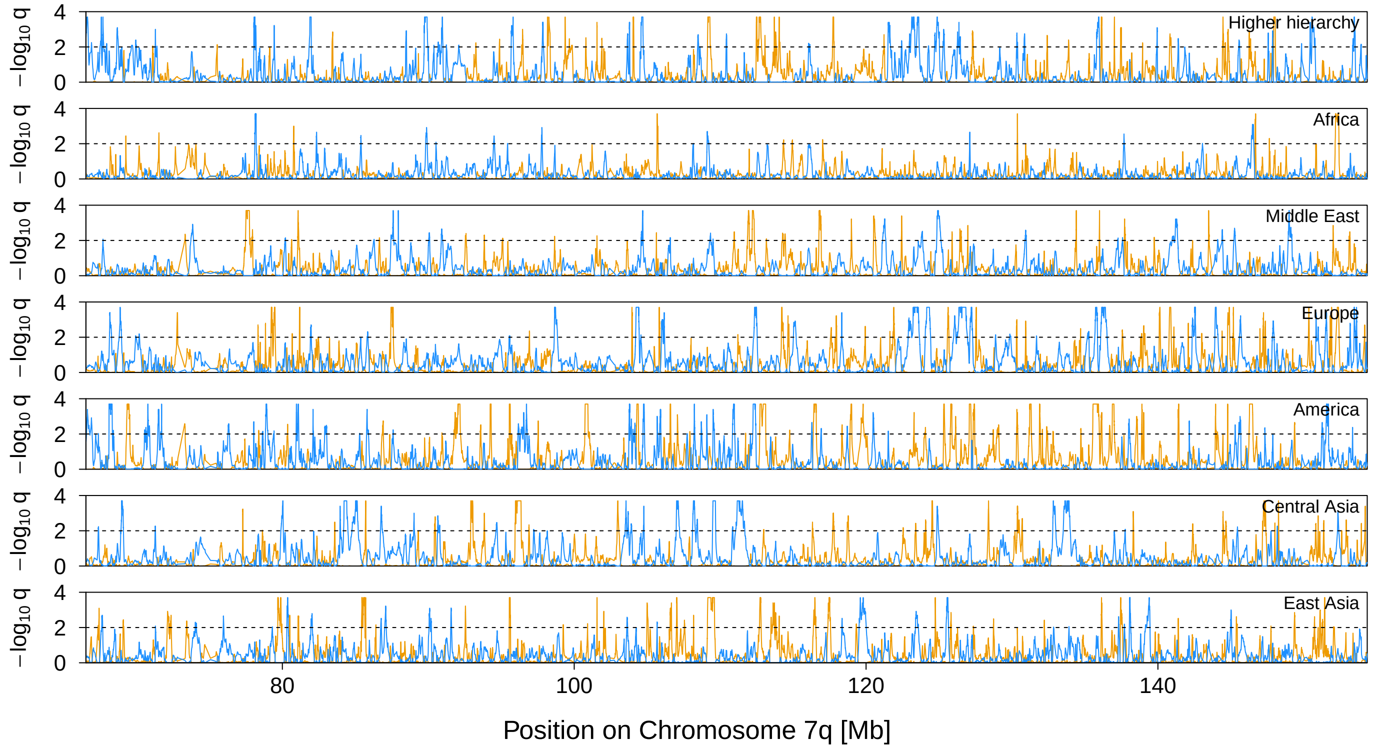


**Figure 17** Signal of selection on Chromosome 7q. The orange and blue lines indicate the locus-specific FDR for divergent (orange) and balancing (blue) selection, respectively. The black dashed line shows the 1% FDR threshold.

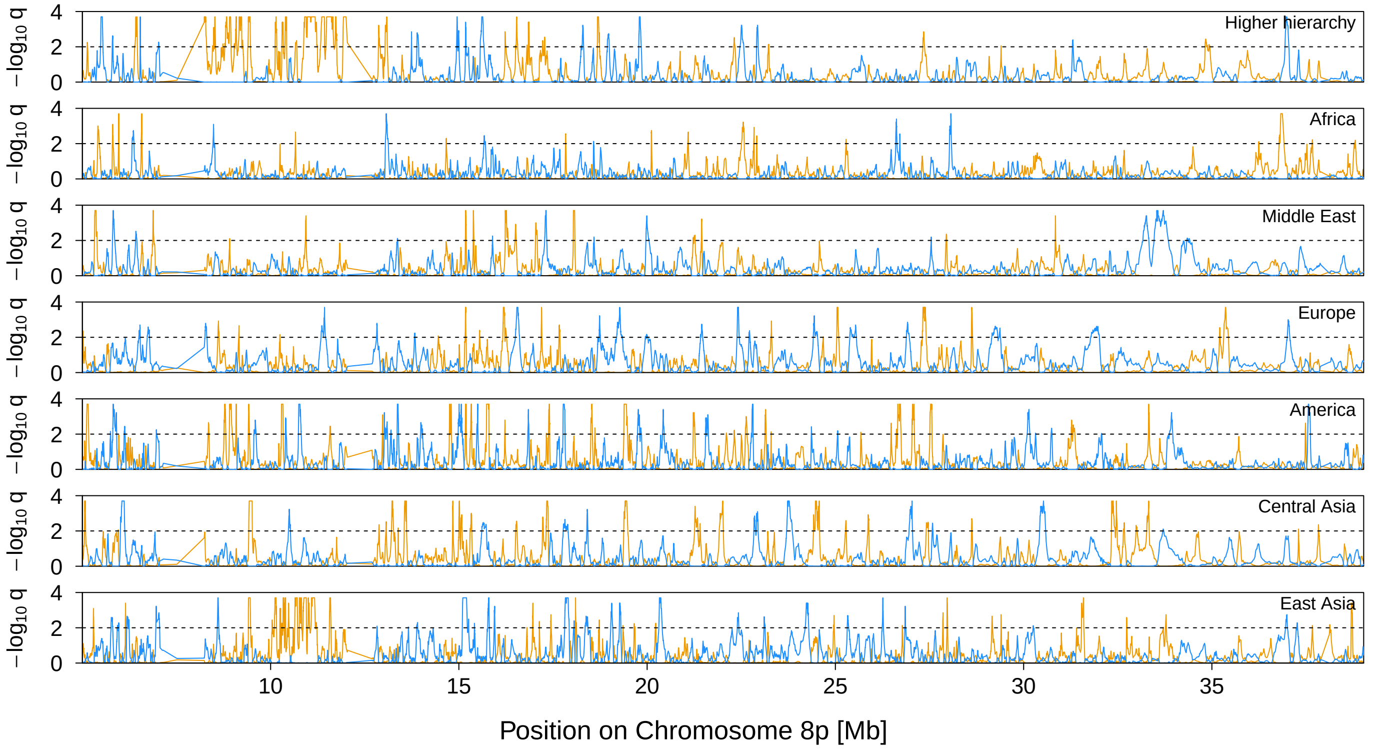


**Figure 18** Signal of selection on Chromosome 8p. The orange and blue lines indicate the locus-specific FDR for divergent (orange) and balancing (blue) selection, respectively. The black dashed line shows the 1% FDR threshold.

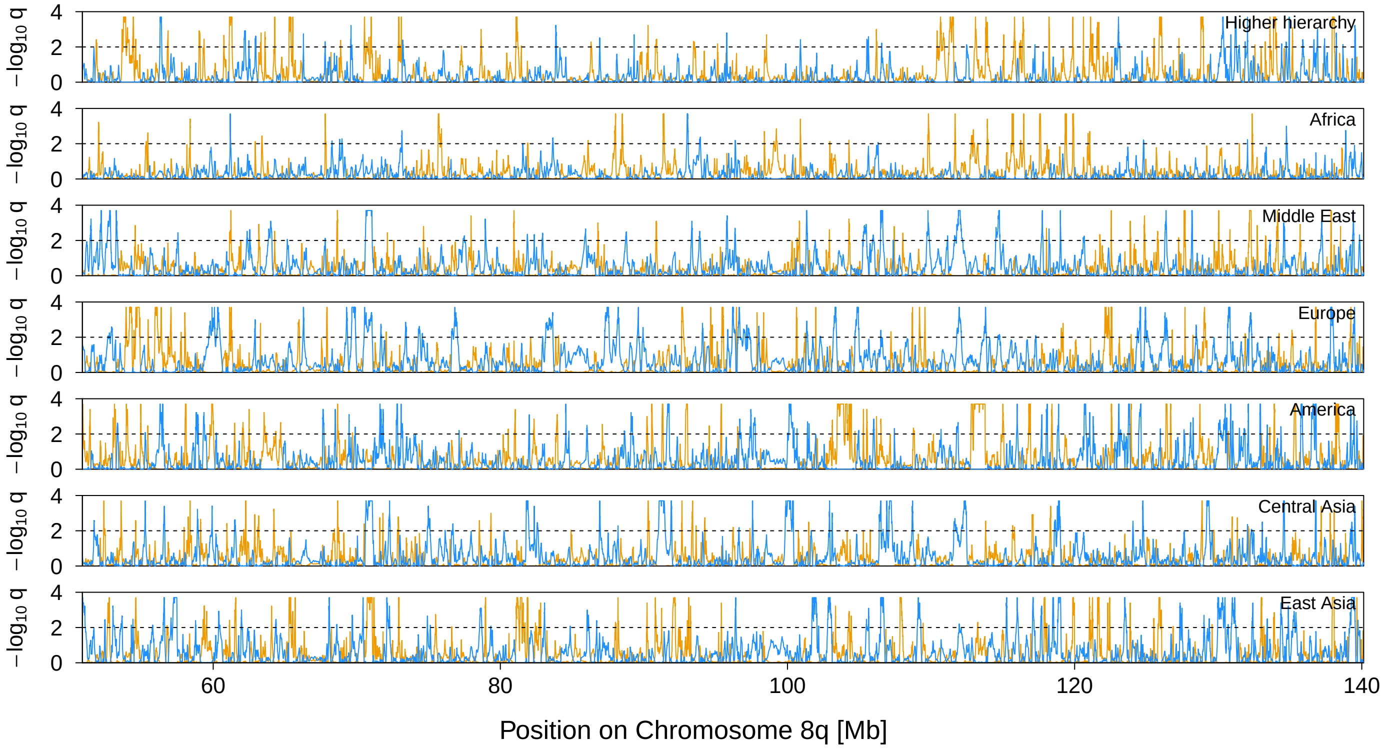


**Figure 19** Signal of selection on Chromosome 8q. The orange and blue lines indicate the locus-specific FDR for divergent (orange) and balancing (blue) selection, respectively. The black dashed line shows the 1% FDR threshold.

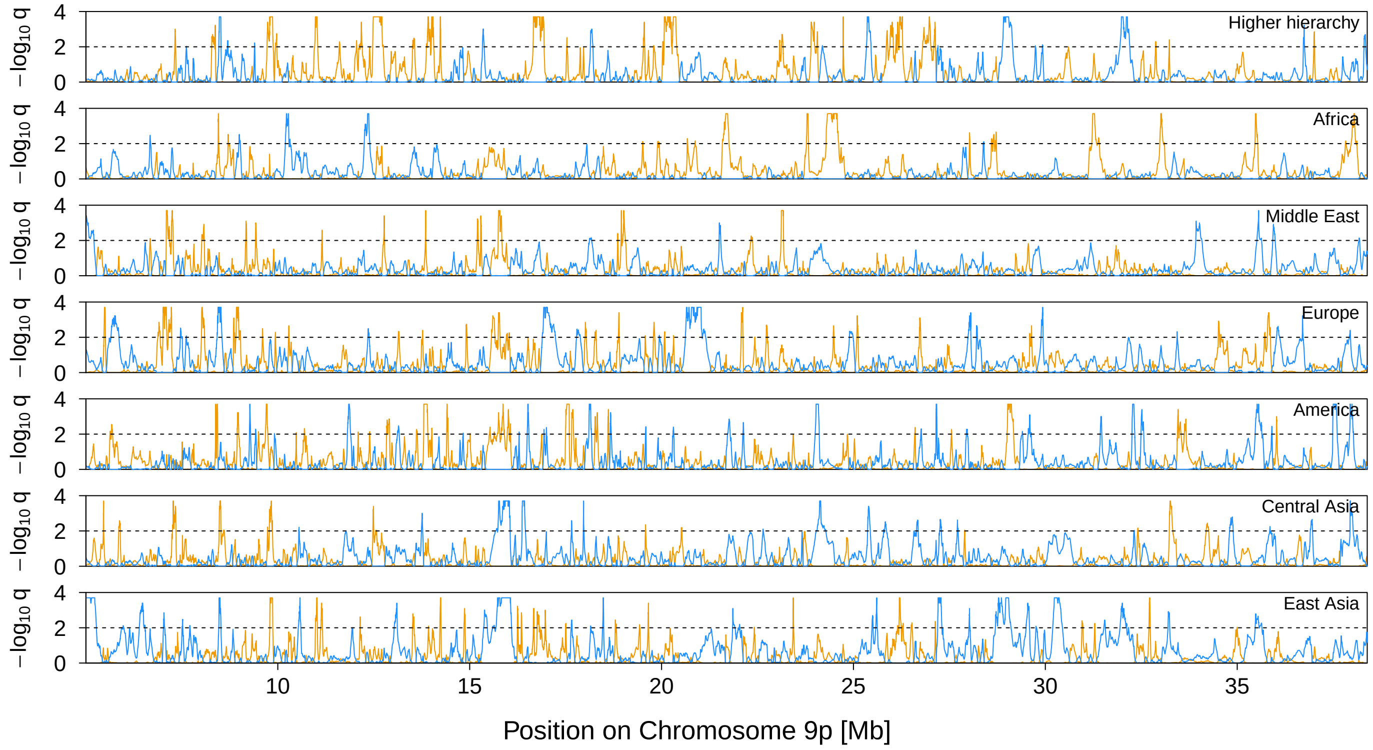


**Figure 20** Signal of selection on Chromosome 9p. The orange and blue lines indicate the locus-specific FDR for divergent (orange) and balancing (blue) selection, respectively. The black dashed line shows the 1% FDR threshold.

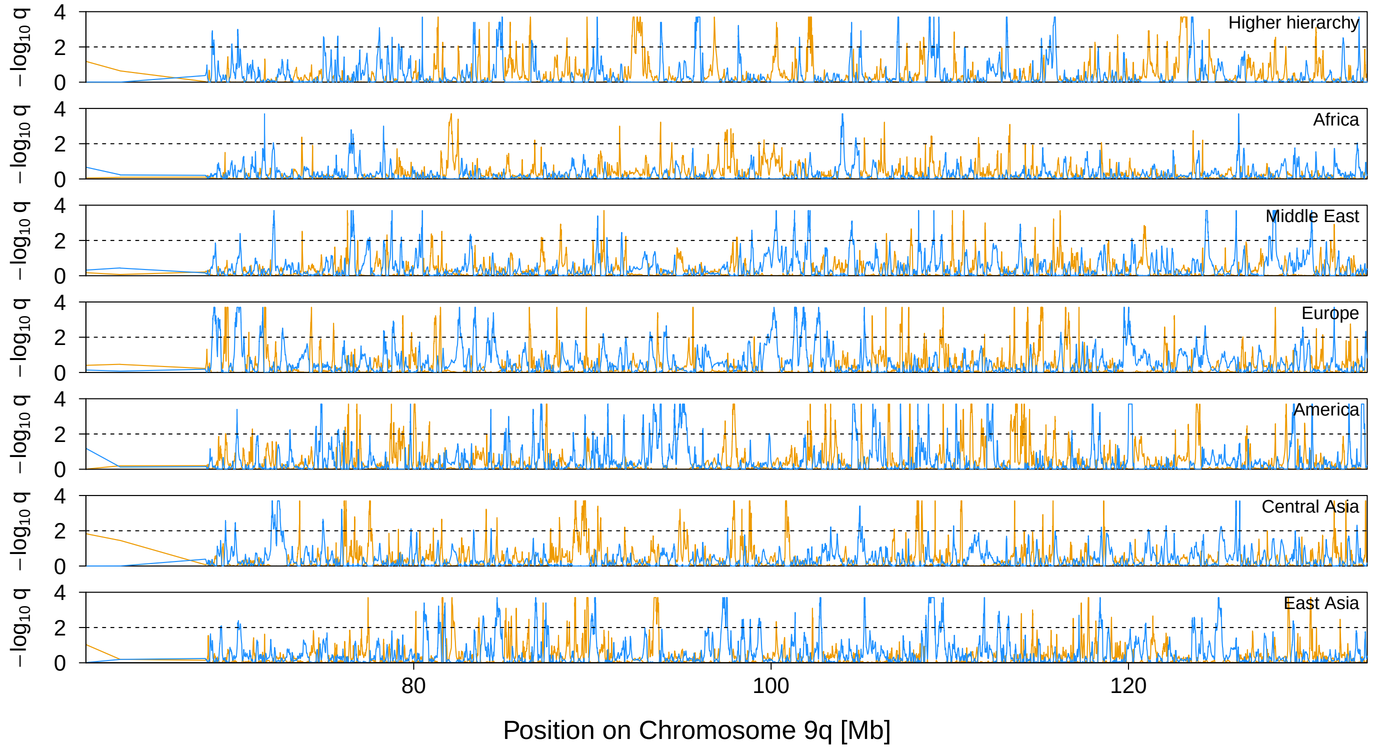


**Figure 21** Signal of selection on Chromosome 9q. The orange and blue lines indicate the locus-specific FDR for divergent (orange) and balancing (blue) selection, respectively. The black dashed line shows the 1% FDR threshold.

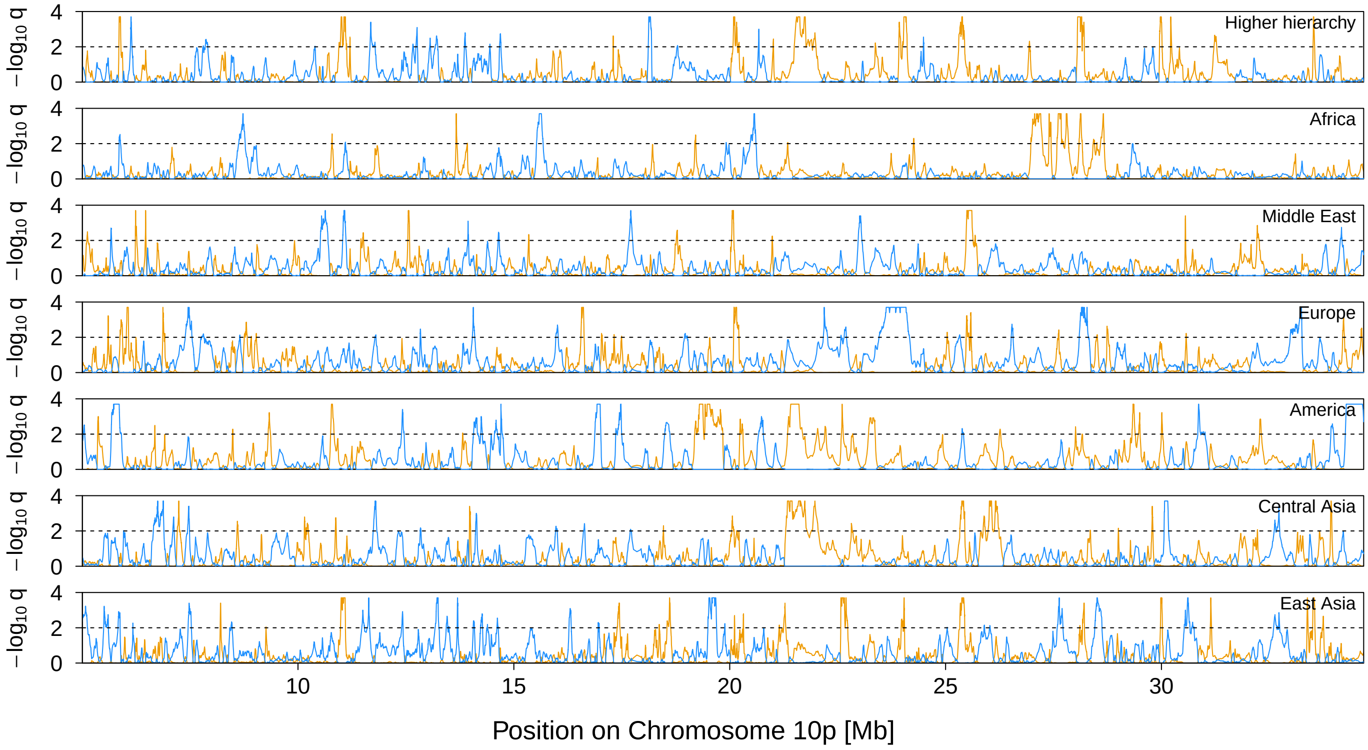


**Figure 22** Signal of selection on Chromosome 10p. The orange and blue lines indicate the locus-specific FDR for divergent (orange) and balancing (blue) selection, respectively. The black dashed line shows the 1% FDR threshold.

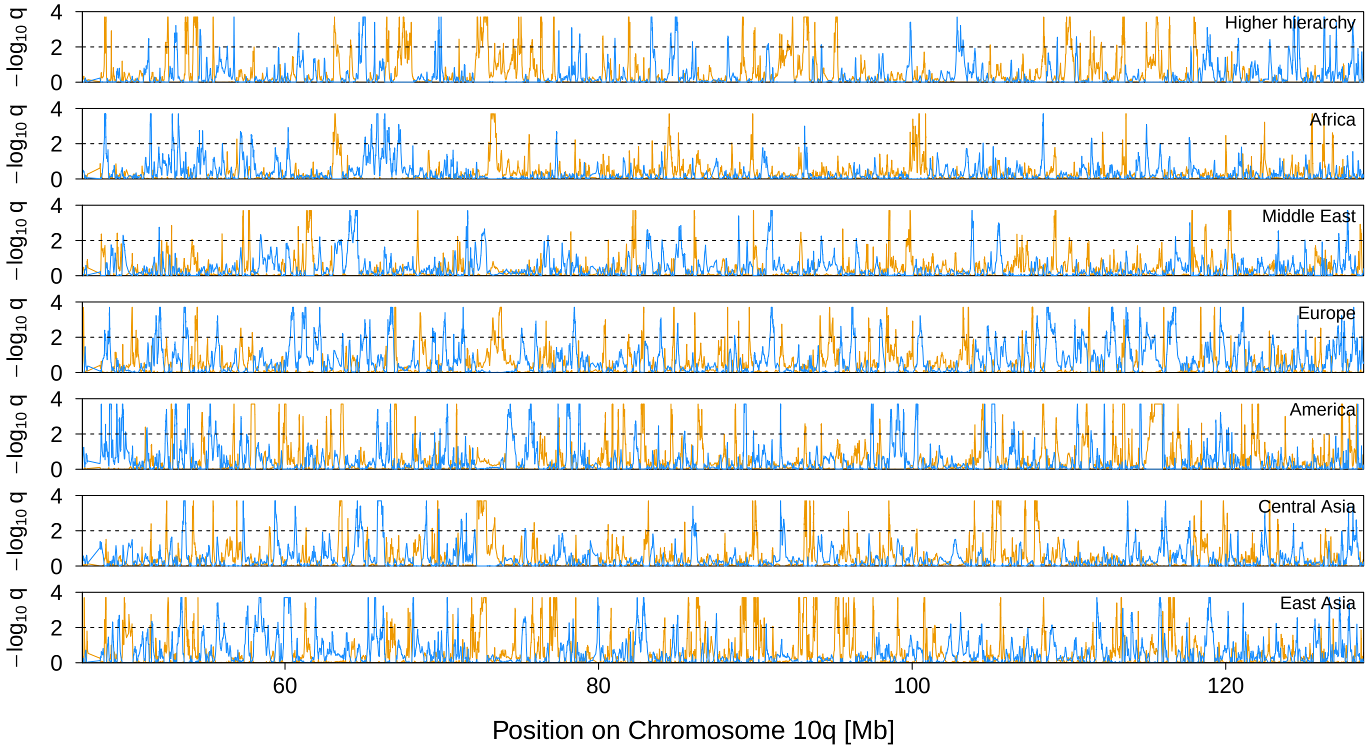


**Figure 23** Signal of selection on Chromosome 10q. The orange and blue lines indicate the locus-specific FDR for divergent (orange) and balancing (blue) selection, respectively. The black dashed line shows the 1% FDR threshold.

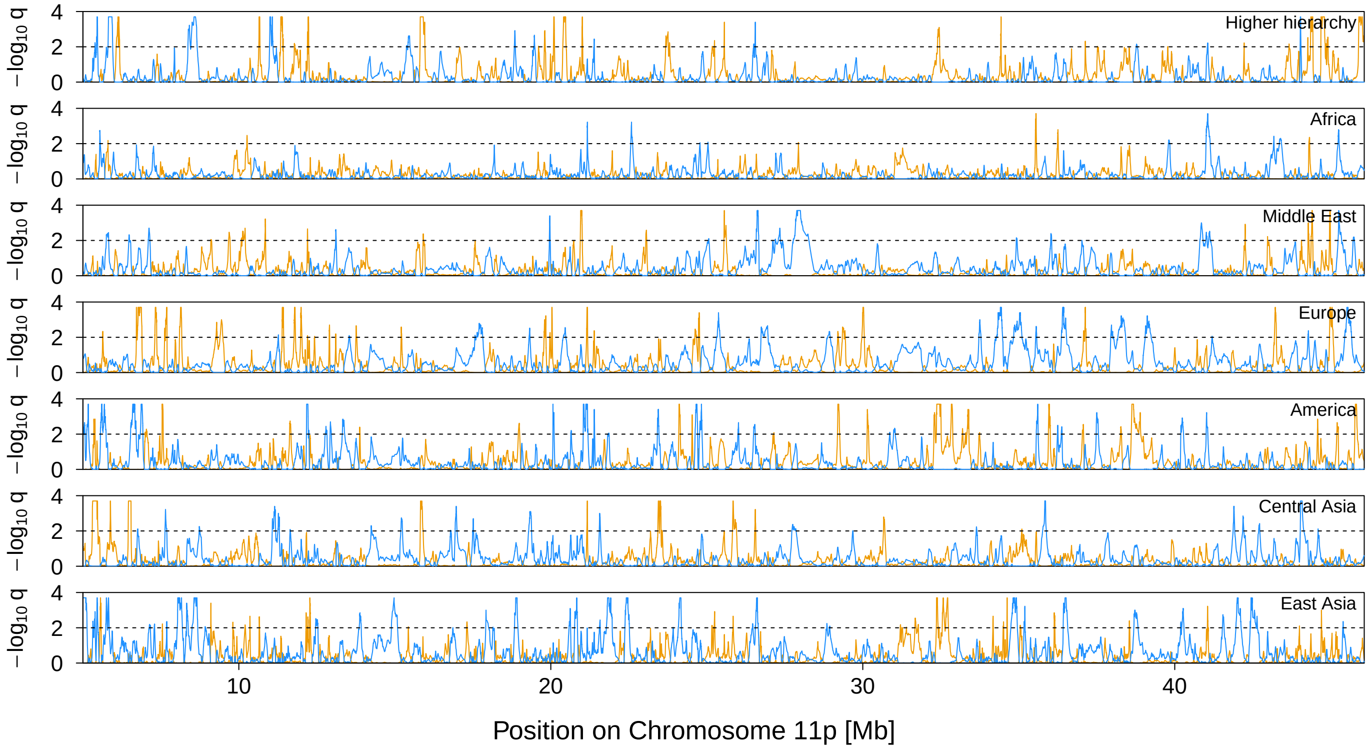


**Figure 24** Signal of selection on Chromosome 11p. The orange and blue lines indicate the locus-specific FDR for divergent (orange) and balancing (blue) selection, respectively. The black dashed line shows the 1% FDR threshold.

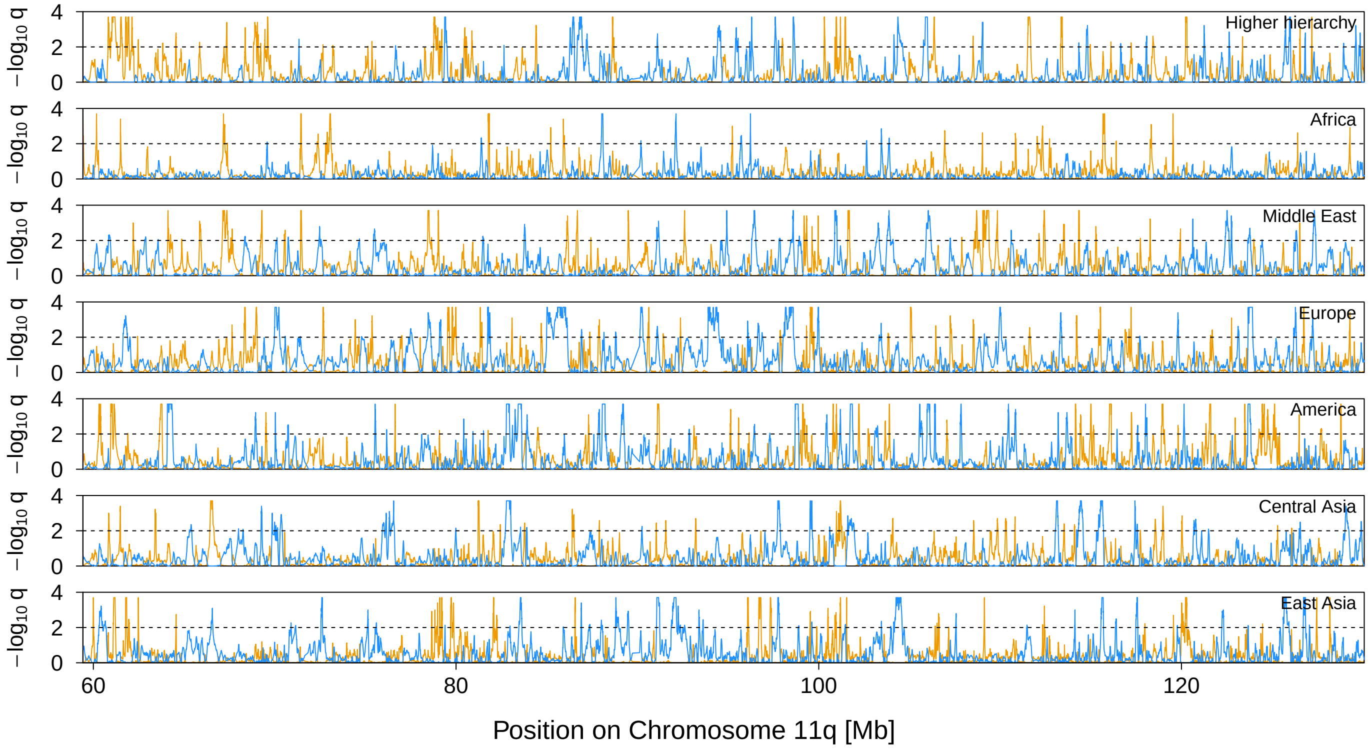


**Figure 25** Signal of selection on Chromosome 11q. The orange and blue lines indicate the locus-specific FDR for divergent (orange) and balancing (blue) selection, respectively. The black dashed line shows the 1% FDR threshold.

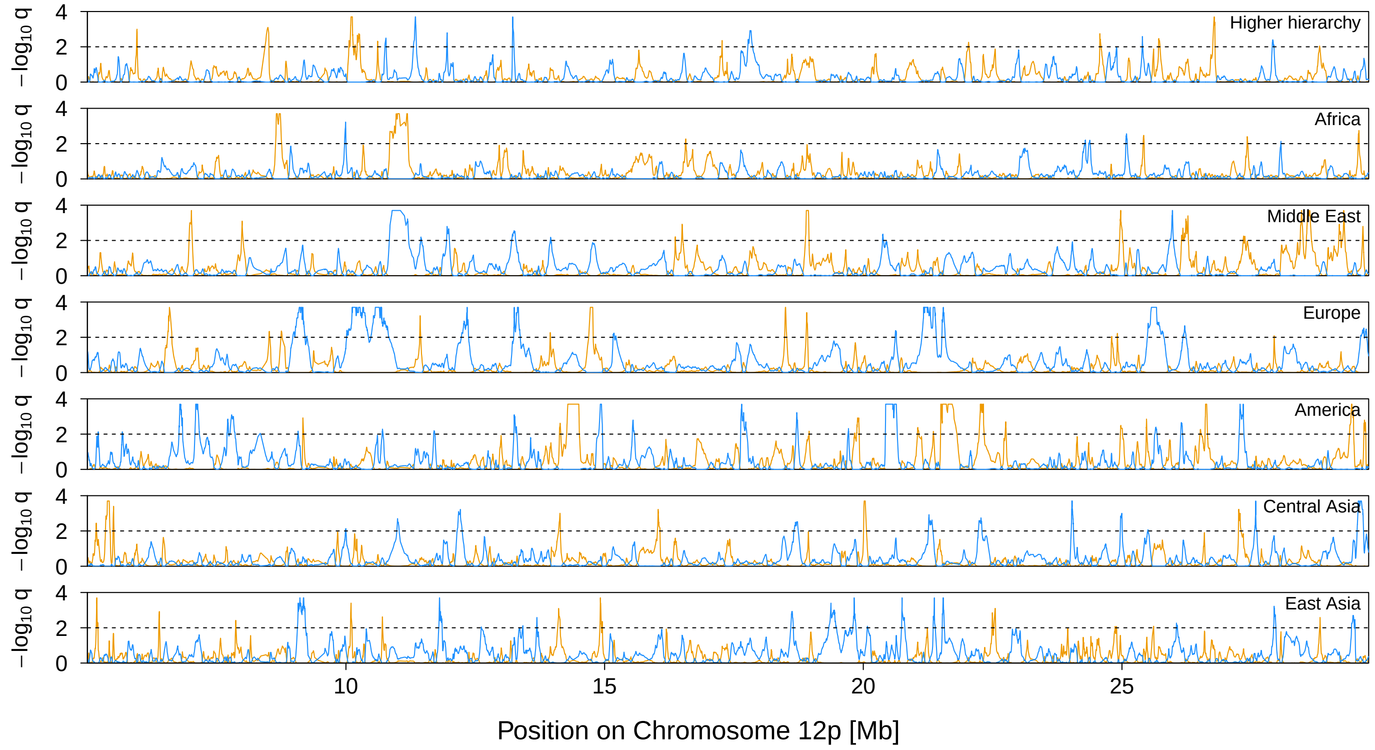


**Figure 26** Signal of selection on Chromosome 12p. The orange and blue lines indicate the locus-specific FDR for divergent (orange) and balancing (blue) selection, respectively. The black dashed line shows the 1% FDR threshold.

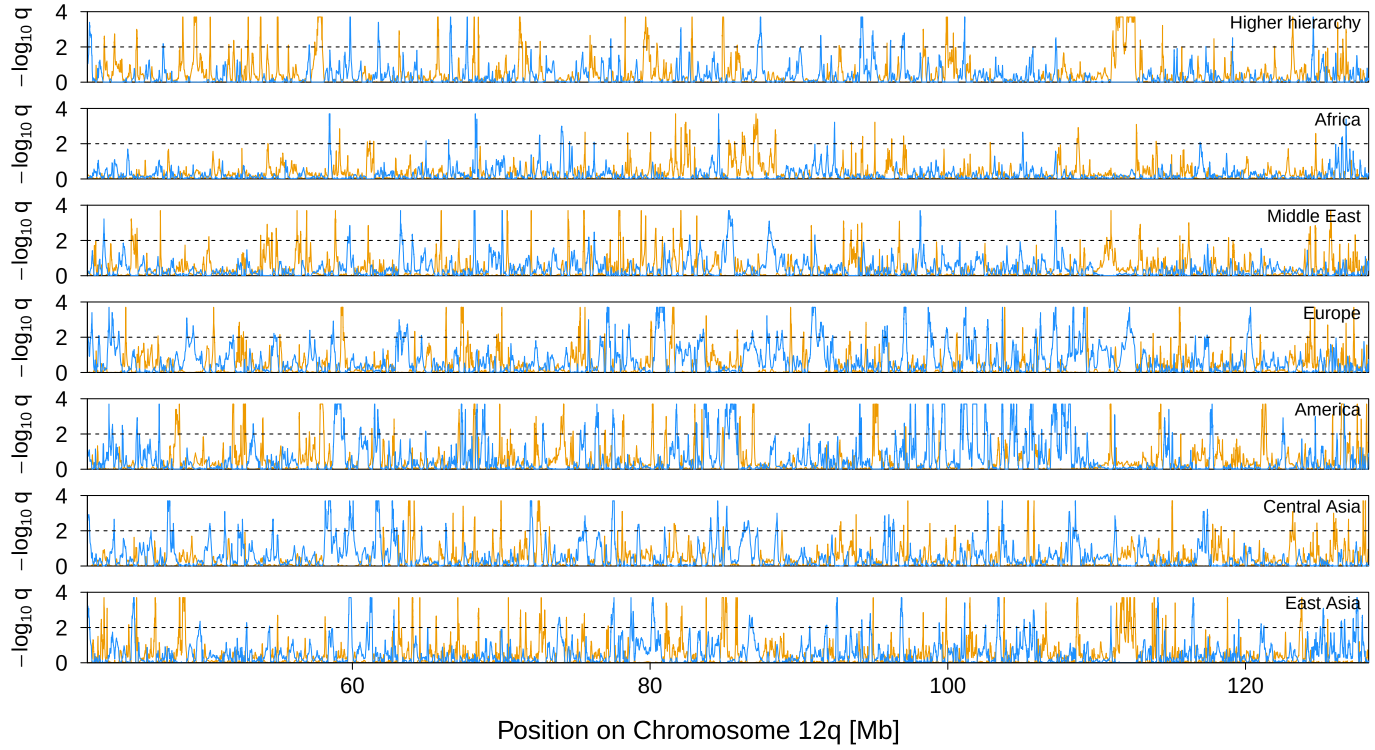


**Figure 27** Signal of selection on Chromosome 12q. The orange and blue lines indicate the locus-specific FDR for divergent (orange) and balancing (blue) selection, respectively. The black dashed line shows the 1% FDR threshold.

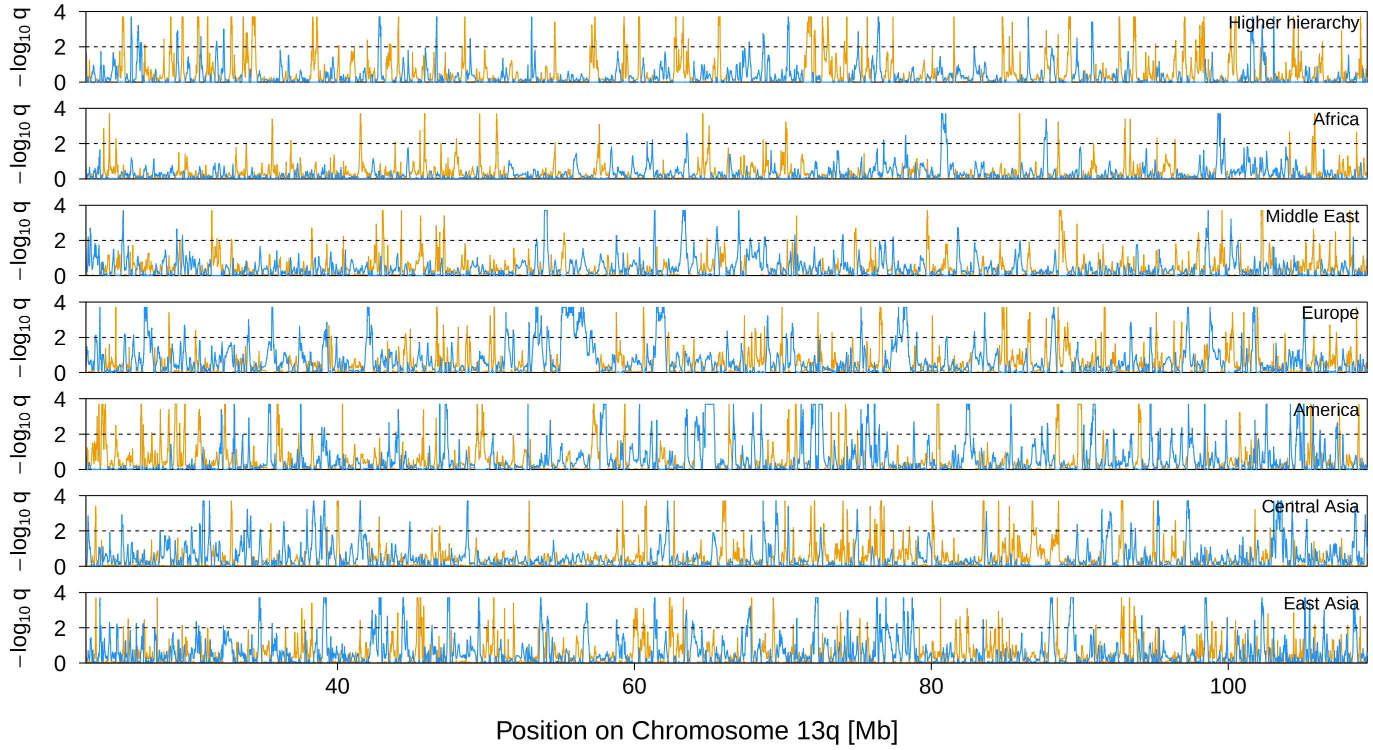


**Figure 28** Signal of selection on Chromosome 13q. The orange and blue lines indicate the locus-specific FDR for divergent (orange) and balancing (blue) selection, respectively. The black dashed line shows the 1% FDR threshold.

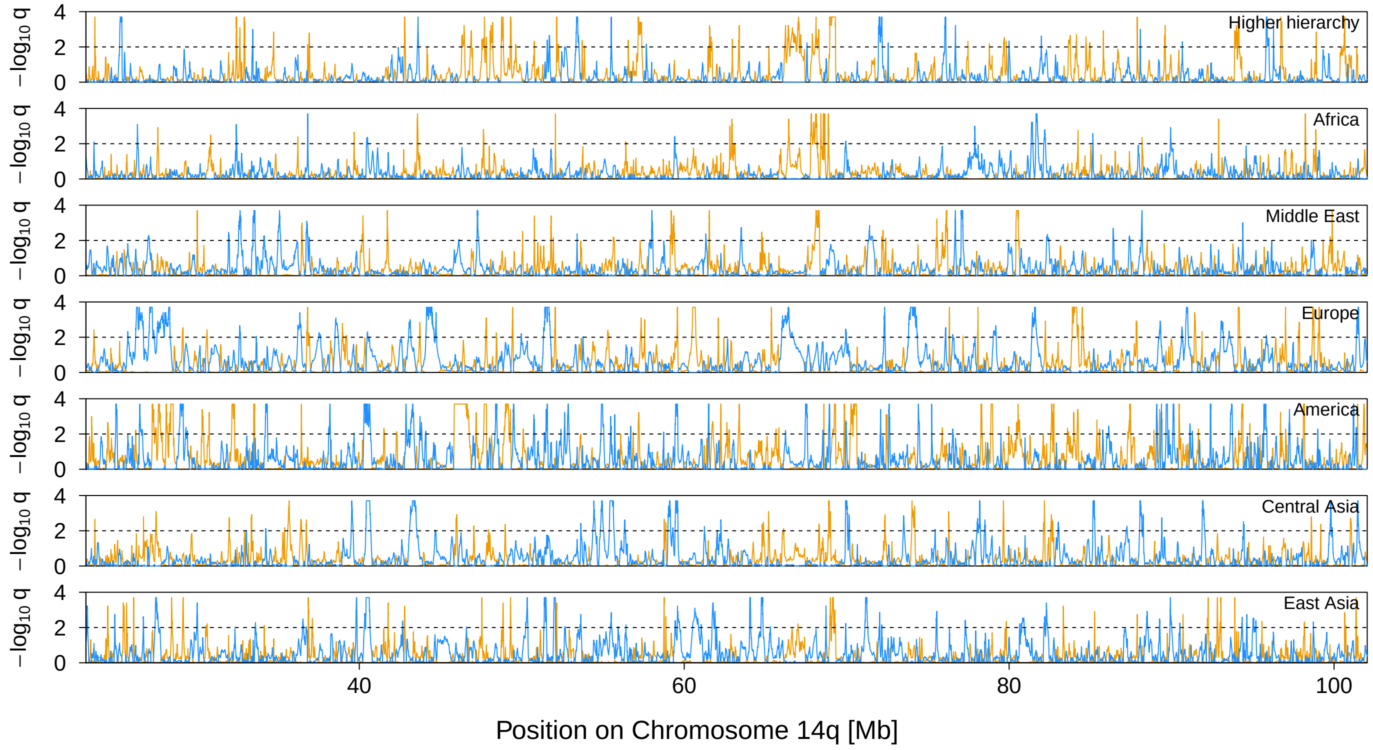


**Figure 29** Signal of selection on Chromosome 14q. The orange and blue lines indicate the locus-specific FDR for divergent (orange) and balancing (blue) selection, respectively. The black dashed line shows the 1% FDR threshold.

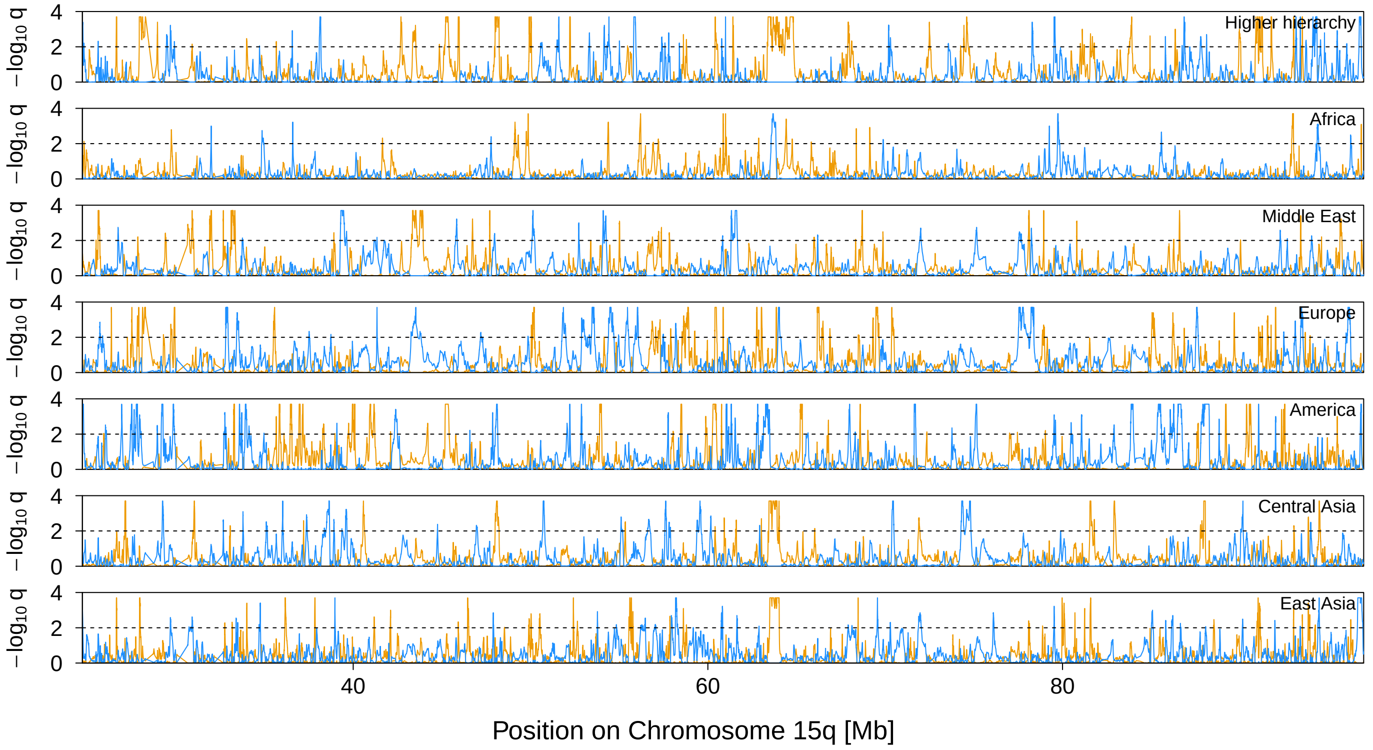


**Figure 30** Signal of selection on Chromosome 15q. The orange and blue lines indicate the locus-specific FDR for divergent (orange) and balancing (blue) selection, respectively. The black dashed line shows the 1% FDR threshold.

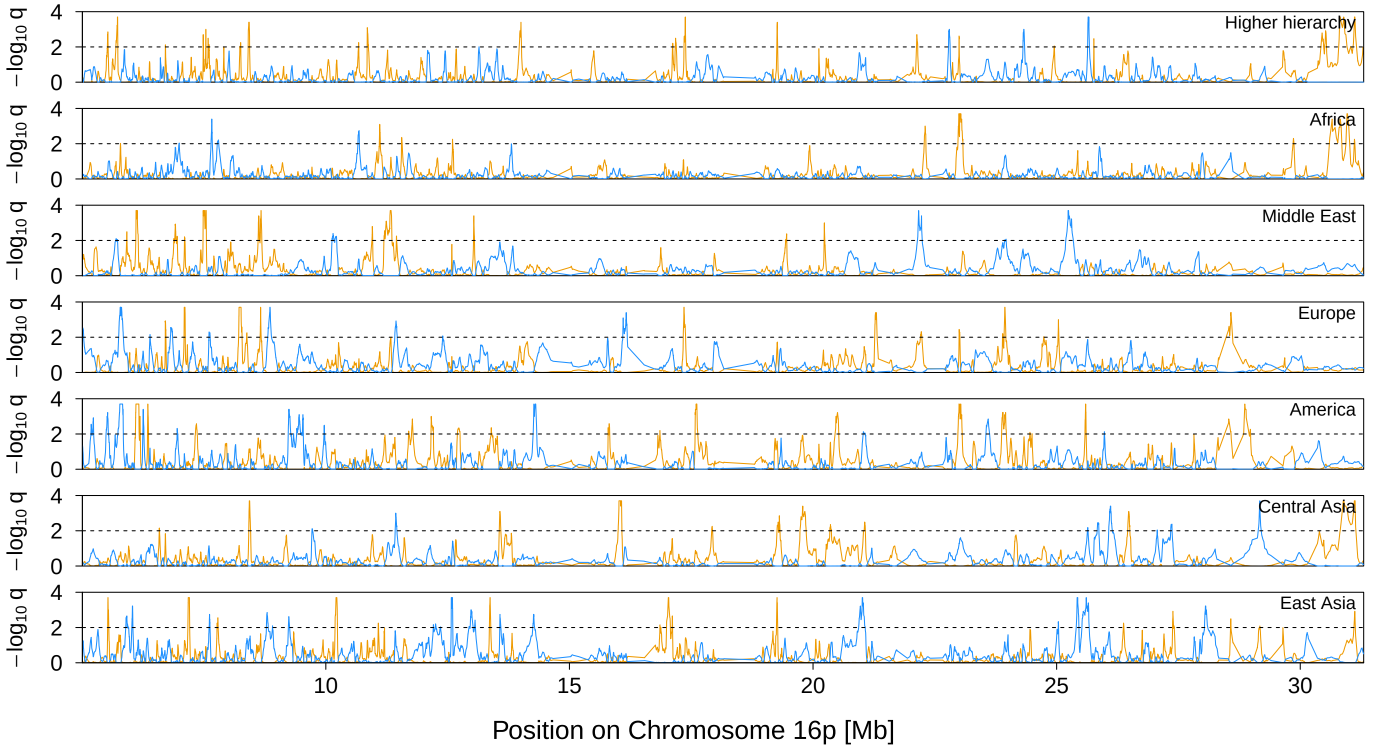


**Figure 31** Signal of selection on Chromosome 16p. The orange and blue lines indicate the locus-specific FDR for divergent (orange) and balancing (blue) selection, respectively. The black dashed line shows the 1% FDR threshold.

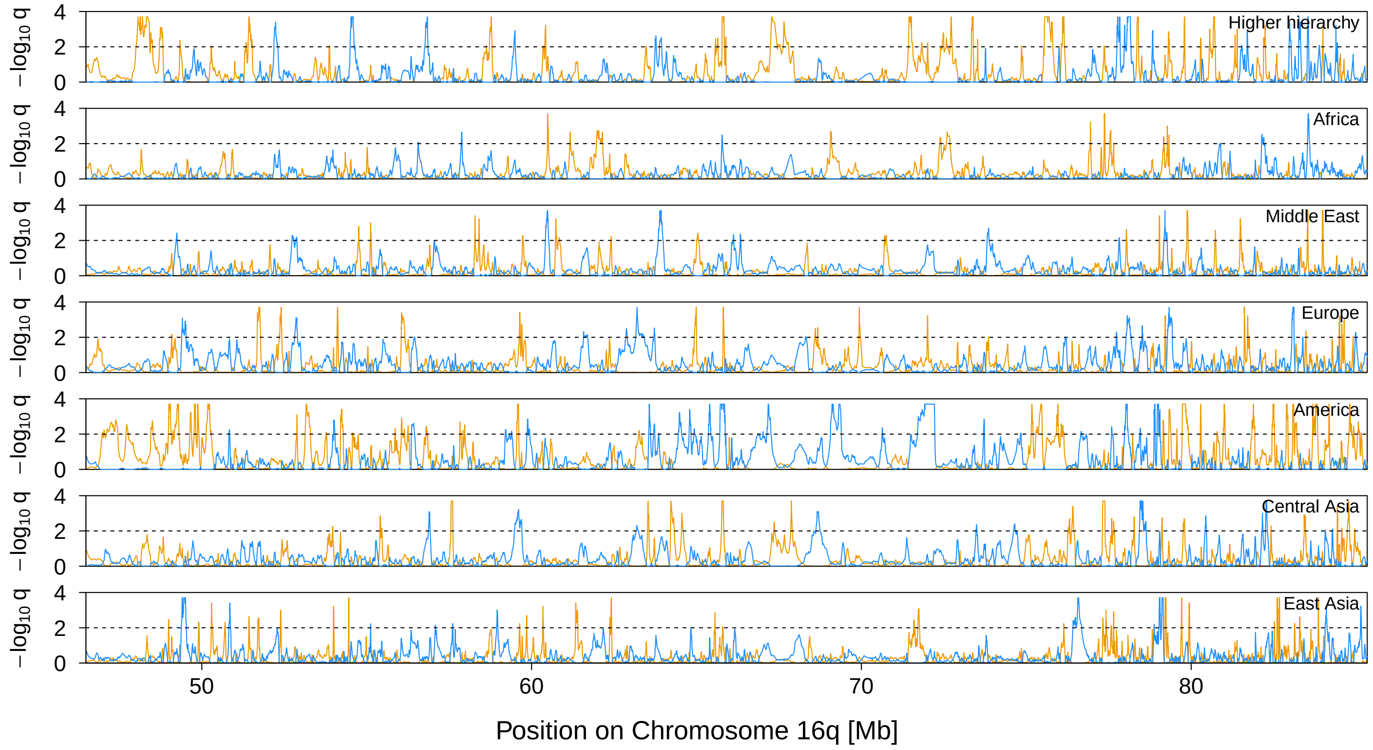


**Figure 32** Signal of selection on Chromosome 16q. The orange and blue lines indicate the locus-specific FDR for divergent (orange) and balancing (blue) selection, respectively. The black dashed line shows the 1% FDR threshold.

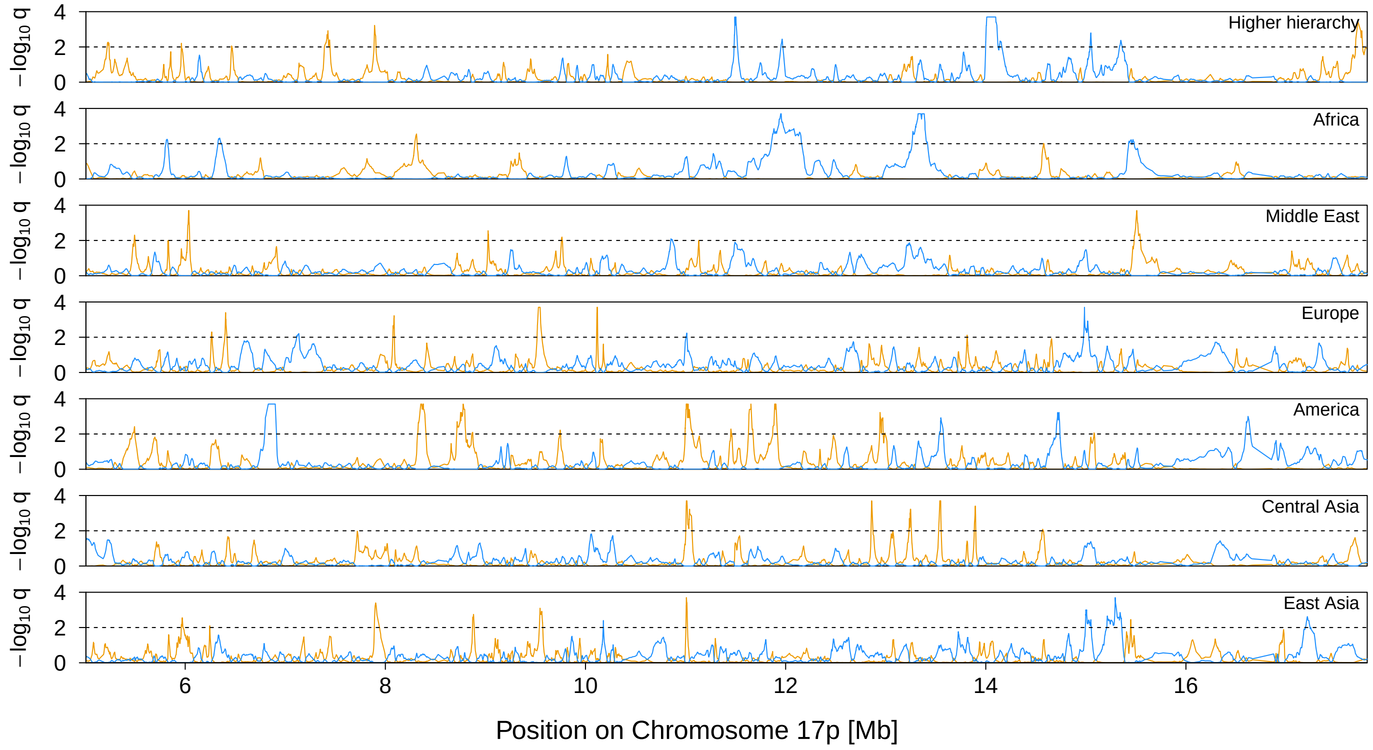


**Figure 33** Signal of selection on Chromosome 17p. The orange and blue lines indicate the locus-specific FDR for divergent (orange) and balancing (blue) selection, respectively. The black dashed line shows the 1% FDR threshold.

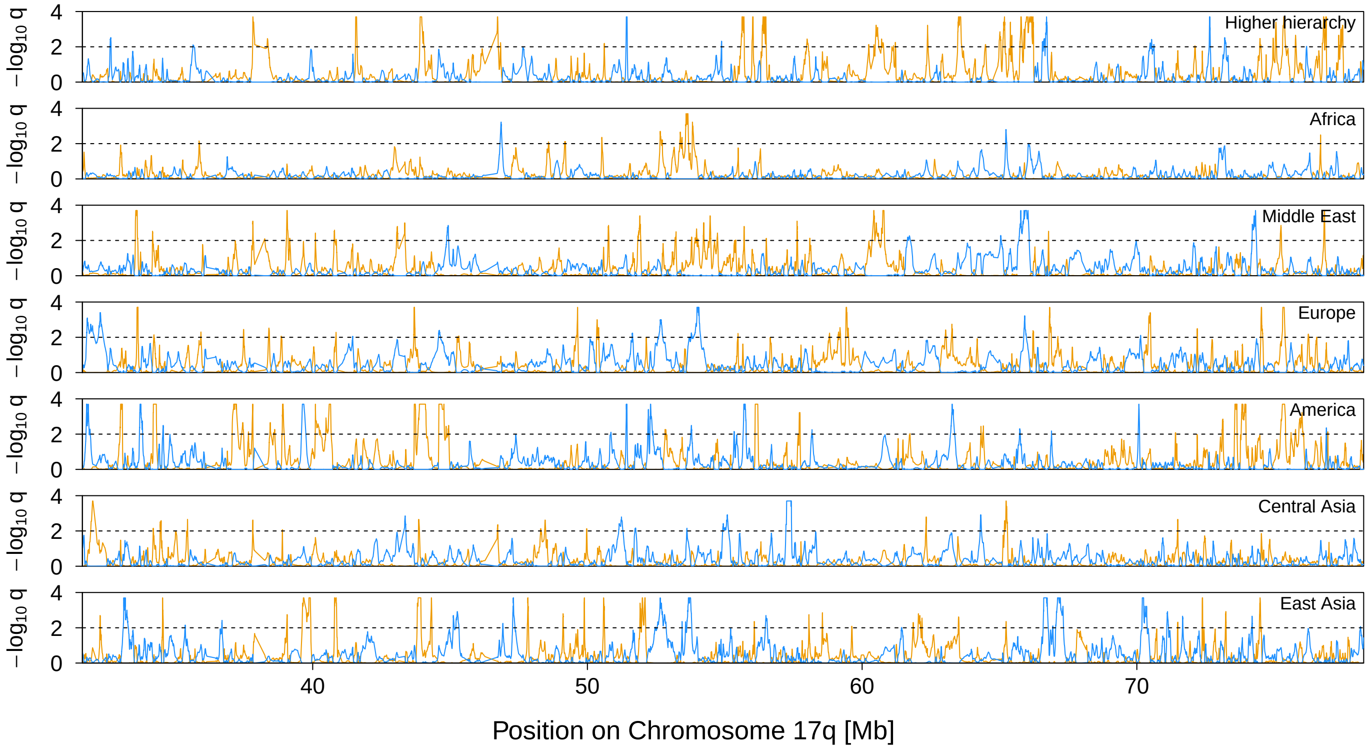


**Figure 34** Signal of selection on Chromosome 17q. The orange and blue lines indicate the locus-specific FDR for divergent (orange) and balancing (blue) selection, respectively. The black dashed line shows the 1% FDR threshold.

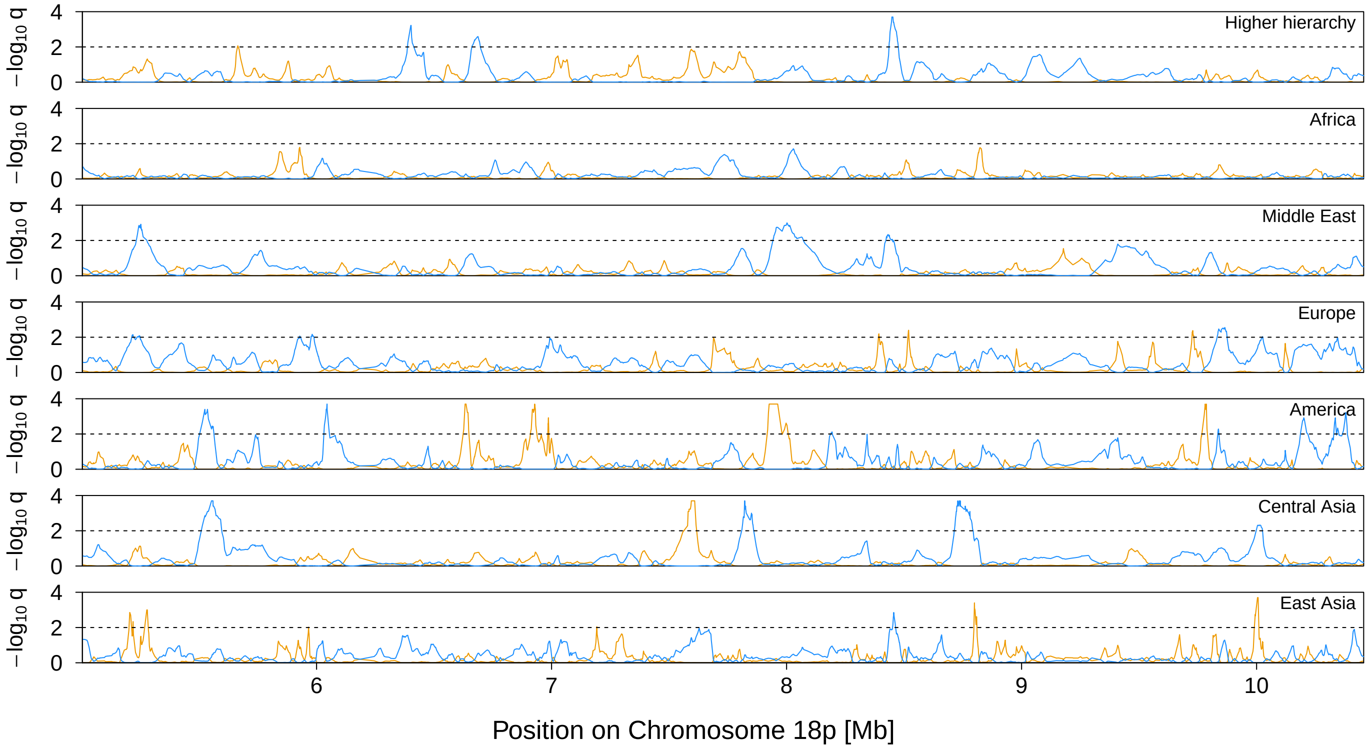


**Figure 35** Signal of selection on Chromosome 18p. The orange and blue lines indicate the locus-specific FDR for divergent (orange) and balancing (blue) selection, respectively. The black dashed line shows the 1% FDR threshold.

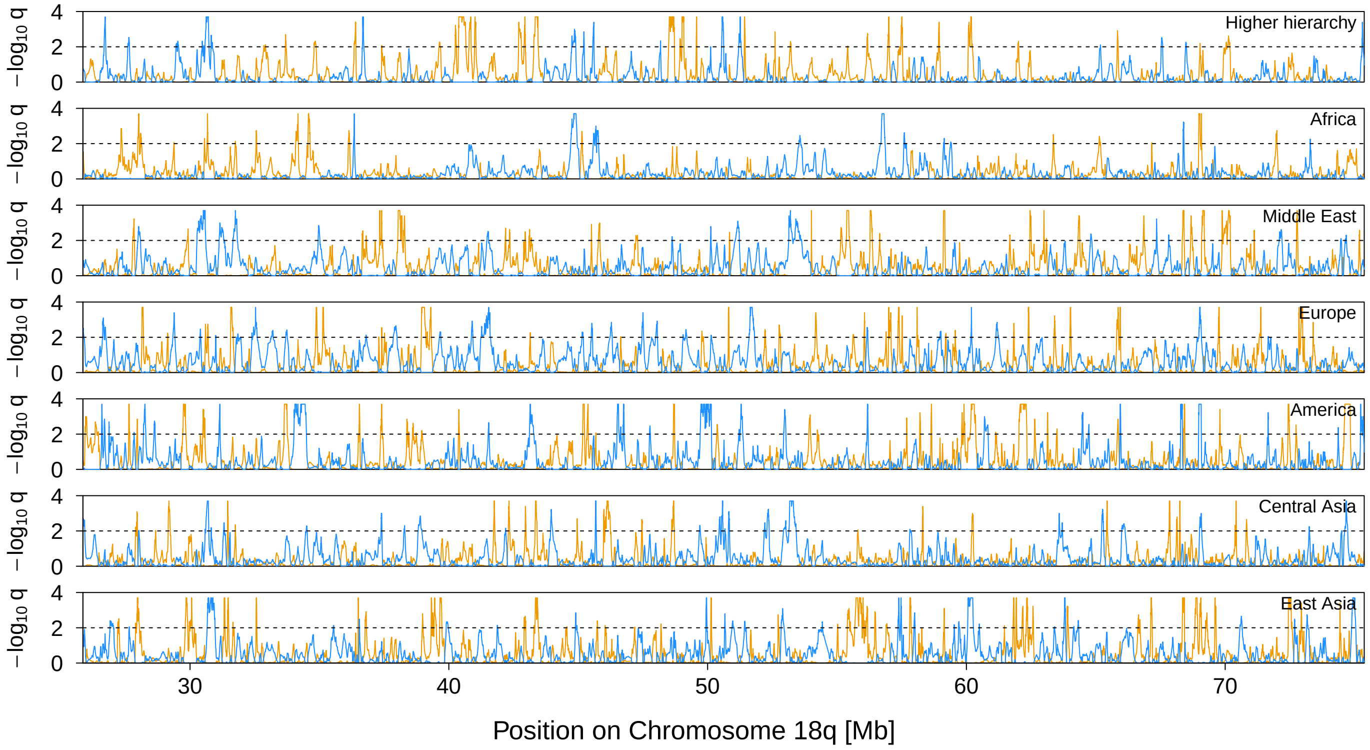


**Figure 36** Signal of selection on Chromosome 18q. The orange and blue lines indicate the locus-specific FDR for divergent (orange) and balancing (blue) selection, respectively. The black dashed line shows the 1% FDR threshold.

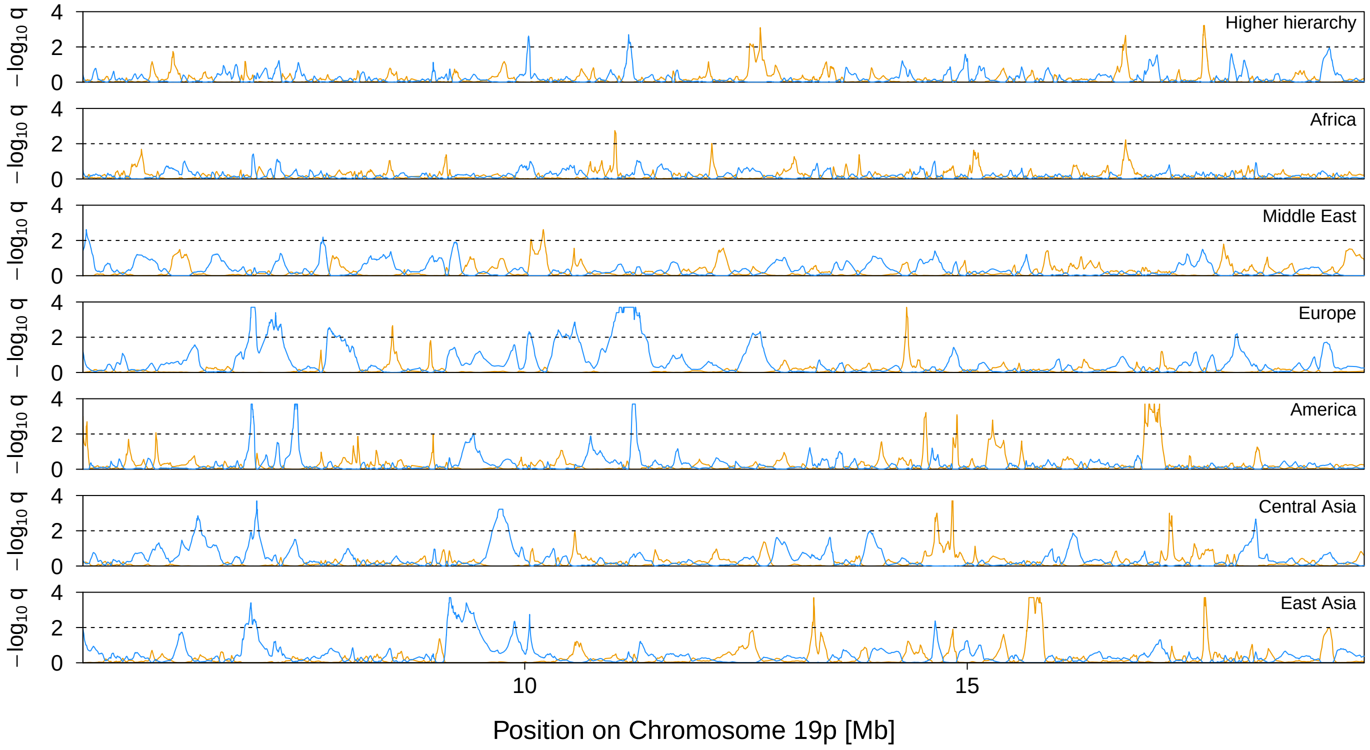


**Figure 37** Signal of selection on Chromosome 19p. The orange and blue lines indicate the locus-specific FDR for divergent (orange) and balancing (blue) selection, respectively. The black dashed line shows the 1% FDR threshold.

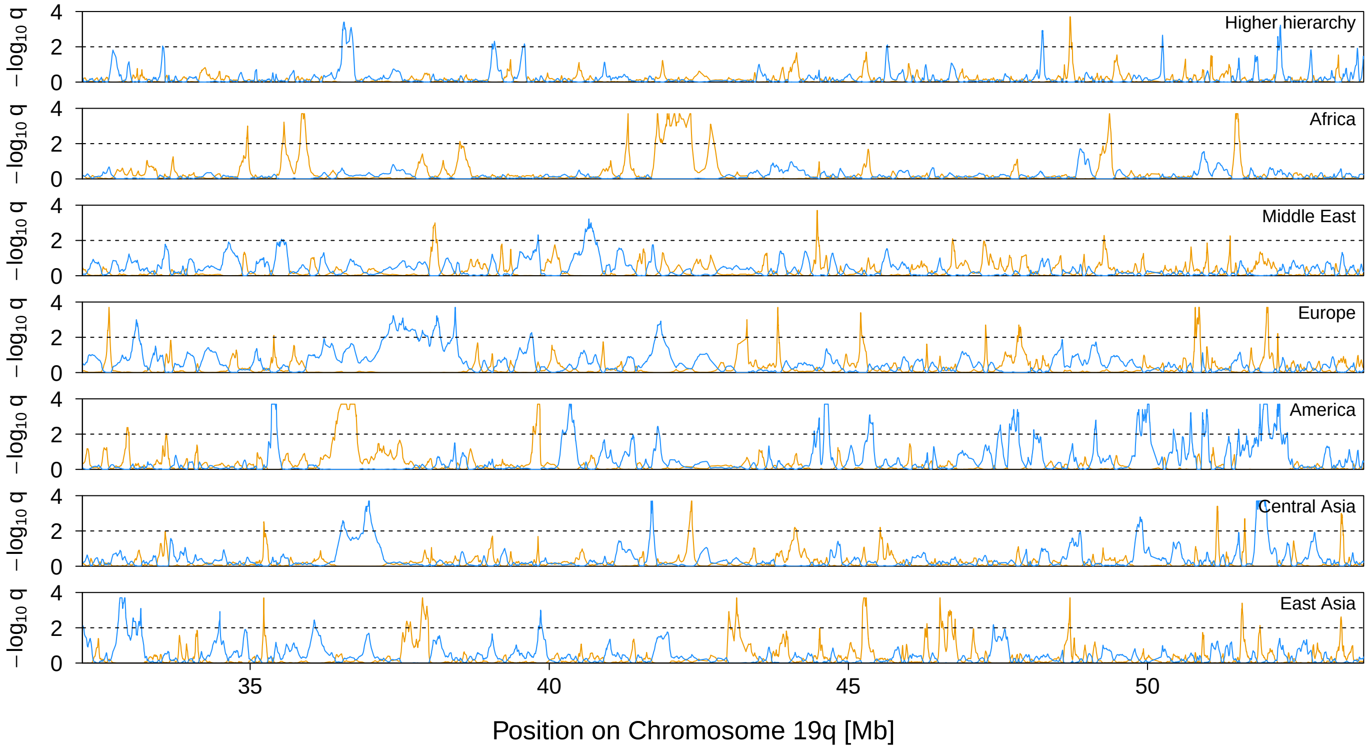


**Figure 38** Signal of selection on Chromosome 19q. The orange and blue lines indicate the locus-specific FDR for divergent (orange) and balancing (blue) selection, respectively. The black dashed line shows the 1% FDR threshold.

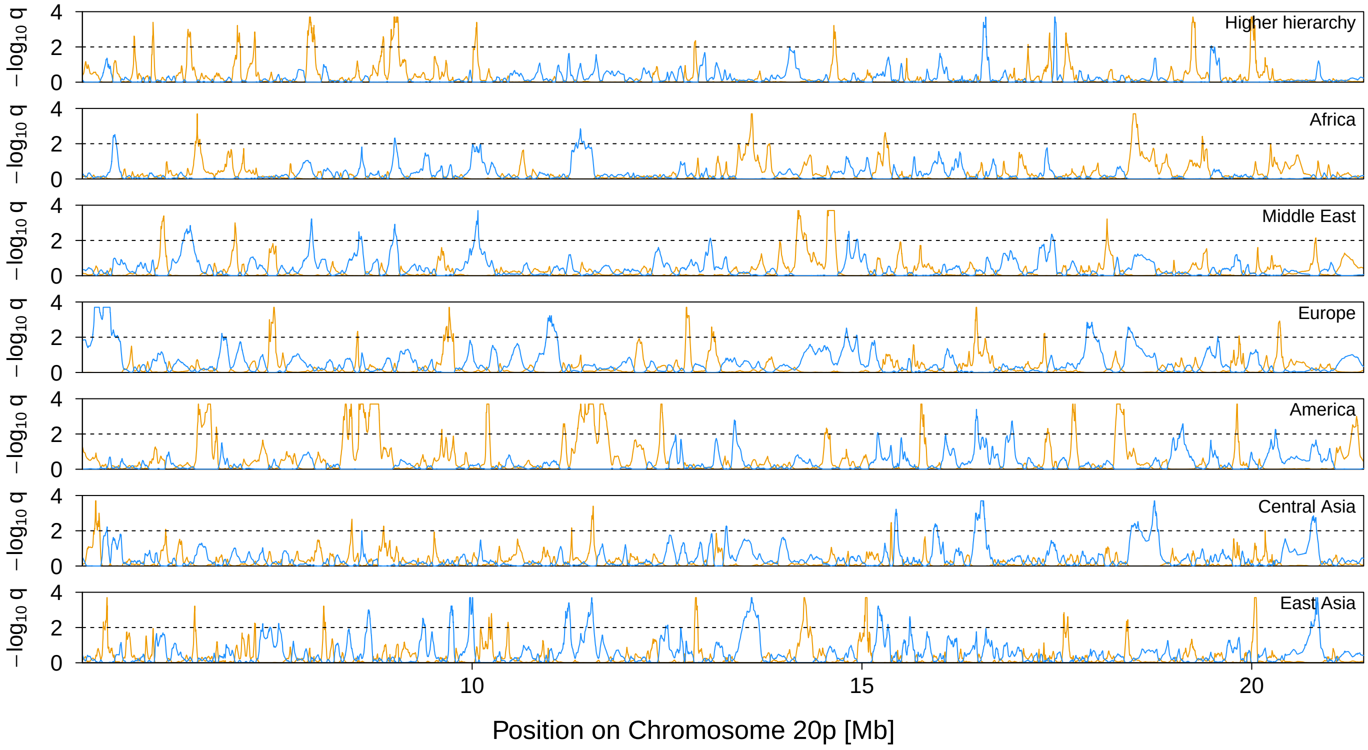


**Figure 39** Signal of selection on Chromosome 20p. The orange and blue lines indicate the locus-specific FDR for divergent (orange) and balancing (blue) selection, respectively. The black dashed line shows the 1% FDR threshold.

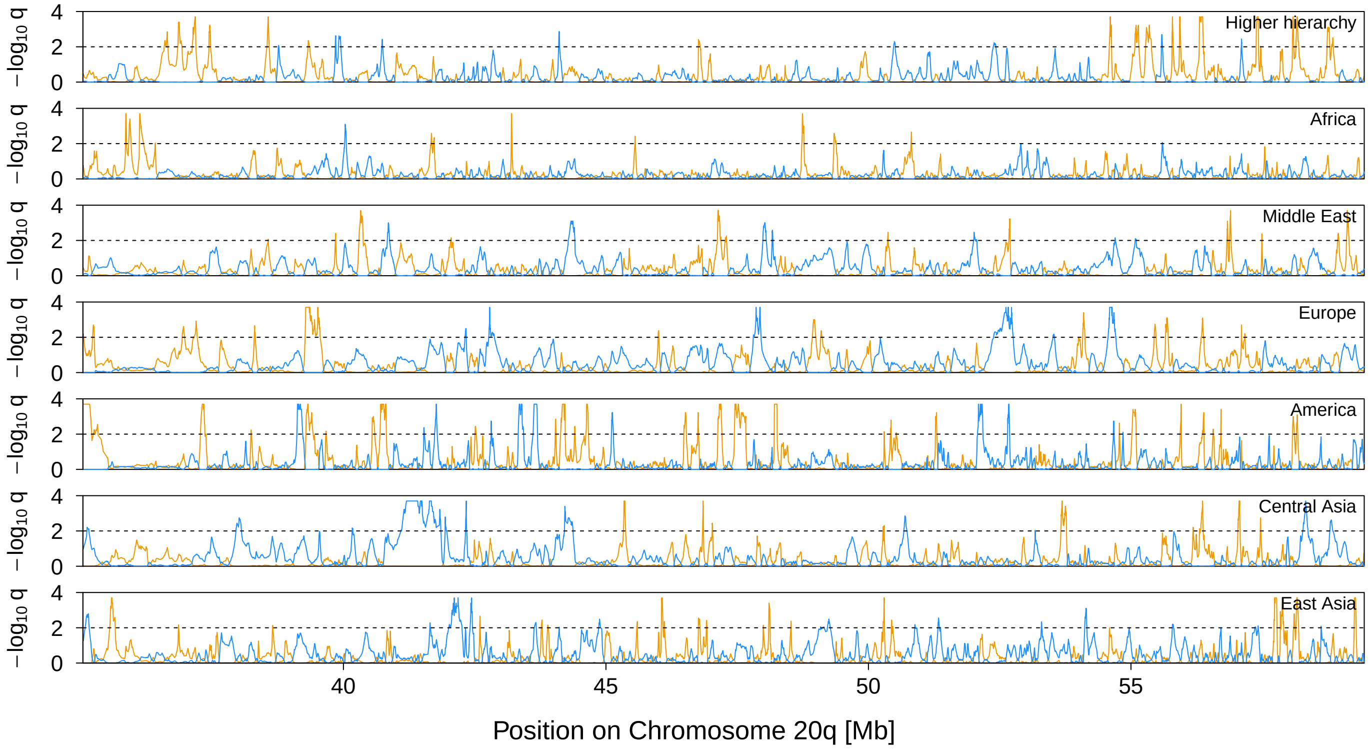


**Figure 40** Signal of selection on Chromosome 20q. The orange and blue lines indicate the locus-specific FDR for divergent (orange) and balancing (blue) selection, respectively. The black dashed line shows the 1% FDR threshold.

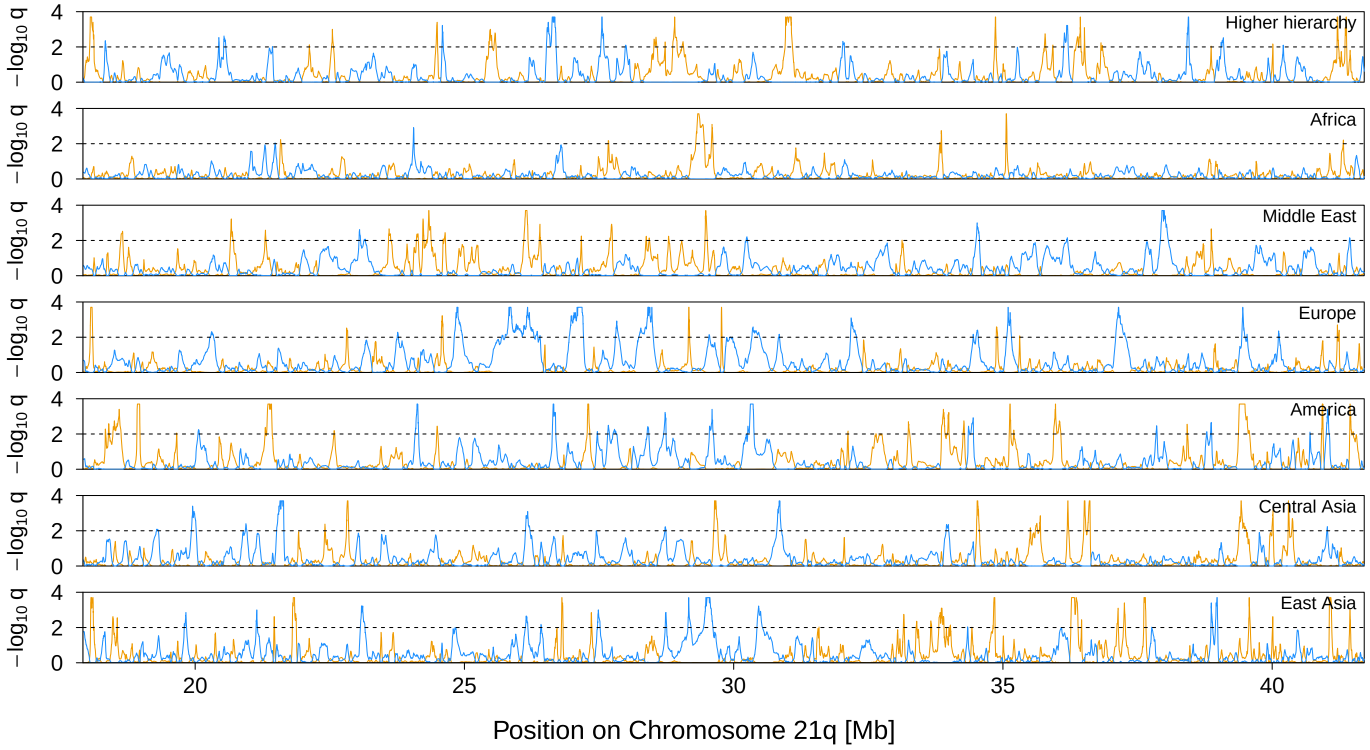


**Figure 41** Signal of selection on Chromosome 21q. The orange and blue lines indicate the locus-specific FDR for divergent (orange) and balancing (blue) selection, respectively. The black dashed line shows the 1% FDR threshold.

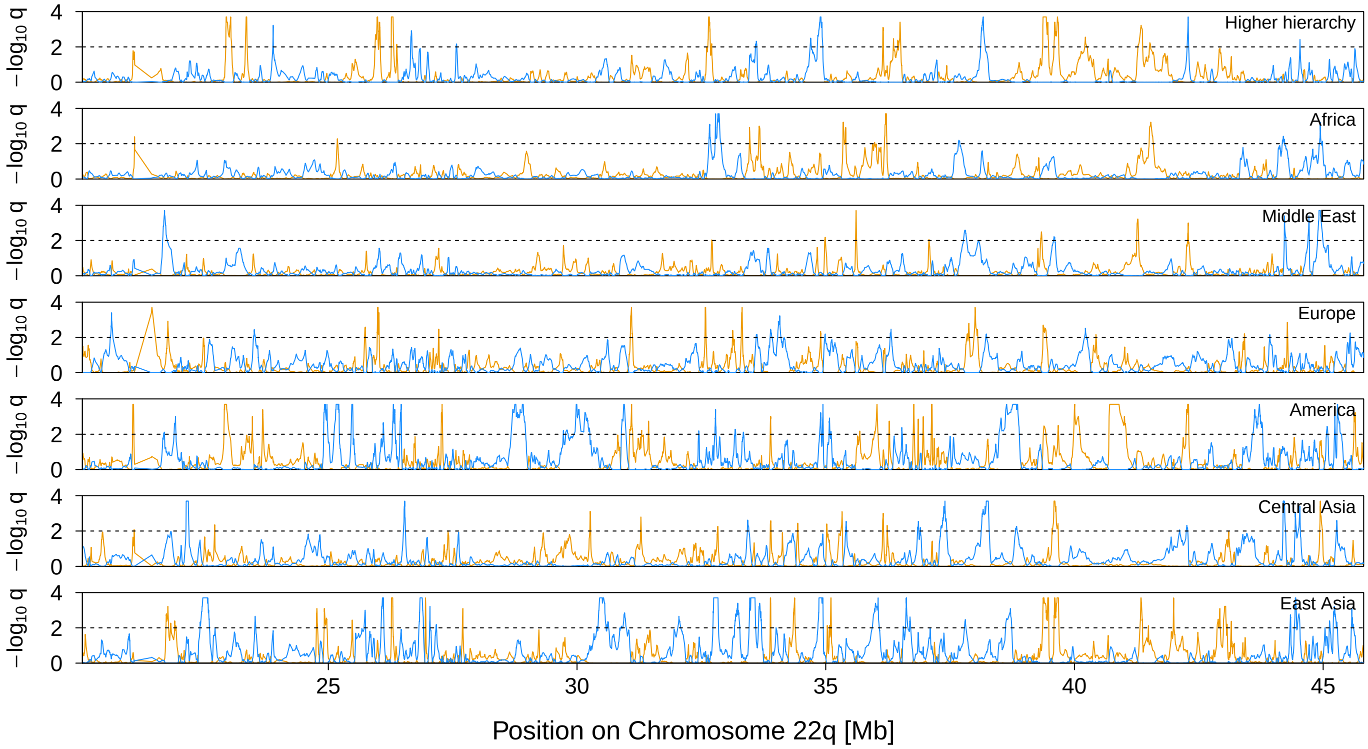


**Figure 42** Signal of selection on Chromosome 22q. The orange and blue lines indicate the locus-specific FDR for divergent (orange) and balancing (blue) selection, respectively. The black dashed line shows the 1% FDR threshold.
